# The SOX9 C-terminus encodes a transferable cBAF-recruitment module for enhancer remodeling and cell-state control

**DOI:** 10.64898/2026.11.09.765385

**Authors:** Akhouri K. Raghawan, Zhixin Li, Xiaolu Wang, Yingbo Huang, Paul Doan, Anoushka C. Gudipudi, Caroline Conway, Supakrit Kositbovornchai, Evan Hsiang, Pearl Y. P. Lie, James M. Cleary, Nilay S. Sethi

**Author notes:** Nilay Sethi, M.D., Ph.D. Department of Medical Oncology 450 Brookline Avenue, Dana 740 Dana-Farber Cancer Institute, Boston, MA 02215. University of California, Davis School of Medicine, Sacramento, CA, USA. These authors contributed equally (Akhouri K. Raghawan, Zhixin Li). These authors contributed equally (Xiaolu Wang, Yingbo Huang, Paul Doan).

## Abstract

Transcription factors select genomic targets, but how they recruit chromatin remodelers to convert DNA occupancy into cell-state-specific accessibility remains unclear. Here, we identify an intrinsically disordered SOX9 C-terminal module that recruits SMARCB1-containing canonical BAF (cBAF) to stem-like enhancers that open during *APC*-mutant adenoma formation. Acute, reversible SOX9 degradation displaces cBAF from ∼80% of co-bound enhancers, exceeding effects on pBAF and ncBAF, and collapses accessibility and stem-like identity; SOX9 re-expression restores these features. C-terminal truncation preserves DNA binding but disrupts cBAF engagement and chromatin opening. Enforced SMARCB1 docking to truncated SOX9 partially restores cBAF recruitment, accessibility, stem-like transcription, and organoid self-renewal, identifying remodeler recruitment as a limiting effector function. The isolated C-terminus supports higher-order cBAF assembly and, when fused to dCas9 as COMET, redirects endogenous cBAF to targeted loci to open chromatin. Disordered TF regions can therefore encode transferable instructions that deploy specific remodeling complexes to control cell state.

**HIGHLIGHTS:**

- The SOX9 C-terminal IDR recruits cBAF to adenoma-emergent enhancers
- SMARCB1 is required for SOX9-dependent enhancer accessibility and stem identity
- Forced SMARCB1 docking restores stem-like enhancer state and self-renewal to SOX9^ΔC^
- The isolated SOX9 module is sufficient to open chromatin via endogenous cBAF

## INTRODUCTION

Sequence-specific transcription factors (TFs) choose genomic targets, but DNA binding alone cannot explain how cell-state-specific chromatin landscapes are built. A central unresolved problem is how TFs selectively recruit broadly acting ATP-dependent chromatin remodelers to particular regulatory elements. Mammalian BRG1/BRM-associated factor (BAF, or SWI/SNF) complexes are central regulators of lineage-specific enhancer function^1–5^ and exist as biochemically distinct canonical BAF (cBAF), polybromo-associated BAF (pBAF), and non-canonical BAF (ncBAF) assemblies. Recent studies have begun to define remodeler-side determinants of TF engagement^6^, raising the reciprocal question of how TFs encode instructions for recruiting particular BAF assemblies to selected regulatory elements. Intrinsically disordered regions (IDRs), which frequently mediate multivalent interactions, transcriptional activation, and biomolecular condensation, are attractive candidates for such functions, but whether a TF IDR can encode a discrete and transferable chromatin-remodeler recruitment activity remains unclear.

Colorectal cancer (CRC) initiation provides a setting in which this question can be resolved during a defined cell-state transition. Somatic mutations that activate oncogenic pathways are widespread in histologically normal epithelia^7–11^, yet only a subset of altered clones persist and progress toward malignancy^12–14^, indicating that mutation must be coupled to nongenetic programs that stabilize abnormal cellular states. In the intestine, *APC* loss initiates neoplasia and drives constitutive WNT-β-catenin signaling^15–17^, expanding canonical stem cell activity^18^, but early lesions also acquire regenerative, fetal-like, and other aberrant stem-like programs together with remodeling of developmental enhancers^19–23^. SOX9 is tightly regulated in normal intestinal epithelium^24,25^ but becomes aberrantly deployed in gastrointestinal neoplasia, where it is required for tumor formation and suppresses differentiation through a stem-like transcriptional program^19,26,27^, including direct regulation of genes such as *PROM1*^26,28^. SOX9 has also been implicated in premalignant epithelial states across multiple tissues^19,29–31^, suggesting that its activity may represent a recurrent mechanism by which oncogenic signaling is converted into a durable, differentiation-resistant cell state.

A key unresolved feature of this biology is how SOX9 converts genomic occupancy into chromatin remodeling. We previously found that SOX9-dependent stem-like activity in CRC requires its intrinsically disordered C-terminal region^26^, suggesting that SOX9 function depends on activities beyond sequence-specific DNA recognition. This separation raises the possibility that the C-terminus recruits an effector complex required to remodel SOX9-bound enhancers. BAF complexes are particularly compelling candidates because they can be selectively deployed by TFs, yet individual BAF assemblies have distinct regulatory functions and genomic distributions^1–5^. Thus, three mechanistic questions remain unresolved: whether SOX9 preferentially engages a specific BAF assembly at stem-like enhancers, whether this interaction is encoded by its C-terminal IDR, and whether remodeler recruitment is sufficient to convert SOX9 DNA occupancy into enhancer accessibility and cell-state control.

Here, we identify the SOX9 C-terminus as a transferable cBAF-recruitment module that couples DNA binding to enhancer remodeling. SOX9 directs SMARCB1-containing cBAF to stem-like enhancers that become accessible during APC-mutant adenoma formation, and SMARCB1/cBAF is required for SOX9-dependent enhancer accessibility and stem-like identity. Acute, reversible degradation of endogenous SOX9 displaces cBAF from these elements and collapses their accessibility. C-terminal truncation uncouples SOX9 occupancy from chromatin opening, whereas enforced SMARCB1 docking partially restores enhancer accessibility, stem-like transcription, and organoid self-renewal. Finally, the isolated C-terminus, fused to dCas9 as COMET, recruits endogenous cBAF and opens targeted chromatin. Together, these findings define an IDR-encoded mechanism by which a lineage TF selectively deploys an endogenous chromatin remodeler to control enhancer state and cell identity.

## RESULTS

### SOX9 associates with BAF cofactors and co-occupies chromatin in CRC

The SOX9 C-terminus is required to suppress differentiation and sustain stem-like programs ^26^. To investigate how this region mediates SOX9 function, we performed rapid immunoprecipitation mass spectrometry of endogenous proteins (RIME), a modified ChIP-MS protocol ^32^, in HT-115 CRC cells expressing V5-tagged SOX9, a C-terminal deletion variant (SOX9^ΔC^), or GFP (Figure 1A). Mass spectrometry of chromatin-bound SOX9 complexes identified 37 proteins whose association depended on the SOX9 C-terminus (Figure 1A), along with an additional 110 proteins enriched with full-length SOX9 relative to SOX9^ΔC^ (Figure S1A and Table S1). Protein interactome analysis revealed enrichment for BRG1/BRM-associated factors (BAF, also known as SWI/SNF) complex components (*P* = 9.2 x 10^-12^) and spliceosome machinery (*P* = 2.4 x 10^-11^) (Figure 1B and Figure S1A-B). To prioritize candidates from the RIME dataset, we examined gene co-expression across TCGA CRC tumors. Genes encoding BAF components displayed stronger coordinated upregulation with SOX9 than other pathway-associated groups represented in the RIME dataset (Figure 1B and Figure S1C-D). Together with our prior finding that Sox9 regulates chromatin accessibility during CRC initiation ^19,26^, these observations nominated BAF cofactors as candidate effectors linking the SOX9 C-terminus to chromatin remodeling.

**Figure 1.**
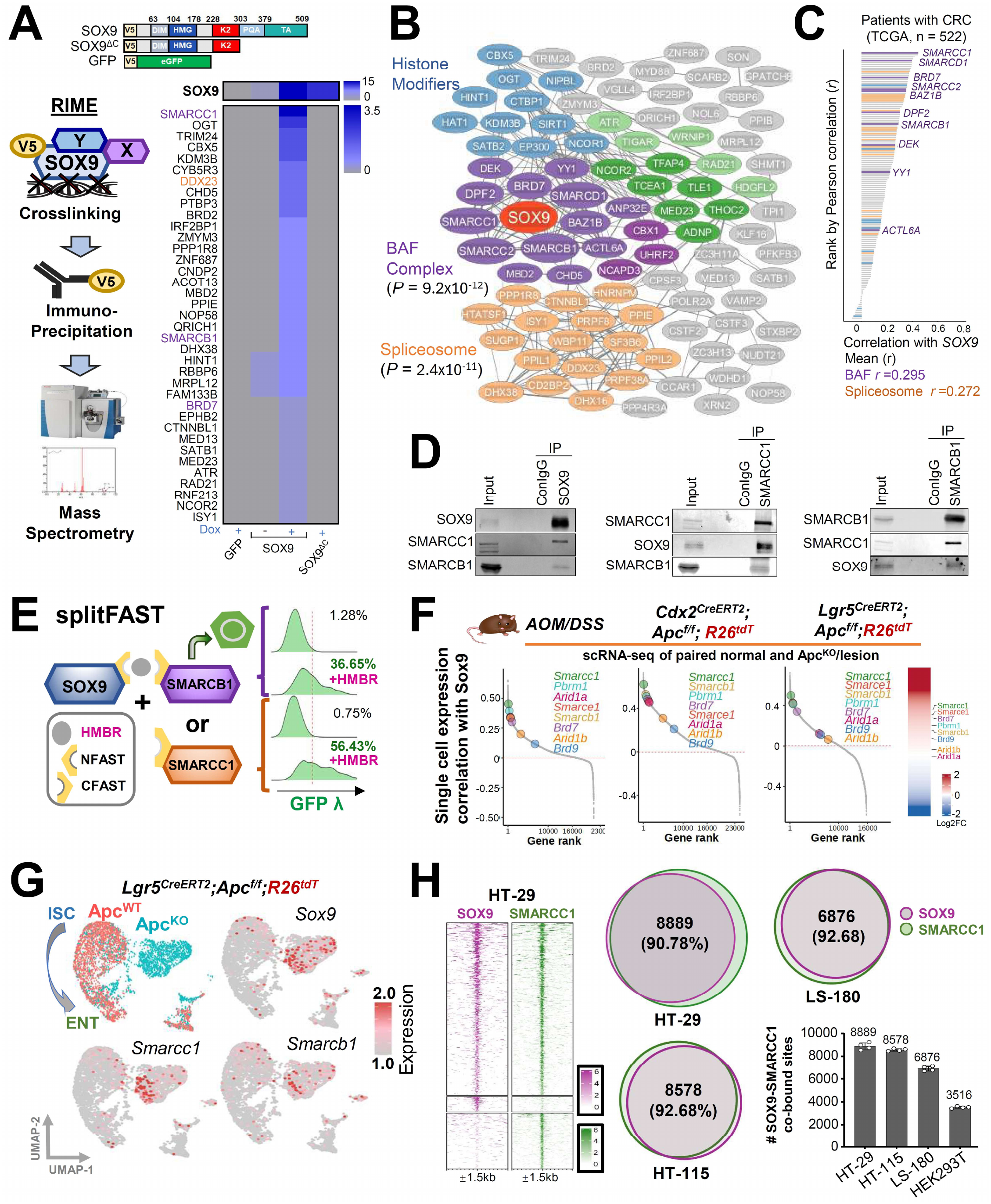
SOX9 associates with BAF cofactors and co-occupies chromatin. **(A)** Schematic of rapid immunoprecipitation mass spectrometry of endogenous proteins (RIME) (left). Heatmap of proteins detected by RIME in HT-115 CRC cells expressing V5-tagged GFP, SOX9 +/-dox induction, or SOX9^ΔC^ (right). **(B)** Protein interactome/pathway analysis of the SOX9 C-terminal interactome by GENETS, highlighting enrichment for BAF complex components and spliceosome factors. **(C)** Ranked correlation of SOX9 expression with genes encoding 147 proteins detected by RIME across TCGA CRC tumors (n =522); BAF components (purple) and spliceosome-associated factors (orange). **(D)** Immunoprecipitation of endogenous SOX9, SMARCC1, and SMARCB1 in HT-115 cells, showing reciprocal interaction among these proteins. **(E)** Split-FAST complementation assay in HEK293T cells co-expressing SOX9^NFAST^ with either SMARCC1^CFAST^ or SMARCB1^CFAST^ +/-HMBR followed by FACS (GFP wavelength) Single-cell RNA-seq analysis of paired normal and neoplastic colon epithelium from AOM/DSS lesions; and FACS-sorted tdT+ small intestines and colons from tamoxifen-induced Lgr5^CreERT2^;R26^tdT^ / Lgr5^CreERT2^;Apc^f/f^;R26^tdT^ (Bala et al., 2023) and Cdx2^CreERT2^;R26^tdT^ / Cdx2^CreERT2^;Apc^f/f^;R26^tdT^ mice, respectively **(G)** UMAPs of scRNA-seq data from Lgr5^CreERT2^;R26^tdT^ and Lgr5^CreERT2^;Apc^f/f^;R26^tdT^ mice (Bala et al., 2023) showing Apc^WT^ and Apc^KO^ cells as well as expression of Sox9, Smarcc1, and Smarcb1 **(H)** CUT&RUN heatmaps showing SOX9 and SMARCC1 occupancy in HT-29 CRC cells (left); Venn-diagram and bar graphs showing overlap of SOX9, SMARCC1, and SMARCB1 bound regions.

To validate the SOX9-BAF association, we performed endogenous immunoprecipitation (IP) of SOX9, SMARCC1, and SMARCB1 in CRC cell lines and confirmed reciprocal physical interaction among these proteins (Figure 1D and Figure S1E). Similar interactions were detected in adenoma organoids derived from Cdx2^Cre^;Apc^cKO^ mice (Figure S1F), supporting the relevance of this association in premalignant and transformed colon epithelial contexts. We further examined these interactions in living cells using split-FAST (fluorescence-activating and absorption-shifting tag) complementation^33^. Co-expression of C-terminus FAST (CFAST)-tagged SOX9 with either N-terminus FAST (NFAST)-tagged SMARCC1 or SMARCB1 in HEK293T cells produced robust fluorescence only in the presence of HMBR (Figure 1E and Figure S1G).

We next asked whether the SOX9-BAF association was reflected in CRC initiation *in vivo*. We generated and analyzed single-cell RNA sequencing (scRNA-seq) profiles of flow-sorted tdT+ normal colon epithelium and adenoma cells from two genetic mouse models of Apc loss (Lgr5^Cre^;R26R^td^ control and Lgr5^Cre^;Apc^cKO^;R26R^td^ mice^19,26^; Cdx2^Cre^;R26R^td^ control and Cdx2^Cre^;Apc^cKO^;R26R^td^ mice). Sox9 was induced together with multiple BAF components following Apc loss, and a similar pattern was observed in AOM/DSS carcinogen-induced colonic lesions (Figure 1F), suggesting that coordinated induction of SOX9 and BAF components is a shared feature across distinct CRC initiation models. Within the Lgr5^Cre^;Apc^cKO^;R26R^tdT^ model, co-expression of Sox9, Smarcc1, and Smarcb1 was especially strong in aberrant stem-like Apc^KO^ cells (Figure 1G and Figure S1H). Spatial transcriptomic analysis of human CRC further supported *in situ* co-upregulation of SOX9 and BAF components (Figure S1I).

To determine whether this biochemical partnership extends to chromatin, we mapped SOX9 and SMARCC1 occupancy using MNase-based CUT&RUN in three CRC cell lines and HEK293T controls. In CRC cells, SOX9 and SMARCC1 showed extensive chromatin co-occupancy, with 7,130– 8,848 co-bound regions across cell lines, corresponding to 92.8-96.5% of SOX9-bound loci (Figure 1H). The number of SOX9-SMARCC1 co-bound sites was substantially lower in HEK293T cells, supporting context-dependent enrichment of this chromatin partnership in CRC (Figure 1H and Table S2). Similarly, 82.4% of Sox9-bound sites were co-occupied by Smarcc1 in Cdx2^Cre^;Apc^KO^ organoids (Figure S1J). Thus, SOX9-associated BAF cofactors are induced in aberrant stem-like states during CRC initiation and converge with SOX9 at chromatin sites in CRC models.

### SOX9 preferentially engages cBAF at stem-like enhancers opened during adenoma formation

BAF subunits assemble into at least three biochemically and functionally distinct complexes: canonical BAF (cBAF), polybromo-associated BAF (pBAF), and non-canonical BAF (ncBAF)^2,3^ (Figure 2A). To determine which assemblies associate with SOX9, we performed endogenous SOX9 IP and chromatin profiling of assembly-specific subunits. SOX9 IP in HT-29 cells recovered cBAF components DPF2 and ARID1A, pBAF components PBRM1 and BRD7, and ncBAF component BRD9, indicating that SOX9 can associate with all three BAF assemblies (Figure S2A).

**Figure 2.**
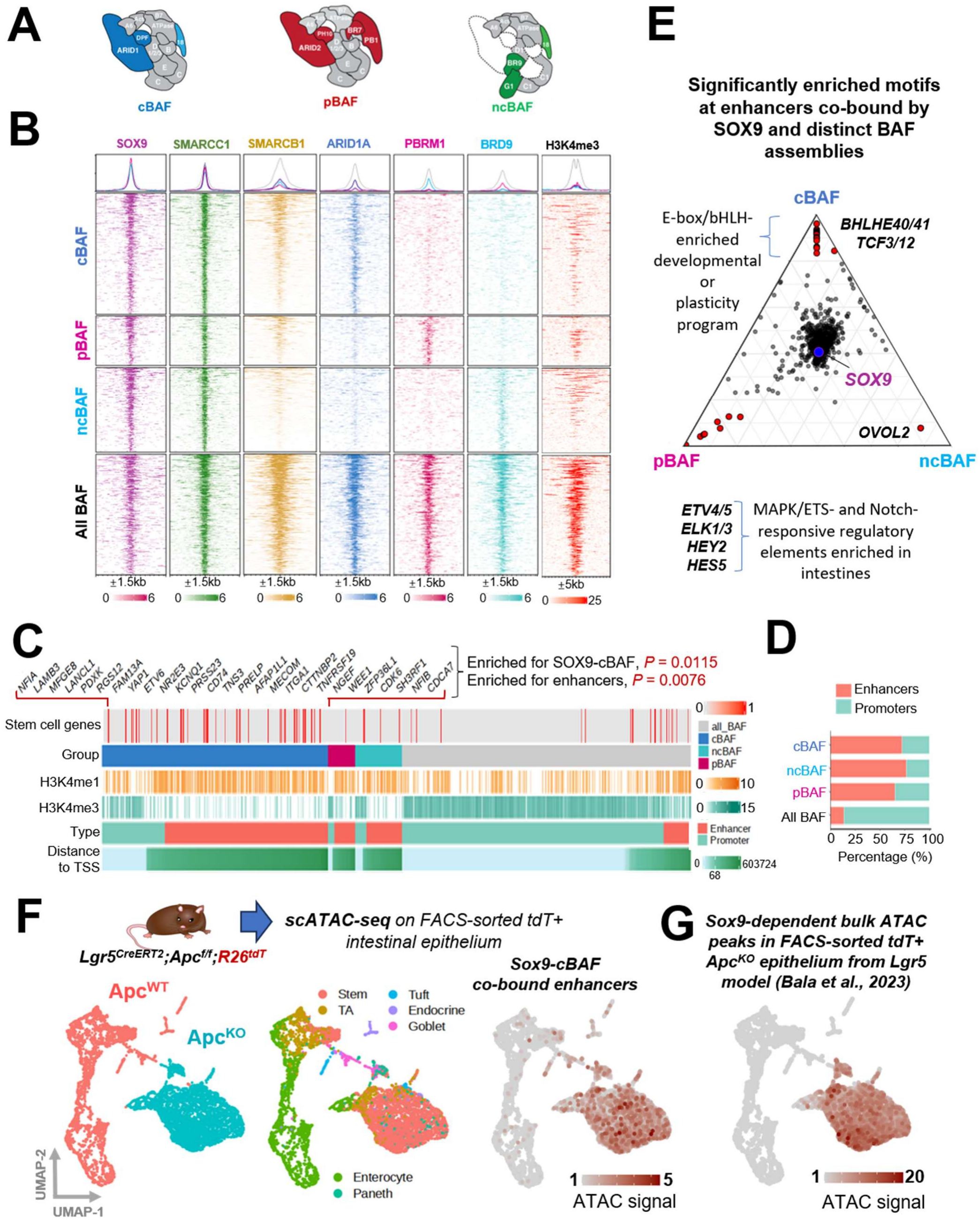
SOX9 preferentially engages cBAF at stem-like enhancers that open during adenoma formation. **(A)** Schematic of canonical BAF (cBAF), polybromo-associated BAF (pBAF), and non-canonical BAF (ncBAF) assemblies. **(B)** CUT&RUN heatmaps showing SOX9, SMARCC1, SMARCB1, ARID1A, PBRM1, BRD9, and H3K4me3 occupancy across cBAF-, pBAF-, ncBAF-, and all-BAF-associated genomic regions. **(C)** Annotation tracks showing gene set association, distance to transcription start site, histone modification state, genomic element type, and BAF assembly class for SOX9-BAF co-bound loci; associated stem-like genes are listed in Table S2. **(D)** Bar graph shows enhancers vs promoters distribution of SOX9-BAF co-bound genomic regions. **(E)** Motif enrichment analysis at enhancers co-bound by SOX9 and specific BAF complexes. **(F)** Single-cell ATAC-seq of FACS-sorted tdT+ intestinal epithelium from Lgr5^CreERT2^;R26^tdT^ and Lgr5^CreERT2^;Apc^f/f^;R26^tdT^ mice, showing UMAPs by genotype, cell type, SOX9-cBAF co-bound enhancer accessibility **(G)** UMAP of Sox9-dependent bulk ATAC peaks derived from FACS-sorted tdT+ intestinal epithelium from tamoxifen-induced Lgr5^CreERT2^;Apc^f/f^;R26^tdT^ and Lgr5^CreERT2^;Apc^f/f^;Sox9^f/+^;R26^tdT^ mice^19^.

We next asked how these assemblies partition across the SOX9-bound regulatory landscape. We mapped SOX9 occupancy together with ARID1A (cBAF), PBRM1 (pBAF), SMARCB1 (shared by cBAF and pBAF), and BRD9 (ncBAF). This analysis revealed that SOX9 co-occupies distinct genomic sites with individual BAF assemblies, alongside a subset of regions shared across all three complexes (Figure 2B). Assembly-specific SOX9-BAF sites were strongly enriched at enhancers, with 70% of SOX9-cBAF sites, 75% of SOX9-pBAF sites, and 65% of SOX9-ncBAF sites localizing to enhancer elements. In contrast, regions co-occupied by SOX9 and all three BAF assemblies were preferentially promoter-proximal (Figure 2B-D and Table S2). Histone-mark profiling across three CRC cell lines showed that assembly-specific SOX9-BAF sites carried active and poised features, whereas shared BAF sites were marked by promoter-associated H3K4me3 (Figure 2C and Figure S2B-S2C). In HEK293T cells, the corresponding CRC-defined enhancer loci showed markedly reduced SOX9 and SMARCB1 occupancy, more modest reduction in SMARCC1 signal, and reduced H3K27ac. By contrast, promoter-proximal shared BAF sites showed more similar chromatin features between HEK293T and CRC cells. Thus, the CRC-specific SOX9-BAF chromatin program is concentrated at enhancers, where SOX9-SMARCB1/cBAF engagement correlates with enhancer activation, whereas promoter-associated BAF occupancy is comparatively conserved across contexts.

We then asked whether these SOX9-BAF enhancer compartments were linked to distinct gene programs. Motif analysis suggested BAF subtype-specific separation of the SOX9 regulatory landscape. SOX9-cBAF sites were enriched for E-box/bHLH motifs associated with developmental plasticity programs, whereas SOX9-pBAF sites were enriched for MAPK/ETS- and Notch-responsive motifs characteristic of intestinal enhancer programs (Figure 2E). Consistent with this distinction, SOX9-cBAF loci were selectively enriched near genes upregulated in stem-like states (Figure 2C and Figure S2D, Fisher’s exact *P* = 0.0115) and were strongly associated with active and poised enhancers rather than promoters (odds ratio = 1.96, *P* = 0.0076), consistent with a role in promoting CRC stem-like transcriptional programs (Figure 2C and Figure S2B). Capture Hi-C data from HT-29 cells^34^ further linked this enhancer compartment to stem-like regulation by identifying physical contacts between SOX9-cBAF enhancers and promoters of stem cell-associated genes (Figure S2E and Table S3). In contrast, SOX9 sites associated with pBAF or ncBAF showed no comparable enrichment for stem cell-associated genes.

Finally, we asked which SOX9-BAF assemblies were associated with chromatin remodeling during CRC initiation. We analyzed single-cell ATAC-seq profiles from flow-sorted normal and adenoma tdT+ epithelial cells in the Lgr5^CreERT2^;Apc^cKO^ and projected SOX9-BAF chromatin classes onto the accessibility landscape (Figure 2F). Strikingly, sites co-bound by SOX9 and cBAF were most strongly enriched in open chromatin of aberrant Apc^KO^ stem-like cells (Figure 2F), whereas sites co-bound by SOX9 with pBAF, ncBAF, or all BAF assemblies did not show comparable adenoma-specific enrichment (Figure S2F). To determine whether this compartment was SOX9-dependent *in vivo*, we compared ATAC-seq profiles from FACS-isolated tdT+ adenoma cells with or without Sox9 deletion in the Lgr5 model^19^. Sox9-dependent accessible regions localized selectively within the Apc loss, stem-like cluster and included sites co-occupied by cBAF (Figure 2G). Together, these findings show that SOX9 engages BAF assemblies in a regulatory-context-specific manner, with SMARCB1-containing cBAF preferentially deployed at stem-like enhancers that become accessible during adenoma formation and remain active in CRC.

### SOX9 recruits and requires SMARCB1/cBAF to remodel stem-like enhancers

Having identified a SOX9-dependent, cBAF-bound enhancer compartment that emerges during tumor initiation, we next asked whether SOX9 instructs cBAF recruitment and stem-like chromatin remodeling. We engineered HT-29 CRC cells with biallelic endogenous *FKBP12^F36V^* degron tags at the SOX9 locus, enabling acute SOX9 degradation upon dTAG^v1^ treatment and subsequent SOX9 restoration after drug withdrawal (Figure S3A). SOX9 depletion promoted differentiation, as shown by increased KRT20 expression (Figure S3B), reduced proliferation (Figure S3C), and impaired colony formation (Figure 3A); these correlated with loss of stem cell signatures and induction of differentiation programs by bulk RNA-seq (Figure S3D, Tables S2 and S4). Time-resolved RNA-seq and ATAC-seq showed that four days of SOX9 depletion progressively dismantled the stem-like CRC state, whereas SOX9 restoration over the subsequent nine days re-established both transcriptional and chromatin features of this program (Figure 3B-C and Figure S3E). Single-nucleus RNA-seq further showed that SOX9 depletion rapidly shifted cells toward a differentiated state, with reduced CytoTrace2-inferred potency^35^ and near-complete loss of the multipotent compartment by day 3 (Figure 3D and Figure S3F). Thus, endogenous SOX9 is a reversible determinant of CRC cell-state plasticity, required to maintain stem-like identity and sufficient to reimpose it after differentiation.

**Figure 3.**
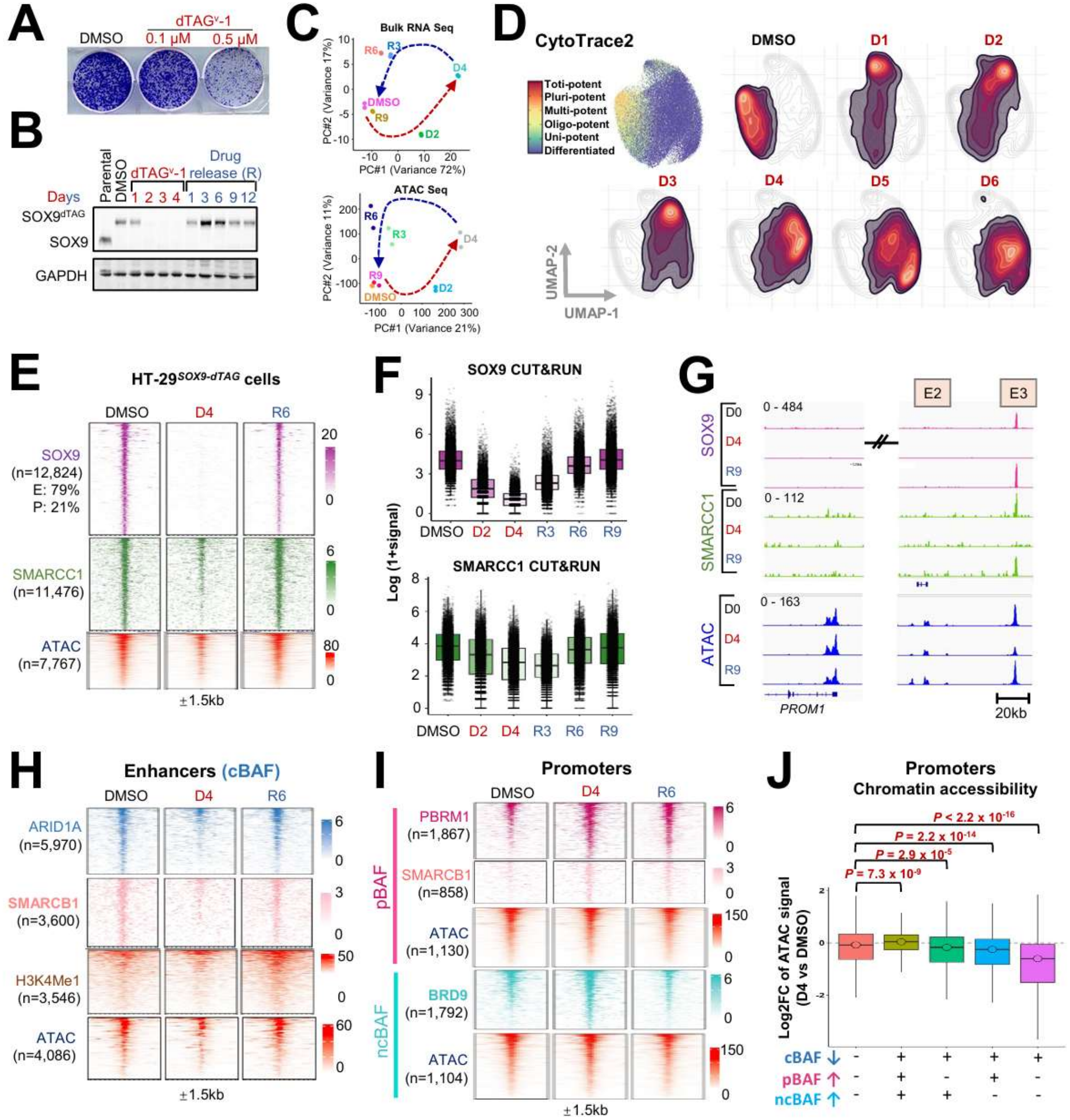
Endogenous SOX9 is required for cBAF occupancy and stem-like enhancer accessibility. **(A)** Colony formation assay in HT-29*^SOX9-dTAG^* treated with 0.1µM and 0.5µM of dTAG^v^-1 **(B)** Immunoblot of SOX9 and GAPDH (loading control) in parental and HT-29*^SOX9-dTAG^* cells at different timepoints of dTAG^v^-1 treatment and release **(C)** PCA plots of bulk RNA-seq data (top) and bulk ATAC-seq data (below) of HT-29*^SOX9-dTAG^* cells at different timepoints dTAG^v^-1 treatment (SOX9 degradation) and drug wash-off (SOX9 restoration) **(D)** UMAPs from single-nucleus RNA-seq data of HT-29*^SOX9-dTAG^* cells at different timepoints dTAG^v^-1 treatment (SOX9 degradation time course) with CytoTrace-based potency visualization. **(E)** CUT&RUN heatmaps for SOX9 and SMARCC1, and chromatin accessibility (ATAC-seq) across SOX9-regulated loci, under DMSO, SOX9-loss upon dTAG^v^-1 treatment for 4 days (D4) and SOX9 restoration after dTAG^v^-1 release for 6 days (R6); E = Enhancers and P= Promoters. **(F)** Box plots show CUT&RUN signals from SOX9 and SMARCC1 in HT-29*^SOX9-dTAG^* cells at different timepoints of dTAG^v^-1 treatment (D2 and D4) and release (R3, R6 and R9). P-values are listed in Table S5. **(G)** Genome browser views the *PROM1* locus, showing an intronic enhancer and two distal enhancers, SOX9 and SMARCC1 CUT&RUN signal, and ATAC-signal from HT-29*^SOX9-dTAG^* cells at different timepoints of dTAG^v^-1 treatment (D4) and release (R6). **(H)** Heatmaps showing ARID1A, SMARCB1 and H3K4Me1 CUT&RUN heatmaps and chromatin accessibility at SOX9-cBAF co-bound enhancer regions under DMSO, dTAG^v^-1 treatment (D4), and release (R6) conditions. **(I)** Heatmaps showing PBRM1, SMARCB1 (pBAF) and BRD9 (ncBAF) occupancy, and associated chromatin accessibility, across SOX9-pBAF/ncBAF co-bound regions under DMSO, dTAG^v^-1 treatment (D4), and release (R6) conditions. **(J)** Box plots of chromatin accessibility across promoter-associated loci stratified by cBAF loss and gain of pBAF/ncBAF assemblies.

We next used this reversible system to determine whether SOX9 directly controls BAF recruitment and enhancer accessibility. We mapped SOX9, core BAF subunits, assembly-specific BAF components, histone modifications, and chromatin accessibility during SOX9 depletion and restoration. Among stringent SOX9-SMARCC1 co-bound regions (n=12,824), SOX9 depletion reduced SMARCC1 occupancy at 90% of sites, with concomitant loss of chromatin accessibility and H3K4me1 at approximately 60% of these regions (Figure 3E-F and Figure S3G; Tables S5-S6). SOX9 restoration reinstated SMARCC1 occupancy, enhancer accessibility, and H3K4me1, including at a distal enhancer of the canonical stem cell gene *PROM1*, which we previously identified as a direct SOX9 target (Figure 3G) ^26^. These findings show that SOX9 is required to maintain, and sufficient to restore, BAF recruitment and accessibility at stem-like regulatory elements.

We then asked which BAF assembly was most dependent on SOX9. At enhancer elements, cBAF occupancy showed the strongest SOX9 dependence: SOX9 degradation reduced ARID1A binding at 83.6% of co-bound sites and SMARCB1 binding at 78.8% of co-bound sites (Figure 3H and Table S6). In contrast, enhancer occupancy by ncBAF and pBAF was less broadly affected, with reduced BRD9 and PBRM1 binding at 59% and 47.5% of co-bound sites, respectively. Loss of cBAF occupancy was accompanied by reduced chromatin accessibility and H3K4me1, all of which were restored upon SOX9 re-expression (Figure 3H and Table S6). Promoter-associated BAF occupancy showed a distinct pattern: loss of cBAF was partially buffered at some promoters by increased pBAF or ncBAF binding, which preserved promoter accessibility in proportion to the degree of alternative BAF recruitment (Figure 3I-J and Table S6). Thus, SOX9 most directly and reversibly controls cBAF-dependent chromatin remodeling at enhancer elements, whereas promoter-associated BAF occupancy is more plastic.

### SOX9 depends on cBAF, and particularly SMARCB1, to enforce a stem-like state

We tested whether cBAF is functionally required for SOX9-mediated cell-state control. We combined reversible SOX9 restoration with inducible suppression of individual BAF subunits and used KRT20 induction as a readout of failed differentiation suppression (Figure S4A). SOX9 depletion increased KRT20 expression, whereas SOX9 restoration returned KRT20 to baseline in control cells (Figure 4A). Suppression of most BAF subunits did not substantially impair this rescue. In contrast, loss of cBAF components, particularly SMARCB1 and ARID1A/ARID1B, compromised SOX9-mediated suppression of KRT20, whereas depletion of the pBAF subunit PBRM1 or the ncBAF subunit BRD9 had little effect (Figure 4A and Figure S4B). Consistent with this functional requirement, SMARCB1 suppression or combined ARID1A/ARID1B suppression impaired restoration of enhancer accessibility after SOX9 re-expression, with little effect at promoters co-bound by multiple BAF assemblies (Figure 4B). Together, these results show that SOX9 recruits cBAF to stem-like enhancers and requires intact cBAF, particularly SMARCB1, to restore enhancer remodeling and suppress differentiation.

**Figure 4.**
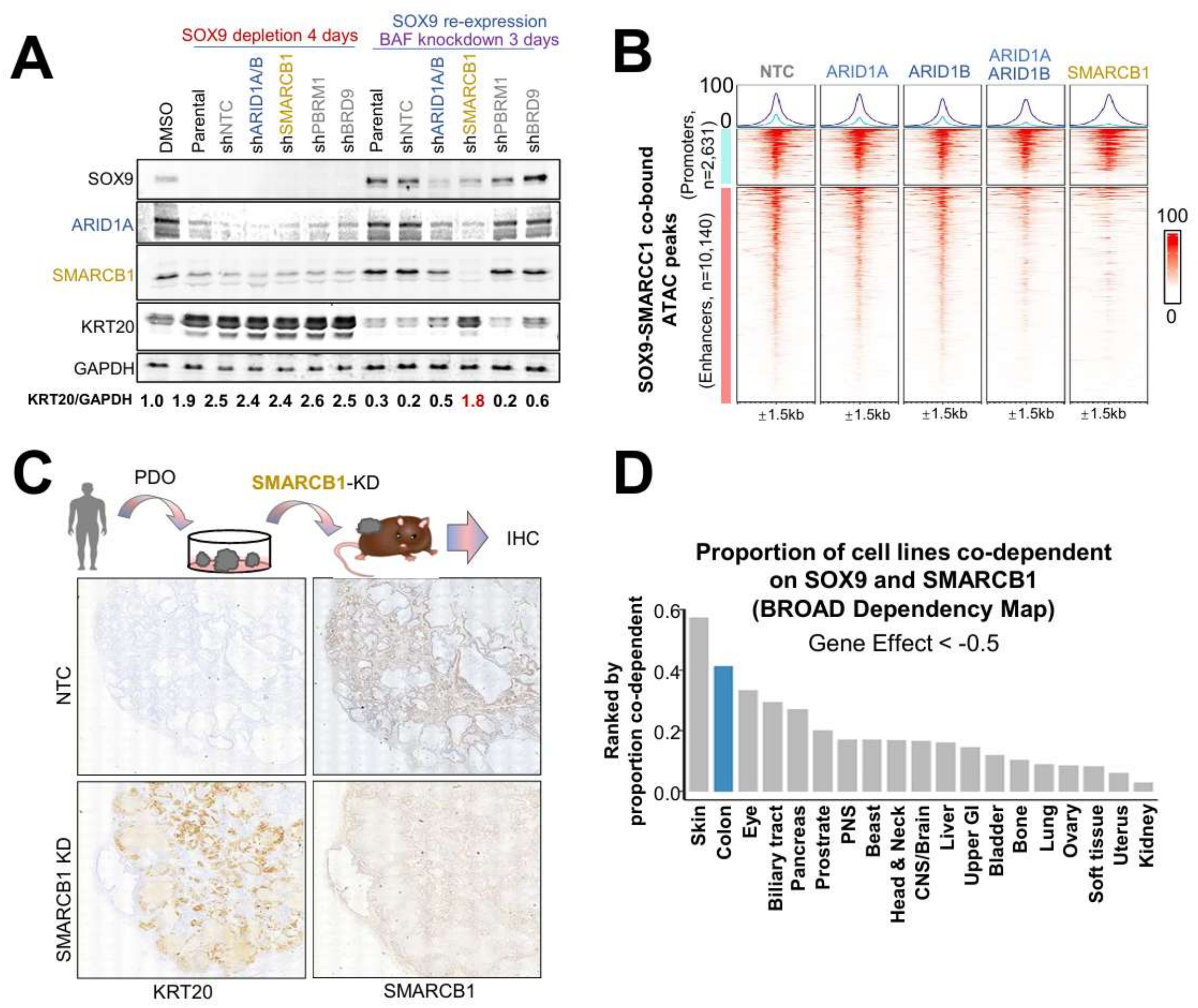
SOX9 depends on cBAF, and particularly SMARCB1, to enforce a stem-like state. **(A)** Immunoblot analysis of SOX9, ARID1A, SMARCB1, KRT20 (differentiation marker), and GAPDH in HT-29*^SOX9-dTAG^* cells following SOX9 depletion, BAF subunit knockdown, and SOX9 re-expression. **(B)** ATAC-seq heatmaps across SOX9-SMARCC1 co-bound regions under BAF subunit knockdown conditions. **(C)** Panel shows SMARCB1 and KRT20 (differentiation marker) levels in IHC of patient-derived xenografts upon SMARCb1-knockdown. **(D)** Bar graph shows proportion of cancer cell lines co-dependent on SOX9 and SMARCB1.

We next asked whether genetic dependencies in CRC models supported this functional relationship between SOX9 and SMARCB1-containing cBAF. Analysis of Broad DepMap datasets revealed dependencies in CRC cell lines and organoid models on SOX9, SMARCB1, SMARCE1, and ARID1A, with weaker effects for other BAF components (Figure S4C). Consistent with these data, SMARCB1, and to a lesser extent ARID1A/ARID1B, was required for HT-29 proliferation and colony formation (Figure S4D-S4E). We previously showed that SMARCB1 suppression reduced growth of HT-29 and patient-derived CRC organoid xenografts *in vivo*^28^; reanalysis of these tumors showed marked KRT20 induction, consistent with differentiation upon SMARCB1 loss (Figure 4C). Across cancer lineages, colon and skin cancer cell lines showed among the highest frequencies of SOX9-SMARCB1 co-dependency (Figure 4D; cut-off >0.5). Within CRC cell lines, SOX9 dependency correlated with ARID1A dependency and, to a lesser extent, SMARCB1 dependency, but showed little relationship with PBRM1 dependency (Figure S4F-S4H). Together, these genetic dependency data support SMARCB1-containing cBAF as a functionally relevant effector of SOX9-dependent stem-like chromatin regulation in CRC. Because SMARCB1 emerged as both a direct SOX9-associated factor and a functional dependency, we next asked how the SOX9 C-terminus engages cBAF and whether this region is sufficient to confer BAF-binding and chromatin-opening activity.

### SOX9 C-terminus encodes a cBAF-engagement module required for enhancer remodeling

The SOX9 C-terminus is required for transcriptional activity and association with BAF cofactors (Figure 1A-B), but whether this region directly controls chromatin remodeling, and which C-terminal domains encode this activity, remains unclear. To address this, we expressed V5-tagged SOX9 C-terminal deletion variants in LS180 CRC cells, chosen for low endogenous SOX9 levels, and profiled chromatin accessibility by ATAC-seq (Figure 5A). Full-length SOX9 (SOX9^FL^) increased chromatin opening whereas deletion of the TA (SOX9^1-379^), TA+PQA (SOX9^1-303^), and TA+PQA+K2 (SOX9^1-228^) domains progressively impaired this activity, with SOX9^1-228^ showing near-complete loss of chromatin-opening function (Figure 5A and Figure S5A). Regions that failed to open after C-terminal deletion were enriched at enhancers linked to stem-like genes, including *PROM1* (Figure 5A and Figure S5B); these only partially overlapped the HT-29 SOX9-cBAF enhancers but converged on the same stem-like target genes, suggesting context-specific enhancer usage. At the *PROM1* locus, both full length and C-terminally truncated SOX9 bound DNA, but only SOX9^FL^ increased chromatin accessibility at enhancers (Figure 5B), indicating that the C-terminus is dispensable for DNA binding but required to convert SOX9 occupancy into productive chromatin remodeling.

**Figure 5.**
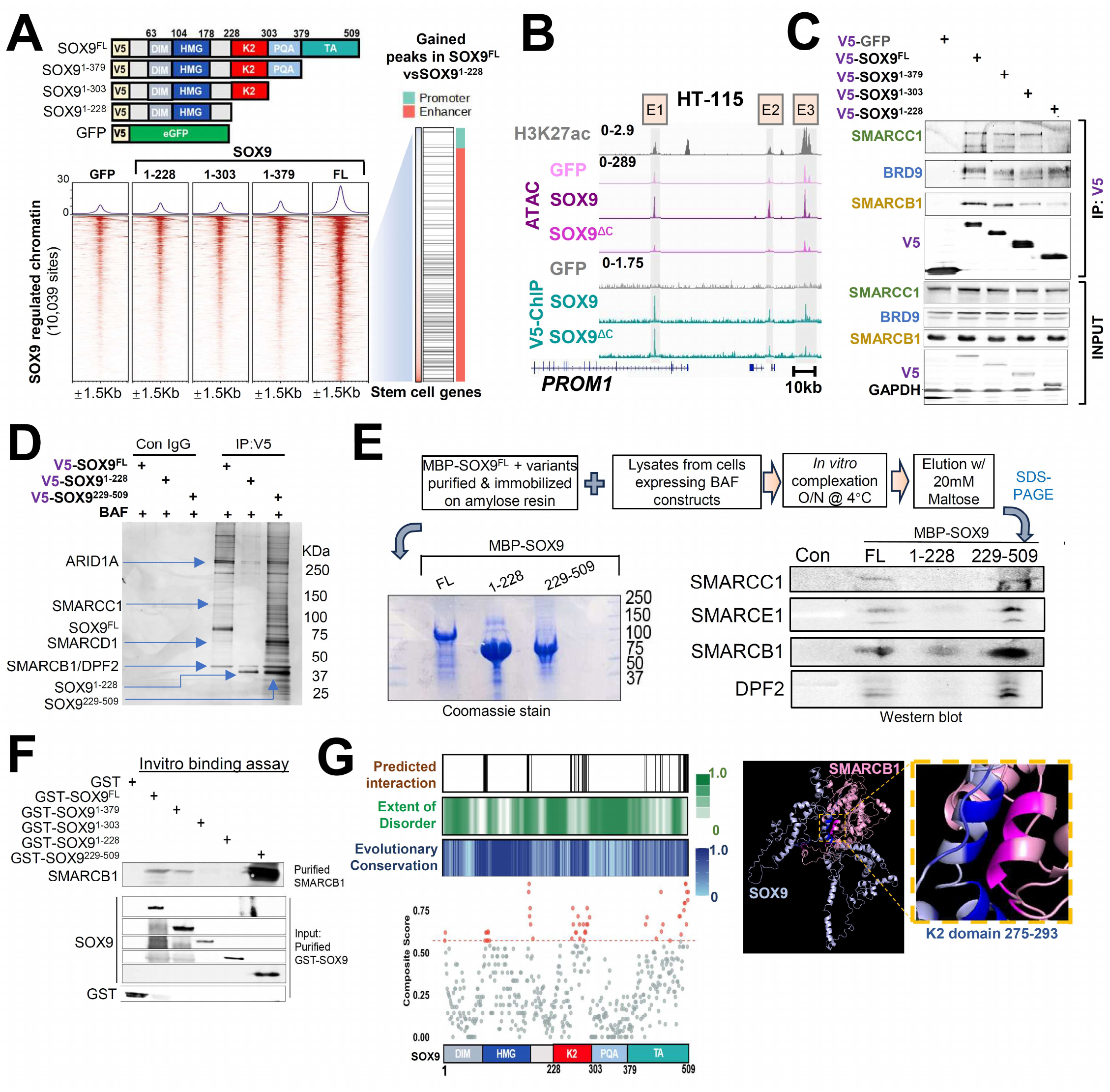
The SOX9 C-terminal IDR encodes a cBAF-recruitment module separable from DNA binding. **(A)** Schematic of SOX9 constructs (top) and ATAC-seq heatmaps showing chromatin accessibility after expression of GFP, SOX9^FL^ (full-length) and SOX9 C-terminal truncation variants (below). Annotation tracks showing stem cell gene set association and promoter/enhancer designation of SOX9^FL^ gained peaks compared to SOX9^1-228^ deletion variant (right). **(B)** Genome browser views the *PROM1* locus, showing an intronic enhancer and two distal enhancers, H3K27ac and V5-ChIP-seq signal from HT-115 cells (Liang et al., 2022), and ATAC-seq peaks from LS180 cells overexpressing GFP, SOX9, or SOX9^1-303^. **(C)** V5 immunoprecipitation followed by immunoblotting for V5, SMARCB1, SMARCC1, and BRD9 in cells expressing GFP, SOX9^FL^, or SOX9 truncation mutants. **(D)** Silver-stained V5 immunoprecipitates from HEK293T cells co-expressing cBAF subunits with SOX9^FL^, SOX9^1-228^, or SOX9^229-509^. **(E)** *In vitro* binding assay using purified MBP-tagged SOX9 variants immobilized on amylose resin and lysates from cells expressing BAF constructs, followed by immunoblotting for cBAF subunits. **(F)** *In vitro* binding assay of purified GST-tagged SOX9 variants with purified SMARCB1. **(G)** Structural models and residue-level annotations of predicted SOX9-SMARCB1 interaction sites, conservation, and disorder.

We next asked whether this chromatin-remodeling function mapped to specific BAF interactions. V5 IP followed by immunoblotting showed that interaction with the ncBAF subunit BRD9 was retained across SOX9 deletion variants, whereas interaction with SMARCB1 progressively diminished with successive loss of C-terminal domains (Figure 5C). SMARCC1 binding was preserved more broadly but was lost with SOX9^1-228^, suggesting a requirement for the K2 domain (Figure 5C and Figure S5C). Together, these data localize the functionally relevant cBAF interaction surface to the SOX9 C-terminus, with the K2 and TA regions emerging as particularly important.

We then tested whether the C-terminus is sufficient to engage cBAF. In HEK293T cells co-expressing cBAF subunits, V5-tagged SOX9^FL^, SOX9^1-228^, and SOX9^229-509^ (TA+PQA+K2) recovered different interactomes: SOX9^1-228^ lost associated proteins, whereas SOX9^229-509^ retained an enriched interactome resembling that of SOX9^FL^ (Figure 5D). In a reconstituted binding system, lysates containing cBAF components were incubated with purified MBP-tagged SOX9 variants immobilized on amylose resin. SOX9^1-228^ showed reduced association with cBAF components, whereas SOX9^229-509^ retained robust complex binding comparable to SOX9^FL^ (Figure 5E). Using purified proteins, we further found that direct SOX9-SMARCB1 interaction progressively weakened with deletion of the TA, PQA, and K2 domains, while SOX9^229-509^ retained direct SMARCB1 binding *in vitro* (Figure 5F). Thus, these cellular, reconstituted, and purified-protein assays show that the SOX9 C-terminus is necessary and sufficient to engage SMARCB1-containing cBAF.

To localize the interface within the C-terminus, we combined mutagenesis with structure prediction. Alanine substitution of conserved residues in the K2 and TA domains partially reduced BAF association, indicating that these residues contribute to, but do not fully account for, the interaction (Figure S5D). This partial, distributed effect is consistent with an intrinsically disordered interface, which typically engages partners through multiple weak contacts rather than a single dominant binding surface. Consistent with this, RoseTTAFold^36^- and ClusPro^37^-based modeling nominated the same K2 and TA regions, conserved residues with comparatively lower predicted disorder, as candidate SMARCB1-interacting surfaces (≤8Å Cα–Cα distance) (Figure 5G), and reciprocally nominated the SMARCB1 RPT region as a candidate binding interface (Figure S5E). Alphafold3 (AF3) modeling of a larger SOX9-cBAF assembly further raised the possibility that disordered regions of SOX9, ARID1A, and other BAF components adopt more ordered conformations upon complex assembly (Figure S5F). Supporting a broader interaction surface with cBAF, reanalysis of published mass spectrometry data^38^ showed ARID1A associating with SOX9 at a magnitude comparable to other BAF cofactors, and TurboID mapping indicated that ARID1A-SOX9 interaction depends on ARID1A IDR1 rather than its DNA-binding domain (Figure S5G). Together, these data map the cBAF-engaging activity of SOX9 to a conserved, likely multivalent interaction surface within the K2 and TA regions of the C-terminus.

### SMARCB1 docking partially restores truncated SOX9 function

Having defined the SOX9 C-terminus as the functionally relevant cBAF-binding module, we next asked whether the defect of SOX9^1-228^ reflects failed SMARCB1/cBAF docking. To test this, we generated a SOX9^1-228^-SMARCB1 fusion construct (SOX9^1-228^-B1) and examined whether enforced SMARCB1 linkage could restore cBAF engagement, chromatin remodeling, and stem-like activity (Figure 6A). This design directly tests whether SMARCB1/cBAF recruitment is a limiting function encoded by the SOX9 C-terminus.

**Figure 6.**
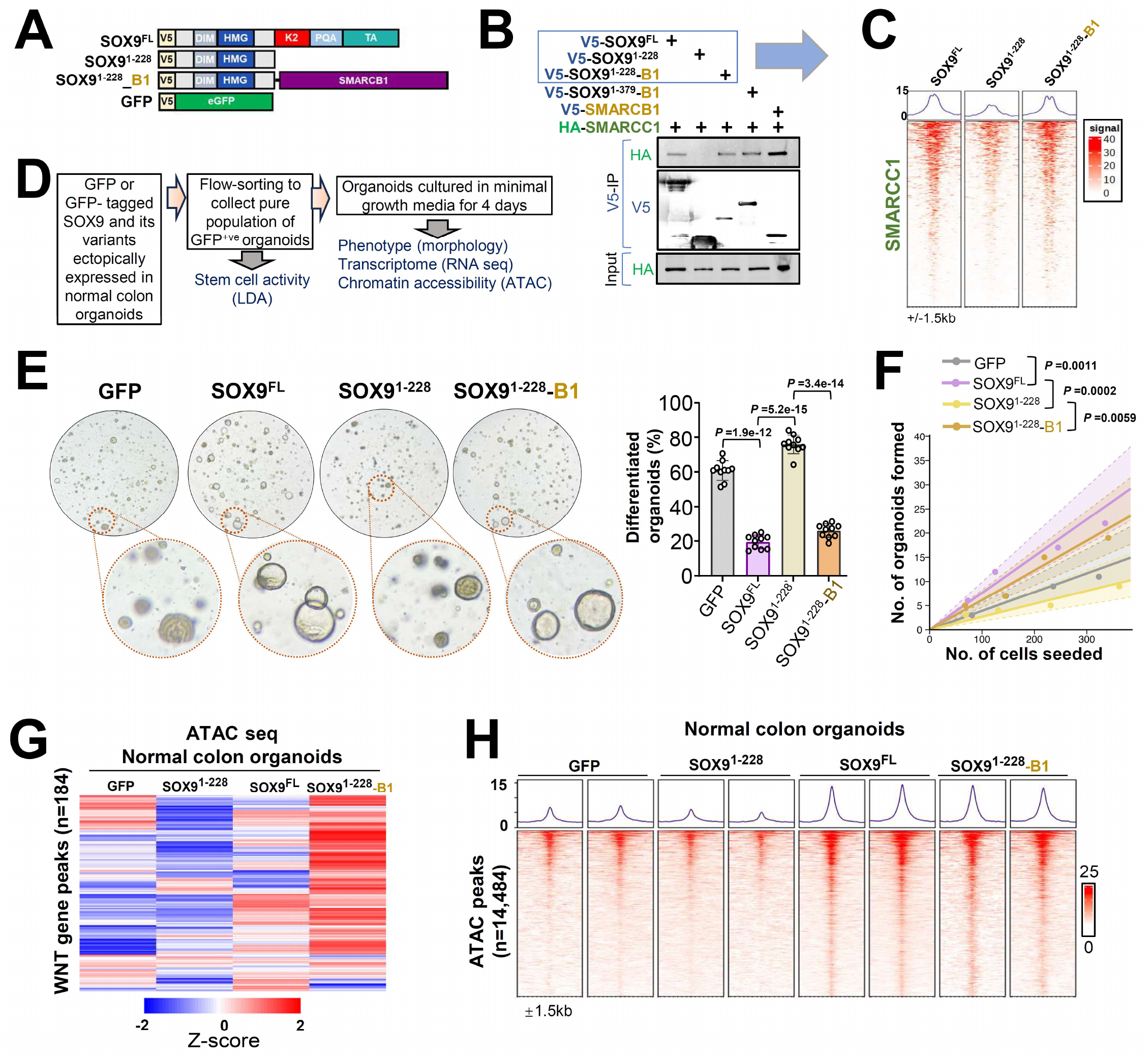
Enforced SMARCB1 docking reconstructs SOX9-dependent enhancer remodeling and self-renewal. **(A)** Schematic of SOX9^FL^, SOX9^1–228^, SOX9^1–228^-SMARCB1 constructs **(B)** V5 immunoprecipitation followed by immunoblot analysis of SMARCC1 in cells expressing SOX9^FL^, SOX9^1–228^, SOX9^1–228^-SMARCB1, SOX9^1–379^-SMARCB1or SMARCB1. **(C)** SMARCC1 chromatin occupancy heatmaps in LS180 cells expressing SOX9^FL^, SOX9^1–228^, SOX9^1–228^-SMARCB1 constructs. **(D)** Workflow of normal colon organoid expressing GFP-tagged construct experiments **(E)** Representative brightfield images (left) and quantification of differentiated organoids (right) of normal colon organoids expressing GFP, SOX9^FL^, SOX9^1–228^, SOX9^1–228^-SMARCB1 constructs under reduced-factor culture conditions. **(F)** Graph shows results from Limiting Dilution Assay plotted as number of organoids formed against number of cells seeded. **(G)** Bulk RNA-seq heatmap of WNT target gene expression across normal colon organoids expressing GFP, SOX9^FL^, SOX9^1–228^, SOX9^1–228^-SMARCB1 **(H)** Bulk ATAC-seq heatmap of WNT-associated peaks across normal colon organoids expressing GFP, SOX9^FL^, SOX9^1–228^, SOX9^1–228^-SMARCB1

We first asked whether enforced SMARCB1 linkage restored molecular engagement with cBAF. Whereas SOX9^1-228^ lost association with cBAF components, fusion to SMARCB1 restored interaction with SMARCC1 to levels comparable to those observed with SOX9^FL^ and SMARCB1 controls (Figure 6B). To determine whether this biochemical rescue extends to chromatin, we assessed SMARCC1 genome occupancy in LS180 cells. SMARCC1 occupancy at SOX9-bound loci was lost upon C-terminal truncation and re-established by the fusion, with SOX9^1-228^-B1 restoring SMARCC1 binding at the specific sites depleted in SOX9^1-228^ (Figure 6C). Because SOX9^1-228^ binds DNA but cannot open chromatin (Figure 5B), the fusion restores cBAF recruitment at these specific loci by targeting SMARCB1 to SOX9-bound sites.

We next asked whether this rescue is sufficient to restore stem-like behavior in a biological system. We leveraged normal colon organoids, which differentiate over time in reduced-factor culture conditions, and introduced GFP-tagged SOX9^FL^, SOX9^1-228^ or SOX9^1-228^-B1 expression constructs (Figure 6D). SOX9^FL^ blocked differentiation and maintained a stem-like cystic morphology, whereas SOX9^1-228^ organoids differentiated similarly to GFP controls (Figure 6E). Strikingly, SOX9^1-228^-B1 substantially restored the stem-like phenotype, suppressing organoid differentiation and promoting cystic growth approaching that induced by SOX9^FL^ (Figure 6E). In limiting dilution assays, SOX9^1-228^-B1 also significantly restored organoid-forming capacity relative to SOX9^1-228^ (Figure 6F and Figure S6A). Thus, enforced SMARCB1 docking partially restores both phenotypic and functional stem-like properties lost upon SOX9 C-terminal truncation.

To determine whether phenotypic rescue was accompanied by recovery of transcriptional and chromatin states, we performed bulk RNA-seq and ATAC-seq. SOX9^1-228^-B1 partially restored the stem-like transcriptional program that was activated by SOX9^FL^ and absent in SOX9^1-228^ and GFP controls (Figure S6B). WNT signaling was robustly restored in SOX9^1-228^-B1 expressing organoids, in some cases exceeding levels observed with SOX9^FL^, consistent with the engineered nature of enforced SMARCB1 linkage (Figure S6C). ATAC-seq showed re-establishment of accessible chromatin at WNT target genes in SOX9^1-228^-B1, which remained closed in SOX9^1-228^ controls (Figure 6G). This rescue extended globally, with SOX9^1-228^-B1 restoring a substantial fraction of the chromatin-opening program observed in SOX9^FL^ and lost in SOX9^1-228^ (Figure 6H). Although the fusion construct also induced distinct transcriptional changes, a dominant effect was recovery of the stem-like transcriptional and chromatin program impaired by SOX9 C-terminal deletion. Together, these findings identify SMARCB1 docking as a central effector function of the SOX9 C-terminus and show that enforced recruitment of SMARCB1-containing cBAF is sufficient to partially restore SOX9-dependent stem-like chromatin regulation.

### The SOX9 C-terminus encodes a transferable cBAF-recruitment module for chromatin opening

Having established that the SOX9 C-terminus is required for cBAF engagement and that enforced SMARCB1 docking can partially restore the activity of C-terminally truncated SOX9, we next asked whether the isolated C-terminus is sufficient to execute a chromatin-remodeling function. To test whether it can support assembly of a higher-order cBAF complex, amylose-purified SOX9^229-509^ was subjected to 10%-30% glycerol gradient sedimentation. Although comparable in size to SOX9^1-228^, SOX9^229-509^ shifted into higher-molecular weight fractions and co-sedimented with SMARCC1, SMARCD1, SMARCB1, and SMARCE1, whereas SOX9^1-228^ did not (Figure 7A). Thus, the SOX9 C-terminus is sufficient not only to bind individual BAF subunits (Figure 5F), but also to support assembly of a higher-order cBAF-containing complex.

**Figure 7.**
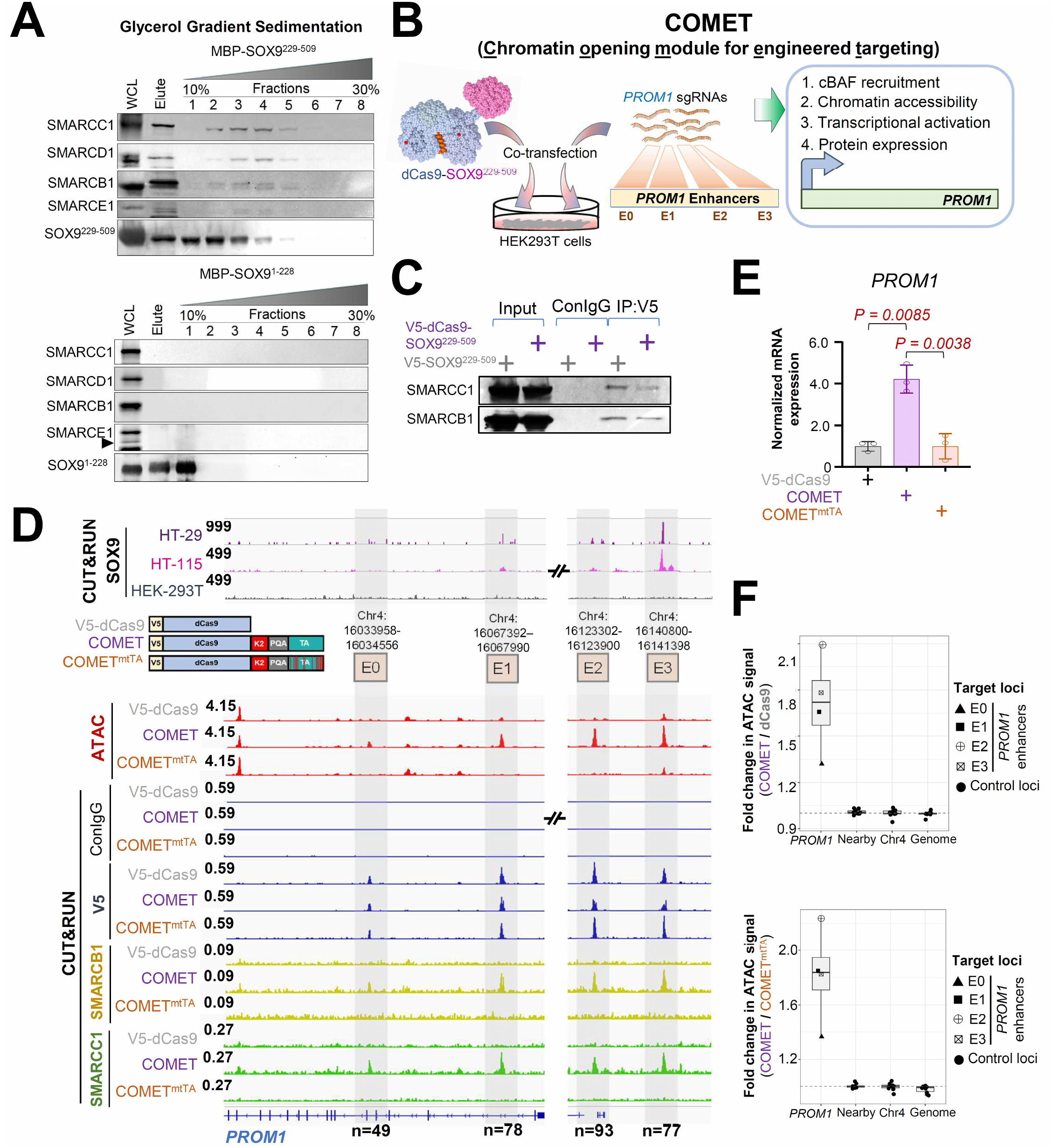
The SOX9 C-terminal module is transferable and directs endogenous cBAF for programmable chromatin opening. **(A)** Glycerol gradient fractionation of complexes associated with SOX9^229-509^ and SOX9^1-228^ followed by immunoblotting for SMARCC1, SMARCD1, SMARCB1, SMARCE1, and SOX9. **(B)** Workflow of Chromatin opening module for engineered targeting (COMET) and schematic of dCas9-SOX9 fusion constructs **(C)** Immunoblot analysis of SMARCC1 and SMARCB1 in V5 immunoprecipitates from cells expressing SOX9^229-509^ or dCas9-SOX9^229-509^. **(D)** Genome browser views of ATAC-seq signal and CUT&RUN signals (endogenous SOX9 in HT-115 and LS-180 CRC cells and non-CRC HEK293T cells; V5, SMARCC1, SMARCB1 in HEK293T cells) at the PROM1 enhancers after targeting dCas9, dCas9-SOX9^229-509^ or dCas9-SOX9^229-509^ -mutantTA with pooled sgRNAs. **(E)** PROM1 mRNA expression in COMET experiment using qRT-PCR. **(F)** Analysis of fold change in ATAC peak signal at *PROM1* enhancers targeted by dCas9-SOX9^229-509^ or dCas9-SOX9^229-509^ -mutantTA

We next asked whether this cBAF-recruitment module could be transferred to an engineered DNA-targeting context. We fused SOX9^229-509^ to catalytically inactive dCas9 to generate COMET, a <u>c</u>hromatin-<u>o</u>pening <u>m</u>odule for <u>e</u>ngineered targeting, and directed it to selected regulatory elements with sgRNAs (Figure 7B). dCas9-SOX9^229-509^ (COMET) retained interaction with SMARCC1 and SMARCB1, indicating that fusion to dCas9 preserves cBAF-binding (Figure 7C). We first targeted COMET in HEK293T cells to the PROM1 locus, where endogenous SOX9 occupancy and PROM1 expression are minimal compared to CRC cell lines (Figure 7D). The sgRNA pool (Table S7) targeted three PROM1 enhancers that are SOX9-bound in CRC cells (E1-E3) and one local intronic element (E0) that is not detectably occupied by SOX9.

Both dCas9 alone and COMET occupied the targeted PROM1 elements, indicating efficient guide-directed DNA binding by both constructs (Figure 7D). However, dCas9 alone did not recruit cBAF or open chromatin. In contrast, COMET recruited endogenous SMARCB1 and SMARCC1, increased local chromatin accessibility at targeted PROM1 elements, and activated PROM1 mRNA and protein expression (Figure 7D-E and Figure S7A). Chromatin opening was concentrated at targeted PROM1 elements and was not observed at nearby chromosome 4 control regions or genome-wide control loci (Figure 7F), indicating that COMET induces local rather than global accessibility changes.

To determine whether this activity depends on the cBAF-engagement surface defined above, we generated COMET^mtTA^, in which SOX9 C-terminal TA residues that contribute to BAF interaction were substituted with alanine. COMET^mtTA^ retained guide-directed chromatin targeting but failed to recruit SMARCB1 and SMARCC1 efficiently, did not increase accessibility at targeted PROM1 elements, and showed markedly reduced PROM1 activation compared with COMET (Figure 7D-F). Thus, DNA targeting is not sufficient for remodeling; the SOX9 C-terminus supplies a cBAF-recruitment function that converts dCas9 occupancy into local chromatin opening and transcriptional activation.

Finally, we tested whether COMET activity was restricted to the native SOX9 regulatory context. Using independent sgRNA pools, we targeted COMET to two unrelated chromosome 1 loci, including a regulatory region near *LGR6*. At both loci, COMET but not dCas9 increased local chromatin accessibility (Figure S7B-C). These results show that the SOX9 C-terminus can be removed from its native protein context and redeployed to remodel non-native genomic sites.

Together, these findings show that the SOX9 C-terminus is sufficient to support higher-order cBAF assembly and to redirect endogenous cBAF to sgRNA-targeted chromatin. COMET extends the biological mechanism defined here into an engineered targeting strategy, demonstrating that a disease-co-opted transcription-factor IDR can encode a transferable cBAF-recruitment function for programmable chromatin opening.

## DISCUSSION

Here, we identify the SOX9 C-terminus as an intrinsically disordered, transferable cBAF-recruitment module that couples DNA binding to enhancer remodeling. These findings separate genomic occupancy and cofactor recruitment, two TF properties that are often treated as one. The SOX9 C-terminal IDR preferentially recruits SMARCB1-containing canonical BAF (cBAF) to bound enhancers, and this activity can be disrupted while SOX9 occupancy remains intact, functionally reconstructed by enforced SMARCB1 docking, and transferred to an unrelated DNA-targeting platform. During colorectal cancer initiation, this recruitment activity operates at stem-like enhancers that become accessible during *APC*-mutant adenoma formation and remain SOX9-dependent in established CRC. Together, these findings define remodeler recruitment as a separable effector step between DNA recognition and chromatin-state control.

The forced-docking experiments provide a direct causal test of this model. Restoring SMARCB1 engagement to an otherwise remodeling-defective SOX9 truncation reconstructs a substantial fraction of the chromatin and transcriptional program, arguing that cBAF recruitment is a limiting effector activity of the C-terminal domain. The rescue is incomplete, indicating that SMARCB1/cBAF recruitment is a major but not exclusive function of this region. Additional contacts with other cBAF subunits, cooperative interactions with transcriptional cofactors, or C-terminal contributions downstream of remodeling may tune enhancer selectivity and transcriptional output. Nevertheless, functional reconstruction through an enforced cofactor contact provides direct evidence that recruitment of the remodeling machinery is a critical step linking SOX9 occupancy to chromatin state.

A broader implication is that TF IDRs can encode specificity for endogenous chromatin-remodeling machinery. IDRs have been widely implicated in transcriptional activation, multivalent protein interactions, and condensate-associated behaviors, but our findings support an additional function in which an IDR serves as a modular effector-recruitment domain. In SOX9, this activity is distributed across conserved regions of the C-terminus, and biochemical reconstitution, mutagenesis, and structural prediction are more consistent with a distributed interaction surface than with a dominant residue or compact binding motif. Such an architecture may be particularly suited to lineage TFs, which must couple stable sequence-specific DNA recognition to context-dependent engagement of regulatory machinery.

Our findings complement an emerging framework of direct TF-BAF engagement. Recent work identified the SWIFT/SMARCD platform as a remodeler-side determinant of transcription-factor binding preferences^6^, while ARID1A-dependent cBAF coordinates sequential TF activity during B-cell fate control and lymphomagenesis^39^. SOX9 has also been shown to act as a pioneer factor capable of opening lineage enhancers and reshaping cell fate in skin stem cells^40^. Our study defines the reciprocal transcription-factor-side logic: an IDR-encoded recruitment module is required to deploy a specific endogenous BAF assembly and its missing activity can be reconstructed by enforced SMARCB1 docking. SOX9 therefore provides a mechanistic example of how lineage TFs can encode not only where they bind, but which chromatin-remodeling machinery is deployed after binding. In this view, SOX9 is not only a marker of plasticity^41–44^, but a modular effector that couples lineage-specific genomic occupancy to cBAF-dependent enhancer remodeling.

This recruitment mechanism operates during a defined cell-state transition in CRC. Although *APC* loss initiates neoplasia, mutation alone does not specify the enhancer architecture that stabilizes the resulting abnormal cell state. Premalignant epithelial cells across tissues can acquire regenerative, stress-associated, or niche-remodeling programs before overt cancer formation^27,30,45^, and SOX9 has emerged as a recurrent feature of several such states^19,29–31^. Our findings provide a direct chromatin-remodeling mechanism for this biology. During *APC*-mutant adenoma formation, SOX9-cBAF regulatory elements selectively gain accessibility in aberrant stem-like cells, while established CRC cells remain acutely dependent on SOX9 for cBAF occupancy, enhancer accessibility, and stem-like identity. We therefore favor a model in which *APC* alteration creates a context in which SOX9 recruits SMARCB1-containing cBAF to selected enhancers, helping install and maintain a differentiation-resistant chromatin state that supports lesion persistence and progression. This model describes how a new regulatory state is built, rather than how cells move between pre-existing states. That distinction is conceptually consistent with recent evidence that transitions between established cell states during KRAS inhibition can occur with comparatively limited chromatin remodeling^46^.

Our results also refine how BAF complex function can be interpreted across regulatory contexts. Although SOX9 associates with cBAF, pBAF, and ncBAF, SOX9-dependent remodeling is preferentially associated with cBAF at enhancers, whereas BAF occupancy at promoter-proximal regions is less selective and can be partially buffered by other assemblies after SOX9 loss. This distinction may help reconcile the context-dependent effects of BAF perturbation in intestinal homeostasis and *Apc*-mutant tumorigenesis^47,48^. Rather than functioning as interchangeable remodeling machines, BAF assemblies may acquire regulatory specificity through the combination of TF context, complex composition, and regulatory-element class. In this view, the biological consequence of perturbing a BAF subunit depends not only on the biochemical complex affected but also on which transcription-factor-defined enhancer compartment that complex serves.

The transferability of the SOX9 C-terminal module provides an orthogonal test of this model. Most engineered chromatin regulators deliver an exogenous catalytic or transcriptional effector to a chosen locus^49–51^. In contrast, COMET uses the SOX9 C-terminus to recruit endogenous cBAF. The isolated domain supports higher-order cBAF assembly as shown by glycerol-gradient sedimentation, and when fused to dCas9 directs endogenous SMARCB1 and SMARCC1 to sgRNA-specified loci, increasing local accessibility and transcription. Importantly, the cBAF-binding-deficient COMET^mtTA^ variant retains guide-directed targeting but loses efficient remodeler recruitment and chromatin opening, separating genomic localization from effector function within the engineered system itself. Activity at non-native loci further indicates that this recruitment function is not restricted to endogenous SOX9 binding sites. COMET therefore demonstrates that the molecular activity encoded by the SOX9 IDR is transferable and provides a route for programmable chromatin opening through recruitment of endogenous remodeling machinery.

More broadly, these findings suggest that TF IDRs can function as modular selectors of chromatin machinery rather than solely as generic activation domains. Such recruitment logic may help explain how alterations that preserve TF abundance or DNA binding nevertheless disrupt enhancer selection, cell identity, or disease-associated plasticity^52–55^. The SOX9–SMARCB1/cBAF interface also provides a defined molecular interaction that could potentially be targeted to destabilize differentiation-resistant states, although whether this interface is pharmacologically tractable remains unknown. The more general principle is that cell-state control can reside in the molecular step between DNA recognition and chromatin remodeling - namely, which endogenous effector complex a transcription factor recruits after binding DNA.

### STUDY LIMITATIONS

Although our data link SOX9-cBAF enhancer activity to *APC*-mutant adenoma formation and establish the requirement for SOX9-dependent cBAF recruitment in CRC models, the interface has not yet been perturbed genetically *in vivo*. Thus, the contribution of the SOX9 C-terminal recruitment module to adenoma initiation or maintenance will require direct testing with interface-selective alleles. In addition, although domain mapping, biochemical reconstitution, mutagenesis, and computational modeling converge on a distributed C-terminal interaction with SMARCB1-containing cBAF, the structural basis of this interaction awaits higher-resolution definition. The partial rescue achieved by enforced SMARCB1 docking indicates that SMARCB1/cBAF recruitment is a major, but not exclusive, function of the SOX9 C-terminus. Finally, enforced SMARCB1 docking and COMET are engineered sufficiency systems. They establish mechanistic modularity and transferability, but broader locus testing and *in vivo* validation will be required to define the generality of IDR-encoded remodeler recruitment across SOX9-dependent tissues and other transcription factors.

## METHODS

### Cell and Organoid culture

All cell lines and organoid cultures were maintained in humidity-controlled incubator maintained at 37°C with 5% CO2. The human colorectal cancer cell lines (HT-115, HT-29 (HTB-38) and LS180 (CL-187 )) and HEK 293 T were maintained in DMEM supplemented with 10 % FBS and 1 % penicillin/streptomycin. Intestine or colon-derived organoids were maintained in DMEM/F12 media containing WRN and supplemented with HEPES (Sigma-Aldrich), 20% FBS (Gibco), 1% penicillin-streptomycin, and 10 mM RhoK inhibitor Y-27632 (Sigma-Aldrich). For passage, colon organoids were dispersed by trypsin-EDTA and transferred to fresh Matrigel. Passage was performed every 3 to 4 days with a 1:3 to 1:5 split ratio.

### Rapid Immunoprecipitation Mass Spectrometry of Endogenous Proteins

Rapid Immunoprecipitation Mass Spectrometry of Endogenous Proteins (RIME) was performed as previously described by Hisham Mohammed et al^32^. Briefly, cells were crosslinked with 1% formaldehyde for 8 minutes at RT and quenched with 0.1 M glycine. Nuclei were isolated in buffer containing 50 mM HEPES-KOH, (pH 7.5), 140 mM NaCl, 1 mM EDTA, 10% (vol/vol) glycerol, 0.5% (vol/vol) NP-40, 0.25% (vol/vol) Triton X-100 and protease inhibitor cocktail (Rosche). Nuclear fraction was lysed in buffer containing 10 mM Tris-HCl (pH 8.0), 100 mM NaCl, 1 mM EDTA, 0.5 mM EGTA, 0.1% (wt/vol) sodium deoxycholate, 0.5% (vol/vol) *N*-lauroylsarcosine and protease inhibitor cocktail. followed by sonication to generate fragmented chromatin. Clarified lysates were incubated overnight at 4°C with V5 antibody (Abcam #ab9116) immobilized on Agarose beads. Beads were subjected to wash with buffer containing 50 mM HEPES (pH 7.6), 1 mM EDTA, 0.7% (wt/vol) sodium deoxycholate, 1% (vol/vol) NP-40 and 0.5M LiCl followed by two washes of 100 mM ammonium bicarbonate before overnight on-bead tryptic digestion. Peptides were desalted using C18 solid-phase extraction columns and analyzed by liquid chromatography–tandem mass spectrometry (LC-MS/MS). Proteins identified in matched IgG control immunoprecipitations were considered nonspecific contaminants and excluded from downstream analyses.

### Immunoprecipitation Assays

Cells or organoids were lysed for 30 minutes at 4°C in buffer containing 25mM Tris-Cl (pH 7.5), 150mM NaCl, 2mM PMSF, 2mM Na3VO4, 10mM NaF, 10µM N-ethyl maleimide (NEM), 1x Protease inhibitor cocktail (Rosche), 2mM EDTA and 0.5% NP40. Lysates were centrifuged at 16000g for 10 minutes at 4°C to clear the debris. Cleared lysates were incubated overnight at 4°C with control IgG or desired antibodies conjugated to agarose beads. Following incubation, unbound lysate was removed and beads were washed with buffer containing 25mM Tris-Cl (pH 7.5), 200mM NaCl, 2mM PMSF, 2mM Na3VO4, 10mM NaF, 10µM N-ethyl maleimide (NEM), 1x Protease inhibitor cocktail (Rosche) and 2mM EDTA. Immunoprecipitated proteins were eluted in Laemmli sample buffer supplemented with 1 mM DTT and analyzed by western blotting. Antibodies used in IP assays are listed in Table S8.

### Animal studies

All animal procedures were approved by the Dana-Farber Cancer Institute Institutional Animal Care and Use Committee (IACUC; protocol 11-009). *Lgr5*^eGFP-IRES-CreERT2^ mice were previously described, and backcrossed onto a C57BL/6J background (>97% purity; Taconic). Conditional intestinal Apc deletion was achieved by crossing Apc^f/f^ mice (a gift from C. Perret) to *Lgr5*^eGFP-IRES-CreERT2^ mice. For lineage tracing, mice were further crossed with R26-LSL-tdTomato mice (JAX #007905) (70), and Sox9 deletion was induced using Sox9^flox^ mice (JAX #013106).

Conditional alleles were activated in 6–8-week-old mice by a single intraperitoneal tamoxifen injection (50 μL sunflower oil containing 10 mg/ml tamoxifen), unless otherwise indicated. For the MNU model, *Lgr5*^eGFP-IRES-CreERT2^ mice received 240 ppm MNU (bioKEMIX) in drinking water every other week for 10 weeks and were monitored for 1 year. For the AOM/DSS model, mice (8–10 weeks old) were injected with AOM (10 mg/kg; Sigma-Aldrich) followed by three cycles of 2% DSS (MP Biomedicals) administered in drinking water for 5 days every 4 weeks. Mice were euthanized 11 weeks after AOM injection. At endpoint, intestinal tissues were collected, flushed with PBS, fixed in 10% formalin, Swiss-rolled, and paraffin-embedded. Fresh tissues were processed for organoid culture, RNA and protein isolation, or flash-frozen for long-term storage.

### Histopathology and IHC

Paraffin-embedded intestines, organoids, or xenograft tumors were serially sectioned and mounted on a slide. Sections were subjected to hematoxylin and eosin (H&E), AB–periodic acid–Schiff (AB-PAS), as well as immunostaining, using standard procedures. For morphological analysis, sections were serially dehydrated in xylene and ethanol and stained with H&E for histological assessment or AB-PAS to identify goblet cells and mucus. For IHC, antigen retrieval was performed using a sodium citrate buffer (pH 6), Trilogy (Sigma-Aldrich Cell Marque), or tris-EDTA (pH 9). Slides were permeabilized using a 0.2% Triton X-100 for 30 minutes at room temperature and blocked with donkey serum for 1 hour. The binding of primary antibody was detected with 3,39-diamino-benzidine-tetrahydrochloride-dihydrate and counterstained with hematoxylin.

### Flow analysis

Intestines from euthanized mice were washed with ice-cold PBS, villi were removed by scraping, and epithelial cells were dissociated in 5 mM EDTA/PBS at 4°C for 20 minutes with intermittent shaking. The epithelial fraction was collected by centrifugation, incubated in prewarmed 4× TrypLE at 37°C for 30 minutes, filtered through a 70 μm strainer, and resuspended in FACS buffer (2%BSA). Organoids were trypsinized to make single cell suspensions and passed through 70 μm strainer before sorting. Cells were trypsinized to make single-cell suspension and passed through 70 μm and/or 40 μm filters before sorting. Sorted cells were collected in 50% FBS/DMEM containing 10 μM Y27632

### Single-cell RNA-seq

Murine intestinal samples were processed as described above and subjected to scRNA-seq with or without cell hashing. For hashing, mouse cells were labeled with TotalSeq Hashtag antibodies (BioLegend #155861 and #155863); Viable cells were resuspended in PBS containing 0.04% BSA at 1,000 cells/μL, and ∼17,000 cells were loaded onto the 10x Genomics Chromium platform. scRNA-seq and cell hashing libraries were generated using Chromium Single Cell 5′ kits (10x Genomics), quality-checked using a Bioanalyzer High Sensitivity DNA Kit (Agilent) and sequenced on an Illumina NovaSeq S4 platform (150-bp paired-end). Sequencing data were processed using Cell Ranger v3.1.0 and aligned to the mm10 reference genome

### Single-nuclear RNA-seq using combinatorial indexing

snRNA-seq of HT-29^SOX9-dTAG^ cells was performed using the “EasySci” RNA protocol (10.1038/s41588-023-01572-y). Briefly, HT-29^SOX9-dTAG^ cells were treated with DMSO (control) or dTAG-^v^1 at a 1μM concentration for various time periods (1 to 6 days). Media were changed every other day. Cells were harvested, and nuclei were extracted and fixed following the “EasySci” protocol. 100,000 nuclei were distributed to each well of the first RT plate. A total of 600,000 nuclei per treatment time point and 1,200,000 nuclei for the DMSO control were distributed into the RT plate. Reverse transcription, ligation, second-strand synthesis, and tagmentation were performed following the standard EasySci protocol. Libraries were prepared by PCR using the first 60 barcodes and sequenced using an Illumina NovaSeq X (paired end, 150 bp).

### Assay for transposase-accessible chromatin sequencing

ATAC-seq libraries were prepared as previously described (72). Briefly, duplicate samples of 50000 flow-sorted cells were lysed to isolate nuclei and subjected to transposition using TDE1 enzyme (Illumina, 20034197). Tagmented DNA was purified using the Zymo DNA Clean & Concentrator-5 kit (D4014), PCR-amplified with NEBNext 2× Master Mix and Illumina adapters and purified using AMPure XP beads (Beckman Coulter). Libraries were sequenced as 150-bp paired-end reads on an Illumina NovaSeq platform.

### CUT&RUN

CUT&RUN experiments in cells or organoids were performed using CUTANA ChIC/CUT&RUN kit (Epicypher, #14-1048) following the manufacturer’s protocol. Briefly, 5 × 10^5^ cells were captured with concanavalin A, permeabilized using 0.01% digitonin, and incubated with 0.5 μg of desired antibody in 100 μL of antibody buffer [20 mM Hepes at pH 7.5, 150 mM NaCl, 0.5 mM spermidine, 1× protease inhibitor cocktails (EDTA-free tablet; Roche), 2 mM EDTA, and 0.01% digitonin] overnight. List of primary antibodies used in CUT&RUN experiments are listed in Table S8. After removing unbound antibody, pAG-MNase (20×) in 50 μL of cell permeabilization buffer was added to the cells and incubated for 10 min at room temperature. Then, CaCl2 (2 mM) was added to activate MNase and incubated at 4°C for 2 hours. The reaction was stopped using 33 μ L of 2× STOP buffer and *Escherichia coli* spike-in DNA (0.5 ng) was added as a control. The DNA from the released chromatin in the supernatant was purified and then quantified using Qubit dsDNA HS assay kit (Agilent Technologies). CUT&RUN libraries were constructed using a NEBNext Ultra II DNA library preparation kit as per manufacturer’s protocol with a few modifications. Briefly, end repair and dA-tailing were conducted on 6 ng of CUT&RUN eluted DNA for 30 min at 20°C followed by 1 hour at 50°C. After adaptor ligation for 30 min at 20°C, the DNA fragments were purified by 1.75× volume of AMPure XP beads (Beckman Coulter) followed by 10 to 12 cycles of PCR amplification with Next Ultra II Q5 master mix. The PCR products were purified with 1× volume of AMPure XP beads. After quantitative and qualitative analysis, libraries with different indexes were pooled and sequenced on Illumina NovaSeq platform with paired-end 150-bp reads.

### Cell proliferation assays

To assess cell viability cellular ATP levels were measured by using the CellTiter-Glo viability assay (Promega) according to the manufacturer’s instructions. Experiments were performed in triplicate in 96-well plates. Colony formation assay was performed by plating 500 live cells / well of a 12-well plate. Cells were grown for 12 days with desired treatment, then fixed with 10% buffered formalin and stained with crystal violet. All experiments were performed in triplicates.

### Lentivirus generation and transduction

To generate lentiviruses, expression vectors were co-transfected into HEK293T cells with the lentiviral packaging constructs psPAX2 and pMD2.g (vesicular stomatitis virus glycoprotein) in a 1:1:1 ratio using X-tremeGENE 9 DNA Transfection Reagent (Roche) according to the manufacturer’s instructions. Cell culture medium was changed the following day, and lentiviral supernatant was harvested 48 and 72 hours later and filtered through a 0.45-μm filter (Millipore). Lentiviruses were aliquoted and stored at −80°C until use. To transduce, organoids were trypsinized to make single-cell suspension, and cells were resuspended in 500 μL of lentiviral supernatant with polybrene (8 μg/ml) and 10 μM Y-27632, centrifuged at 600*g* at 32°C for 1 hour, and incubated for 6 hours in a culture incubator at 37°C and 5% CO2. The transduced cells were suspended in 30 to 50 μL of Matrigel and cultured with WRN medium containing 10 μM Y-27632. Pure lines were generated by either antibiotic selection or by repeated flow sorting. For transduction in cells, Lentivirus was resuspended in culture media containing polybrene (8 μg/ml) and added to cells plated at 70% confluency. Cells were grown for 2-3 days and then selected for pure population by growing in appropriate antibiotics or by repetitive flow-sorting.

### Purification of SOX9

MBP-tagged SOX9 or its truncation variants were transfected using PEI (plasmid:PEI ratio of 1:5) in suspension Expi293 cells for 48 hours. Cells were harvested by centrifugation (1500g; 15minutes; 4°C) and lysed in buffer containing 25mM Tris-Cl (pH 7.5), 150mM NaCl, 2mM PMSF, 2mM Na3VO4, 10mM NaF, 10µM N-ethyl maleimide (NEM), 1x Protease inhibitor cocktail (Rosche), 2mM EDTA and 0.5% NP40. Lysates were incubated with amylose resins at 4°C for 16 hours. Unbound supernatant was discarded and resins were washed with wash buffer containing 25mM Tris-Cl (pH 7.5), 200mM NaCl, 2mM PMSF, 2mM Na3VO4, 10mM NaF, 10µM N-ethyl maleimide (NEM), 1x Protease inhibitor cocktail (Rosche) and 2mM EDTA. Integrity of the purified protein was analyzed by Coomassie staining. Purified proteins were stored at 4°C in buffer containing 25mM Tris-Cl (pH 7.5), 150mM NaCl, 2mM PMSF, 1x Protease inhibitor cocktail (Rosche) and 25% glycerol.

For SOX9-pulldown experiments, lysates of Expi293 cells expressing BAF components were incubated with purified SOX9, immobilized on amylose resins for 4 hours at 4°C. Unbound lysates were discarded and bound proteins were washed three times followed by western blot analysis.

### Glycerol gradient sedimentation assay

MBP-tagged SOX9^229-509^ or SOX9^1-228^ were co-transfected with cBAF components using PEI (plasmid:PEI ratio of 1:5) in suspension Expi293 cells for 48 hours. Cells were harvested by centrifugation (2500g; 30 minutes; 4°C) and lysed in buffer containing 25mM Tris-Cl (pH 7.5), 150mM NaCl, 2mM PMSF, 2mM Na3VO4, 10mM NaF, 10µM N-ethyl maleimide (NEM), 1x Protease inhibitor cocktail (Rosche), 2mM EDTA and 0.5% NP40. Lysates were incubated with amylose resins at 4°C for 16 hours. Unbound supernatant was discarded and resins were washed with buffer containing 25mM Tris-Cl (pH 7.5), 200mM NaCl, 2mM PMSF, 2mM Na3VO4, 10mM NaF, 10µM N-ethyl maleimide (NEM), 1x Protease inhibitor cocktail (Rosche) and 2mM EDTA. Bound complexes were eluted with wash-buffer supplemented with 20mM Maltose for 2 hours at 4°C. Eluted SOX9-BAF complexes were concentrated by passing through >100KDa filter. Concentrated protein samples were loaded on 10-30% dis-continuous glycerol gradient and sedimentation performed at 100,000g for 16 hours at 4°C. 12 equal fractions were collected, dialyzed to remove glycerol content, concentrated using a >100KDa filter and analyzed by western blot.

### Limiting Dilution Assay

For limiting dilution assays, organoids were transduced with desired overexpression constructs and flow sorted to select for pure population. Counted number of organoids were plated in 20 μL matrigel and cultured in reduced growth factor media for 4-6 days. Images acquired under bright-field and analyzed for differentiation phenotype.

### COMET Experiments

cDNA encoding SOX9 (aa 229-509) was cloned into C-terminal end of nuclease dead Cas9 in pJB vector backbone to express under SFFV constitutive promoter. The pJB backbone contains a P2A-GFP cassette to enable identification of transfected cells. dCas9 alone or dCas9 fusion with SOX9^229-509^ were co-transfected with *PROM1*-targeting sgRNA library in HEK293T cells for 48 hours. Cells were maintained under sub-confluent conditions and re-transfected 72 hours later. After a total of 6 days, flow-sorted pure population of cells co-expressing dCas9 and sgRNA were utilized for ATAC seq, CUT&RUN and qPCR analyses. List of all sgRNA sequences is in Table S7 and primers used for PROM1 qRT-PCR are in Table S9.

### RNA isolation and qPCR

Total RNA was isolated using the RNeasy Mini Kit (QIAGEN), and cDNA was synthesized using iScript Reverse Transcription Supermix (Bio-Rad). RT-qPCR was performed using Power SYBR Green Master Mix (Thermo Fisher Scientific) on a CFX96 Real-Time PCR System (Bio-Rad). Primers were obtained from published studies or designed using Primer-BLAST and synthesized by Integrated DNA Technologies. Relative gene expression was normalized to ACTIN or GAPDH. Primer sequences are listed in Table S8.

### Western blotting

Cells or organoids were lysed in RIPA buffer supplemented with a protease inhibitor cocktail (Roche). Whole-cell extracts were resolved by SDS–polyacrylamide gel electrophoresis, transferred to polyvinylidene difluoride membranes, and probed with indicated primary antibodies. Bound antibodies were detected with horseradish peroxidase (HRP)–conjugated secondary antibodies and chemiluminescent HRP substrate. List of Primary antibodies utilized are listed in Table S8.

### Computational Methods

#### Raw sequencing data quality control and preprocessing

Raw sequencing reads from all genomic assays (ATAC-seq, CUT&RUN, and bulk RNA-seq) were assessed for quality using fastp v0.23.4^56^. Adapter sequences were automatically detected and trimmed, low-quality bases (Phred score < 20) were removed from read ends, and reads shorter than 36 bp after trimming were discarded. Per-library quality metrics including duplication rates, GC content, and Q20/Q30 base fractions were inspected before downstream alignment. All reads passing quality control were retained for alignment.

#### RNA-sequencing preprocessing and differential expression analysis

Total RNA was isolated using the RNeasy Mini Kit (Qiagen) and sequenced on a NovaSeq 6000 platform (Novogene Corporation Inc.) with poly-A capture, paired-end 150 bp reads (non-directional library preparation; ≥6-10 Gb per sample). Raw reads were quality-assessed using fastp v0.23.4^56^ and aligned to the human reference genome assembly GRCh38 using STAR v2.7.11a^57^, allowing only uniquely mapped reads. Gene-level read counts were quantified using RSEM v1.3.3^58^.

Differential expression (DE) analysis was performed using DESeq2 v1.44.0^59^. Raw counts were normalized by the DESeq2 median-of-ratios method. Genes with a false discovery rate (FDR) < 0.05 were considered differentially expressed. DE analyses were performed across the following comparisons: (i) SOX9 degradation (dTAGv-1 treated, day 4) versus DMSO control; (ii) SOX9 re-expression (drug release, day 6-13) versus degraded cells; (iii) individual BAF subunit knockdowns (shSMARCB1, shARID1A, shARID1A/B, shPBRM1, shBRD9) versus non-targeting control (shNTC); and (iv) SOX9 truncation mutants (SOX9^1-228^, SOX9^1-228-B1^) versus SOX9 full-length in normal colon organoids.

Gene set enrichment analysis (GSEA v4.3.3)^60^ was performed on pre-ranked gene lists ordered by the DESeq2 Wald statistic. Published gene signatures for intestinal stem cells (ISC), colon enterocytes, and stem cell programs were obtained from the Broad Institute MSigDB^60,61^ and from Haber et al. (2017)^62^, Munoz et al. (2012)^63^, Kazakevych et al. (2017)^64^, and Wang et al. (2020)^65^. A normalized enrichment score (NES) with FDR < 0.25 was considered significant, per GSEA recommended thresholds. Results were visualized as GSEA enrichment plots.

#### ATAC-seq data processing and chromatin accessibility analysis

ATAC-seq reads were aligned to the human reference genome GRCh38 using Bowtie2 v2.3.5^66^ with default paired-end parameters. Duplicate reads were identified and removed using Picard MarkDuplicates (Broad Institute). Reads mapping to mitochondrial chromosomes (chrM) and regions listed in the ENCODE blacklist were excluded. To assess library quality, fragment size distributions and transcription start site (TSS) enrichment scores were computed using deepTools v3^67^ and ATACseqQC^68^.

Peak calling was performed on deduplicated, blacklist-filtered BAM files using MACS2^69^ with parameters: *-f BAMPE -q 0.01 -g hs --keep-dup all*. Only peaks with a q-value < 1 × 10^−5^ were retained for downstream analyses. Replicate peak sets were merged to generate a consensus peak set using BEDTools v2.30.0^70^, retaining peaks present in at least two replicates. Normalized signal tracks (bigWig) were generated using deepTools bamCoverage with the following options: *--binSize 10 -- smoothLength 30 --normalizeUsing RPGC --effectiveGenomeSize 2730871774 --extendReads*. Genomic annotation of peaks (promoter versus distal enhancer) was performed using ChIPseeker v1.34.1^71^, defining promoters as regions within ±1000 bp of an annotated TSS (UCSC hg38 RefSeq). Peaks outside this window were classified as distal regulatory elements (putative enhancers).

Differential chromatin accessibility analysis between experimental conditions was performed using DiffBind v3.14.0^72^ with DESeq2 as the underlying statistical engine. Differentially accessible regions (DARs) were defined by |log^2^FC| > 1 and FDR < 0.05. The following pairwise comparisons were performed: (i) SOX9-degraded (dTAGv-1, day 4) versus DMSO control in HT29*SOX9-dTAG* cells; (ii) SOX9 re-expression (drug release day 6-13) versus day 4 degradation; (iii) individual BAF subunit knockdowns versus shNTC following SOX9 re-expression; and (iv) SOX9 truncation mutants (SOX9^1-228^, SOX9^1-303^, SOX9^1-379^) and SOX9^1-228-B1^ fusion versus full-length SOX9 in normal colon organoids. SOX9-dependent regulatory elements were operationally defined as loci exhibiting significant accessibility loss upon SOX9 degradation and restored accessibility upon drug release (rescue).

Heatmaps of accessibility signal and average profile plots were generated using deepTools computeMatrix (reference-point mode, ±1.5 kb or ±5 kb as indicated) and plotHeatmap/plotProfile. Read counts at peak regions were extracted using deepTools multiBamSummary. Genome browser tracks were visualized in IGV or UCSC Genome Browser.

#### CUT&RUN sequencing data processing and occupancy analysis

CUT&RUN paired-end reads were aligned to GRCh38 using Bowtie2 v2.3.5^66^ with parameters: *--local --very-sensitive --no-mixed --no-discordant --phred33 -I 10 -X 700*. *E. coli* spike-in reads were simultaneously aligned to the *E. coli* genome (strain K-12 MG1655, GenBank accession U00096.3) for normalization purposes. Spike-in normalization scaling factors were calculated based on the inverse proportion of reads mapping to the E. coli genome and were applied during bigWig generation for visualization, as well as to normalized read counts for quantitative comparisons across conditions. Peaks were first called on individual replicates and only reproducible peaks (present in ≥2 replicates) were retained before merging for visualization. Technical replicates (n = 2-4 per condition) were merged after alignment to maximize peak calling sensitivity. Reads mapping to the ENCODE blacklist were removed.

Peak calling for TF (SOX9, SMARCC1, SMARCB1) and histone marks (H3K4me1, H3K4me3, H3K27ac, H3K27me3) was performed using MACS2^69^. For TF CUT&RUN, narrow peak calling was used with parameters: *--format BAM --p 1e-5 --keep-dup all*. For broad histone marks (H3K4me1, H3K27me3), broad peak calling was employed (*--broad --broad-cutoff 0.1*). Consensus peak sets for each factor across replicates were generated using BEDTools intersect, retaining peaks reproducible in ≥2 replicates. Signal-to-noise quality was assessed by computing the fraction of reads in peaks (FRiP score); samples with FRiP < 0.01 were excluded.

Co-occupancy analysis between SOX9 and individual BAF subunits (SMARCC1, SMARCB1, ARID1A, PBRM1, BRD9) was performed by intersecting peak sets using BEDTools intersect with a minimum 1 bp overlap. The percentage of SOX9-bound sites co-occupied by each BAF subunit was calculated. To classify SOX9-co-bound sites according to BAF subcomplex identity (cBAF, ncBAF, pBAF), peaks were annotated based on the presence or absence of subunit-specific markers: cBAF sites were defined as peaks co-bound by SMARCB1 and ARID1A but not BRD9 or PBRM1; ncBAF sites by BRD9 occupancy without PBRM1 or SMARCB1/ARID1A; and pBAF sites by PBRM1 occupancy without BRD9. Sites positive for multiple subunit combinations were assigned to the all-BAF category. This classification was implemented in R using GenomicRanges v1.50.0^73^.

To assess the dependency of BAF occupancy on SOX9, CUT&RUN signal for SMARCC1 and SMARCB1 was quantified at SOX9-bound sites in DMSO control, SOX9-degraded (day 4), and drug-release conditions using deepTools multiBamSummary followed by plotHeatmap. Significant changes in BAF occupancy upon SOX9 loss were identified using DiffBind with the same thresholds described above. The reciprocal experiment—assessing SOX9 occupancy upon SMARCB1 knockdown—was similarly analyzed to determine whether SOX9 chromatin binding is independent of BAF.

#### Histone modification profiling and enhancer state classification

CUT&RUN signal for histone modifications (H3K4me1, H3K4me3, H3K27ac, H3K27me3) was quantified at BAF subcomplex-specific peak sets and SOX9-dependent regulatory elements. Poised enhancers were operationally defined as genomic loci with high H3K4me1 signal and low H3K4me3 and H3K27ac signal under basal conditions (DMSO), indicative of an inactive but primed chromatin state. Bivalent promoters were identified as promoter-annotated regions co-marked by H3K4me3 and H3K27me3. Histone modification heatmaps were generated with deepTools computeMatrix (±5 kb) and plotHeatmap. The histone state classification was cross-referenced with SOX9 and BAF co-occupancy data to define the subset of SOX9-cBAF co-bound poised enhancers associated with stem-like gene programs.

#### Single-cell RNA-sequencing analysis

Sequencing reads were demultiplexed, aligned to the mouse reference genome (mm10) or human GRCh38, and unique molecular identifier (UMI) counts were collapsed using Cell Ranger v7.1.0 (10x Genomics). Ambient RNA contamination was removed using CellBender v0.3.0^74^ prior to downstream processing. Doublets were identified and removed using DoubletFinder v2.0^75^.

Downstream quality control, normalization, dimensionality reduction, and clustering were performed using Seurat v4^76^ in RStudio or Scanpy^77^ in Python. Cells were retained if they had ≥100 and ≤25,000 unique features and <25% mitochondrial read fraction. Counts were log-normalized (NormalizeData, scale factor = 10,000) and the top 2,000 highly variable features were identified (FindVariableFeatures). Data were scaled and principal component analysis (PCA) was performed. The first 10 principal components were used for graph-based clustering (FindNeighbors, FindClusters; resolution = 0.3) and UMAP visualization (RunUMAP, dims = 1:10). Cell type identity was assigned based on established intestinal epithelial marker genes (e.g., *Lgr5*, *Ascl2* for stem cells; *Alpi*, *Fabp2* for enterocytes; *Muc2* for goblet cells; *Chga* for enteroendocrine cells).

For multi-sample integration (paired APC^WT^ and APC^KO^ intestinal epithelium across AOM/DSS, Cdx2^CreERT2^;Apc^f/f^;R26^tdT^, and Lgr5^CreERT2^;Apc^f/f^;R26^tdT^ mouse models), batch correction was performed using Seurat IntegrateData with the default CCA-based anchor integration method^76^. Following integration, PCA and UMAP were recomputed on the integrated object. Gene signature scoring was performed using AddModuleScore with published ISC, BAF complex, and spliceosome gene sets. Single-cell expression correlations with *Sox9* were computed using Pearson correlation on log-normalized expression values across all cells, and genes were ranked by correlation coefficient to identify co-expressed programs. Developmental potency was estimated using CytoTRACE^78^, which scores cells based on transcriptional diversity as a proxy for stemness.

#### Single-cell ATAC-sequencing analysis

scATAC-seq data from FACS-sorted tdTomato^+^ intestinal epithelium (Lgr5^CreERT2^;Apc^f/f^;R26^tdT^ mice) were processed using the ArchR v1.0.2 framework^79^. Reads were aligned to mm10 using the ArchR built-in alignment function (Bowtie2 v2.3.5, for paired-end reads). Arrow files were generated with a minimum TSS enrichment score of 4 and a minimum of 1,000 fragments per cell. Doublets were removed using the ArchR filterDoublets function. Dimensionality reduction was performed using iterative LSI (Latent Semantic Indexing), and UMAP embedding was computed on the LSI dimensions. Gene activity scores were computed using the ArchR GeneScoreMatrix. Chromatin accessibility scores at SOX9-cBAF co-bound enhancers and Sox9-dependent bulk ATAC peaks (from Bala et al., 2023^80^) were projected onto the scATAC-seq UMAP by summing signal over relevant peak sets using the ArchR addModuleScore function.

#### BAF subcomplex classification and subcomplex-specific occupancy

To distinguish sites occupied by cBAF, ncBAF, or pBAF subcomplexes, CUT&RUN peak sets for SMARCB1 (cBAF-specific subunit), BRD9 (ncBAF-specific), PBRM1 (pBAF-specific), ARID1A (cBAF), and SMARCC1 (shared core) were hierarchically intersected. For each genomic locus co-bound by SOX9 and SMARCC1, subcomplex identity was assigned as follows: (1) *cBAF:* co-occupied by SMARCB1 and ARID1A, negative for BRD9 and PBRM1; (2) *ncBAF:* co-occupied by BRD9, negative for SMARCB1 and PBRM1; (3) *pBAF:* co-occupied by PBRM1, negative for BRD9 and SMARCB1; (4) *all-BAF:* sites positive for multiple subcomplex markers. Loci exclusively bound by SOX9 and SMARCC1 without assignment to a specific subunit were analyzed as a separate category. Peak widths were compared across cBAF, ncBAF, pBAF, and all-BAF groups using Wilcoxon rank-sum tests to assess differences in site breadth at enhancers versus promoters.

#### Integrative analysis of chromatin accessibility and gene expression

To define the core SOX9-BAF regulatory network, we integrated ATAC-seq, CUT&RUN, and RNA-seq datasets. Regulatory elements were assigned to the nearest gene using ChIPseeker with a search window of ±1000 bp from the TSS. For each gene, the closest annotated SOX9-cBAF co-bound peak was identified. The core regulatory target set was defined as genes satisfying all of the following criteria: (i) presence of a SOX9-SMARCC1-SMARCB1 co-bound site within 100 kb; (ii) significant loss of chromatin accessibility at the associated peak upon SOX9 degradation (|log^2^FC| > 1, FDR < 0.05); (iii) significant downregulation in RNA-seq upon SOX9 degradation (|log^2^FC| > 0.5, FDR < 0.05). Pearson correlations between accessibility changes and expression changes across conditions were computed gene-wise and summarized as scatter plots. Wnt target gene enrichment at SOX9-regulated ATAC peaks in organoid ATAC-seq data was assessed using the published Ctnnb1-deletion downregulated gene list (Fevr et al., 2007^81^) as a reference set. Overlap significance was assessed by Fisher’s exact test.

For the SOX9 truncation mutant analysis, ATAC-seq signal at SOX9-regulated sites (n = 10,039) was extracted for each truncation (SOX9^1-228^, SOX9^1-303^, SOX9^1-379^, SOX9^FL^) and GFP control using deepTools multiBamSummary. Chromatin accessibility at these sites was compared across constructs to identify the minimal domain required for chromatin remodeling. Regions gaining accessibility upon introduction of SOX9^1-228-B1^ (SMARCB1 fusion) but not SOX9^1-228^ alone were identified by DiffBind (|log^2^FC| > 1, FDR < 0.05) and annotated for genomic feature type and stem cell gene proximity.

#### Rapid immunoprecipitation mass spectrometry of endogenous protein (RIME) analysis

RIME was performed to identify SOX9-interacting chromatin-associated proteins. Raw mass spectrometry data were analyzed using MaxQuant v2.0^82^ against the UniProt human proteome database with a 1% peptide and protein FDR threshold. Label-free quantification (LFQ) intensities were computed with a minimum of 2 unique peptides per protein. Proteins identified in the control IgG pulldown or absent in ≥1 SOX9 replicate were excluded. Enrichment over IgG control was assessed by log^2^ LFQ ratio (SOX9-IP/IgG). Functional pathway enrichment analysis of RIME-identified proteins was performed using STRING v12.0^83^ and Enrichr^84^ against GO Biological Process, CORUM protein complexes, and KEGG Pathway databases. Heatmap visualization of top-ranked interactors was generated in R using the ComplexHeatmap v2.14.0 package^85^.

#### Analysis of public datasets

TCGA CRC RNA-seq data (n= 522 tumors, n= 275 matched normals) were downloaded from the UCSC Xena browser (https://xenabrowser.net/) and used to assess relative expression of BAF complex subunits in CRC versus normal colon. Pairwise Pearson correlations between SOX9 and individual BAF subunit expression levels were computed on log^2^(RSEM+1) normalized values. Spatial transcriptomic data for human colorectal samples were obtained from the UCSC Xena single-cell browser and analyzed for co-expression of SOX9, SMARCC1, SMARCB1, ARID1A, PBRM1, BRD7, and BRD9 in tumor versus normal tissue regions. Public H3K27ac ChIP-seq data from colorectal cancer enhancer subtypes were downloaded from Orouji et al., 2022 (PMID: 34059508)^77^ and integrated with SOX9 CUT&RUN peaks to annotate regulatory elements. DepMap CRISPR-Cas9 loss-of-function gene effect scores for BAF subunits in CRC cell lines (n = 48) and CRC organoids (n = 10) were downloaded from the DepMap portal (https://depmap.org/portal/) and plotted as box plots stratified by subunit identity.

#### Data visualization

Genome-wide heatmaps and average signal profiles were generated using deepTools computeMatrix and plotHeatmap/plotProfile. Signal tracks were visualized in the Integrative Genomics Viewer (IGV v2.16^86^) and UCSC Genome Browser. Venn diagrams for peak overlap were generated using the R package VennDiagram v1.7.3. Scatter plots, box plots, and bar graphs were produced in R v4.3.1 using ggplot2 v3.4.4. PCA plots for bulk RNA-seq and ATAC-seq were generated using DESeq2 plotPCA and the prcomp function in R, respectively. UMAP embeddings were visualized using Seurat DimPlot and FeaturePlot functions. All heatmaps for gene expression were generated using ComplexHeatmap v2.14.0

#### Statistical analysis and reproducibility

Experiments were performed in biological triplicate unless otherwise stated. Data are presented as mean ± SD. For pairwise comparisons between two groups, an unpaired two-tailed Student’s *t*-test was used when data were normally distributed (Shapiro-Wilk test, *P* > 0.05); otherwise, a Wilcoxon rank-sum (Mann-Whitney U) test was applied. For multi-group comparisons, a Kruskal-Wallis test with Dunn’s post-hoc correction was used. Statistical significance thresholds are denoted as: \**P* < 0.05; \*\**P* < 0.01; \*\*\**P* < 0.001; \*\*\*\**P* < 0.0001; ns, not significant. For genomic analyses, differential thresholds of |log^2^FC| > 1 and FDR < 0.05 were used for ATAC-seq and CUT&RUN, and |log^2^FC| > 0.5 and FDR < 0.05 for RNA-seq unless otherwise stated. In all box plots, boxes span the interquartile range (IQR, 25th-75th percentile), the central line denotes the median, and whiskers extend to 1.5× IQR. Statistical analyses were performed in R v4.3.1, GraphPad Prism 9.0, and Microsoft Excel.

#### Computational pipeline for EasySci single nuclear RNA-seq processing

Fastq files were demultiplexed using Illumina ‘bcl2fastq’ software with settings from the EasySci GitHub (https://github.com/JunyueCaoLab/EasySci). Gene counts were generated following the same pipeline. The refdata-gex-GRCh38_and_GRCm39-2024-A was used as the reference genome. High-quality nuclei were retained based on the following criteria: (i) the number of UMIs within the range of 200 to 10,000; (ii) the number of detected genes within the range of 100 to 10,000; and (iii) the percentage of mitochondrial genes less than 20%. Scrublet was used to identify and remove potential doublets. Scanpy was used to process snRNA-seq data. Two thousand highly variable genes (HVGs) were selected by ranking the variance of 3,000 dispersion-based gene selections. PCR batch effects were corrected using “Harmony” with standard settings. Leiden clustering was performed, and cluster “0” was removed due to low UMI counts. The gene-by-nuclei count matrix was later exported to Seurat for subsequent CytoTrace2 analysis. A total of 47,673 nuclei were subjected to UMAP visualization and clustering using the first 20 dimensions. We then performed cellular potency calculations using CytoTrace2 with its standard settings. The relative CytoTrace scores were used for visualization.

#### Structure simulation and prediction for interacting residues

SOX9 and SMARCB1 structures were simulated using RosettaFold^3^0. To capture conformational variability arising from intrinsically disordered regions, the top five ranked models for SOX9 (509 aa) and SMARCB1 (385 aa) were used to generate structural ensembles. All 25 pairwise combinations of SOX9 and SMARCB1 ensemble members were subjected to rigid-body docking using ClusPro, which samples approximately 70,000 rotations per protein pair and evaluates roughly 10⁹–10¹⁰ conformations per run. The top 10 lowest-energy balanced-forces models from each run were retained, yielding a total of 250 docked complexes.

Inter - chain residue contacts were identified by calculating Cα - Cα Euclidean distances between all SOX9 and SMARCB1 residue pairs in each docked complex. Residue pairs with Cα distances ≤8 Å were considered interacting, consistent with established protein - protein interface definitions encompassing hydrogen bonding, electrostatic, and van der Waals interactions. For each residue, interaction frequency was calculated as the number of docked complexes in which it participated in at least one inter-chain contact, while run-level frequency was defined as the number of independent docking runs (out of 25) in which the residue was detected at the interface.

Evolutionary conservation scores for both proteins were obtained from ConSurf and normalized to [0,1]. A composite interaction score was computed per residue as the product of normalized run-level frequency and normalized ConSurf conservation score. Residues at or above the 90th percentile of the composite score distribution were classified as high-confidence interacting residues. Feature inter-correlation analysis confirmed that interaction frequency and conservation are orthogonal signals (R² = 0–0.01 across all partner pairs), justifying their multiplicative combination. All contact analysis and scoring were performed in R using the bio3d, dplyr, and ggplot2 packages.

To simulate structural model of SOX9-cBAF assembly, AlphaFold3-Multimer was used to perform sequential complexation modeling. Full-length human protein sequences of SOX9, SMARCC1, SMARCD1, ARID1A, and SMARCB1, together with the SOX9-bound DNA sequence, were submitted in stepwise increasing compositional complexity. For each complex, inter-chain contacts between SOX9 and each BAF subunit were identified using an 8 Å Cα–Cα distance threshold applied to the top-ranked AlphaFold3 model. Contact residues were mapped onto SOX9’s linear sequence and overlaid with per-residue PONDR VLXT disorder scores and ConSurf conservation scores to contextualize interfacial residues within functional and structural domains. Visualization of all structural models and interfacial residues was performed in PyMOL.

## Data availability

All data that support the findings of this study are available within the paper and its supplementary files. Raw and processed sequencing data are available in the Sequence Read Archive (SRA) at the NCBI Center with the accession number PRJNA1472915 and Gene Expression Omnibus (GEO) with the accession number GSE335210. All processed data are also available at https://doi.org/10.5281/zenodo.20447027. Source data are provided with this paper. All other data supporting the findings of this study are available from the corresponding author on reasonable request.

## Code availability

All bioinformatic analysis tools and pipelines used in this study are documented in the method section. Codes are available from the corresponding author upon reasonable request.

## Declaration of generative AI and AI-assisted technologies in the manuscript preparation process

During the preparation of this work, the authors used ChatGPT (OpenAI) to assist with language editing and revision of manuscript text. After using these tools, the authors reviewed and edited the content as needed and take full responsibility for the content of the published article.

## Author contributions

Conceptual: A.K.R., N.S.S.; Methodology: A.K.R., Z.L., X.W., Y.H., P.D., A.C.G, N.S.S.; Investigation: A.K.R., Z.L., X.W., Y.H., P.D., A.C.G, S.K., E.H., P.Y.P.L; Data analysis, statistics, and visualization: A.K.R., Z.L., X.W., Y.H., P.D., A.C.G, S.K., E.H.; Fund acquisition: J.M.C, N.S.S.; Supervision: N.S.S. Writing – original draft: A.K.R., Z.L., N.S.S; Writing – reviewing & editing: A.K.R., Z.L., X.W., Y.H., P.D., J.M.C., N.S.S.

## Acknowledgements

We thank Sethi lab members, including Siddhi Balamurali, Sarah Malafa, Xi Pu, Ananya Khosla, and Adrian Kalaw, for technical support; Glenn Dranoff, Jun Qi, Adam Sperling, Matt Oser, Ramesh Shivdasani, and others for insightful discussions and/or review of the manuscript; Dana-Farber/Harvard Cancer Center for the use of the Specialized Histopathology Core, which provided histology and immunohistochemistry service; Harvard Digestive Disease Center and NIH grant P30DK034854 for core services, resources, technology, and expertise. Dana-Farber/Harvard Cancer Center is supported in part by an NCI Cancer Center Support Grant # NIH 5 P30 CA06516. This work was funded by the Department of Defense (CA201084), Innovative Research Fund (DFCI), American Cancer Society (RSG-24-1321463, DBG-25-1500423), and NIH/NCI (1R01CA292507, 1R01CA325533) to N.S.S and philanthropic support from the Jimmy Fund Walk (Opiela Family), Craig Baskin, Dave & Carol Fischer, Dr. Diane T. Coulopoulos, Howard & Wendy Cox, and Peggy & Greg Strakosch to the Sethi Lab.

Supplementary Information is available for this paper.

Correspondence and requests for materials should be addressed to N.S.S.

## REFERENCES

1. Whyte, W.A., Orlando, D.A., Hnisz, D., Abraham, B.J., Lin, C.Y., Kagey, M.H., Rahl, P.B., Lee, T.I., and Young, R.A. (2013). Master transcription factors and mediator establish super-enhancers at key cell identity genes. Cell 153, 307–319. 10.1016/j.cell.2013.03.035.

2. Mashtalir, N., D’Avino, A.R., Michel, B.C., Luo, J., Pan, J., Otto, J.E., Zullow, H.J., McKenzie, Z.M., Kubiak, R.L., St Pierre, R., et al. (2018). Modular Organization and Assembly of SWI/SNF Family Chromatin Remodeling Complexes. Cell 175, 1272–1288 e1220. 10.1016/j.cell.2018.09.032.

3. Yan, Z., Cui, K., Murray, D.M., Ling, C., Xue, Y., Gerstein, A., Parsons, R., Zhao, K., and Wang, W. (2005). PBAF chromatin-remodeling complex requires a novel specificity subunit, BAF200, to regulate expression of selective interferon-responsive genes. Genes Dev 19, 1662–1667. 10.1101/gad.1323805.

4. Vierbuchen, T., Ling, E., Cowley, C.J., Couch, C.H., Wang, X., Harmin, D.A., Roberts, C.W.M., and Greenberg, M.E. (2017). AP-1 Transcription Factors and the BAF Complex Mediate Signal-Dependent Enhancer Selection. Mol Cell 68, 1067–1082 e1012. 10.1016/j.molcel.2017.11.026.

5. Boulay, G., Sandoval, G.J., Riggi, N., Iyer, S., Buisson, R., Naigles, B., Awad, M.E., Rengarajan, S., Volorio, A., McBride, M.J., et al. (2017). Cancer-Specific Retargeting of BAF Complexes by a Prion-like Domain. Cell 171, 163–178 e119. 10.1016/j.cell.2017.07.036.

6. Jain, S.U., Williamson, K.E., Ying, A.W., Turner, A.M., Jiang, R.J., Raval, S., So, K., Allison, M.J., Sankar, A., Same Guerra, D.D., et al. (2026). A SWI/SNF-specific Ig-like domain, SWIFT, is a transcription factor binding platform. Science 392, eaeb3627. 10.1126/science.aeb3627.

7. Jonason, A.S., Kunala, S., Price, G.J., Restifo, R.J., Spinelli, H.M., Persing, J.A., Leffell, D.J., Tarone, R.E., and Brash, D.E. (1996). Frequent clones of p53-mutated keratinocytes in normal human skin. Proc Natl Acad Sci U S A 93, 14025–14029. 10.1073/pnas.93.24.14025.

8. Martincorena, I., Roshan, A., Gerstung, M., Ellis, P., Van Loo, P., McLaren, S., Wedge, D.C., Fullam, A., Alexandrov, L.B., Tubio, J.M., et al. (2015). Tumor evolution. High burden and pervasive positive selection of somatic mutations in normal human skin. Science 348, 880–886. 10.1126/science.aaa6806.

9. Lee-Six, H., Olafsson, S., Ellis, P., Osborne, R.J., Sanders, M.A., Moore, L., Georgakopoulos, N., Torrente, F., Noorani, A., Goddard, M., et al. (2019). The landscape of somatic mutation in normal colorectal epithelial cells. Nature 574, 532–537. 10.1038/s41586-019-1672-7.

10. Li, R., Di, L., Li, J., Fan, W., Liu, Y., Guo, W., Liu, W., Liu, L., Li, Q., Chen, L., et al. (2021). A body map of somatic mutagenesis in morphologically normal human tissues. Nature 597, 398–403. 10.1038/s41586-021-03836-1.

11. Fowler, J.C., King, C., Bryant, C., Hall, M.W.J., Sood, R., Ong, S.H., Earp, E., Fernandez-Antoran, D., Koeppel, J., Dentro, S.C., et al. (2021). Selection of Oncogenic Mutant Clones in Normal Human Skin Varies with Body Site. Cancer Discov 11, 340–361. 10.1158/2159-8290.CD-20-1092.

12. Gupta, P.B., Pastushenko, I., Skibinski, A., Blanpain, C., and Kuperwasser, C. (2019). Phenotypic Plasticity: Driver of Cancer Initiation, Progression, and Therapy Resistance. Cell Stem Cell 24, 65–78. 10.1016/j.stem.2018.11.011.

13. Pennycuick, A., Teixeira, V.H., AbdulJabbar, K., Raza, S.E.A., Lund, T., Akarca, A.U., Rosenthal, R., Kalinke, L., Chandrasekharan, D.P., Pipinikas, C.P., et al. (2020). Immune Surveillance in Clinical Regression of Preinvasive Squamous Cell Lung Cancer. Cancer Discov 10, 1489–1499. 10.1158/2159-8290.CD-19-1366.

14. Jaiswal, S., Fontanillas, P., Flannick, J., Manning, A., Grauman, P.V., Mar, B.G., Lindsley, R.C., Mermel, C.H., Burtt, N., Chavez, A., et al. (2014). Age-related clonal hematopoiesis associated with adverse outcomes. N Engl J Med 371, 2488–2498. 10.1056/NEJMoa1408617.

15. Fearon, E.R., and Vogelstein, B. (1990). A genetic model for colorectal tumorigenesis. Cell 61, 759– 767. 10.1016/0092-8674(90)90186-i.

16. Kinzler, K.W., and Vogelstein, B. (1996). Lessons from hereditary colorectal cancer. Cell 87, 159–170. 10.1016/s0092-8674(00)81333-1.

17. Morin, P.J., Sparks, A.B., Korinek, V., Barker, N., Clevers, H., Vogelstein, B., and Kinzler, K.W. (1997). Activation of beta-catenin-Tcf signaling in colon cancer by mutations in beta-catenin or APC. Science 275, 1787–1790. 10.1126/science.275.5307.1787.

18. Korinek, V., Barker, N., Morin, P.J., van Wichen, D., de Weger, R., Kinzler, K.W., Vogelstein, B., and Clevers, H. (1997). Constitutive transcriptional activation by a beta-catenin-Tcf complex in APC-/-colon carcinoma. Science 275, 1784–1787. 10.1126/science.275.5307.1784.

19. Bala, P., Rennhack, J.P., Aitymbayev, D., Morris, C., Moyer, S.M., Duronio, G.N., Doan, P., Li, Z., Liang, X., Hornick, J.L., et al. (2023). Aberrant cell state plasticity mediated by developmental reprogramming precedes colorectal cancer initiation. Sci Adv 9, eadf0927. 10.1126/sciadv.adf0927.

20. Qin, X., Cardoso Rodriguez, F., Sufi, J., Vlckova, P., Claus, J., and Tape, C.J. (2023). An oncogenic phenoscape of colonic stem cell polarization. Cell 186, 5554–5568 e5518. 10.1016/j.cell.2023.11.004.

21. Vasquez, E.G., Nasreddin, N., Valbuena, G.N., Mulholland, E.J., Belnoue-Davis, H.L., Eggington, H.R., Schenck, R.O., Wouters, V.M., Wirapati, P., Gilroy, K., et al. (2022). Dynamic and adaptive cancer stem cell population admixture in colorectal neoplasia. Cell Stem Cell 29, 1213–1228 e1218. 10.1016/j.stem.2022.07.008.

22. Han, T., Goswami, S., Hu, Y., Tang, F., Zafra, M.P., Murphy, C., Cao, Z., Poirier, J.T., Khurana, E., Elemento, O., et al. (2020). Lineage Reversion Drives WNT Independence in Intestinal Cancer. Cancer Discov 10, 1590–1609. 10.1158/2159-8290.CD-19-1536.

23. Mzoughi, S., Schwarz, M., Wang, X., Demircioglu, D., Ulukaya, G., Mohammed, K., Zorgati, H., Torre, D., Tomalin, L.E., Di Tullio, F., et al. (2025). Oncofetal reprogramming drives phenotypic plasticity in WNT-dependent colorectal cancer. Nat Genet 57, 402–412. 10.1038/s41588-024-02058-1.

24. Bastide, P., Darido, C., Pannequin, J., Kist, R., Robine, S., Marty-Double, C., Bibeau, F., Scherer, G., Joubert, D., Hollande, F., et al. (2007). Sox9 regulates cell proliferation and is required for Paneth cell differentiation in the intestinal epithelium. J Cell Biol 178, 635–648. 10.1083/jcb.200704152.

25. Mori-Akiyama, Y., van den Born, M., van Es, J.H., Hamilton, S.R., Adams, H.P., Zhang, J., Clevers, H., and de Crombrugghe, B. (2007). SOX9 is required for the differentiation of paneth cells in the intestinal epithelium. Gastroenterology 133, 539–546. 10.1053/j.gastro.2007.05.020.

26. Liang, X., Duronio, G.N., Yang, Y., Bala, P., Hebbar, P., Spisak, S., Sahgal, P., Singh, H., Zhang, Y., Xie, Y., et al. (2022). An Enhancer-Driven Stem Cell-Like Program Mediated by SOX9 Blocks Intestinal Differentiation in Colorectal Cancer. Gastroenterology 162, 209–222. 10.1053/j.gastro.2021.09.044.

27. Chen, Q., Weng, K., Lin, M., Jiang, M., Fang, Y., Chung, S.S.W., Huang, X., Zhong, Q., Liu, Z., Huang, Z., et al. (2023). SOX9 Modulates the Transformation of Gastric Stem Cells Through Biased Symmetric Cell Division. Gastroenterology 164, 1119–1136 e1112. 10.1053/j.gastro.2023.01.037.

28. Spisak, S., Chen, D., Likasitwatanakul, P., Doan, P., Li, Z., Bala, P., Vizkeleti, L., Tisza, V., De Silva, P., Giannakis, M., et al. (2024). Identifying regulators of aberrant stem cell and differentiation activity in colorectal cancer using a dual endogenous reporter system. Nat Commun 15, 2230. 10.1038/s41467-024-46285-w.

29. Cardoso, E.C., Lee, H., England, F.J., Cho, H., Lu, R., Varankar, S.S., Park, M.S., Rekhtman, N., Koo, B.K., Simons, B.D., et al. (2026). Early fibrotic niches establish tumour-permissive microenvironments. Nature 653, 254–264. 10.1038/s41586-026-10399-6.

30. Skrupskelyte, G., Rojo Arias, J.E., Ajith, H., Dang, Y., Rossetti, D., Han, S., Tang, M.K.S., Bejar, M.T., Colom, B., Fowler, J.C., et al. (2026). Precancerous niche remodelling dictates nascent tumour persistence. Nature. 10.1038/s41586-026-10157-8.

31. Sethi, N.S., Kikuchi, O., Duronio, G.N., Stachler, M.D., McFarland, J.M., Ferrer-Luna, R., Zhang, Y., Bao, C., Bronson, R., Patil, D., et al. (2020). Early TP53 alterations engage environmental exposures to promote gastric premalignancy in an integrative mouse model. Nat Genet 52, 219–230. 10.1038/s41588-019-0574-9.

32. Mohammed, H., Taylor, C., Brown, G.D., Papachristou, E.K., Carroll, J.S., and D’Santos, C.S. (2016). Rapid immunoprecipitation mass spectrometry of endogenous proteins (RIME) for analysis of chromatin complexes. Nat Protoc 11, 316–326. 10.1038/nprot.2016.020.

33. Tebo, A.G., and Gautier, A. (2019). A split fluorescent reporter with rapid and reversible complementation. Nature Communications 10, 2822. 10.1038/s41467-019-10855-0.

34. Orlando, G., Law, P.J., Cornish, A.J., Dobbins, S.E., Chubb, D., Broderick, P., Litchfield, K., Hariri, F., Pastinen, T., Osborne, C.S., et al. (2018). Promoter capture Hi-C-based identification of recurrent noncoding mutations in colorectal cancer. Nat Genet 50, 1375–1380. 10.1038/s41588-018-0211-z.

35. Kang, M., Gulati, G.S., Brown, E.L., Qi, Z., Avagyan, S., Armenteros, J.J.A., Gleyzer, R., Zhang, W., Steen, C.B., D’Silva, J.P., et al. (2025). Improved reconstruction of single-cell developmental potential with CytoTRACE 2. Nat Methods 22, 2258–2263. 10.1038/s41592-025-02857-2.

36. Baek, M., DiMaio, F., Anishchenko, I., Dauparas, J., Ovchinnikov, S., Lee, G.R., Wang, J., Cong, Q., Kinch, L.N., Schaeffer, R.D., et al. (2021). Accurate prediction of protein structures and interactions using a three-track neural network. Science 373, 871–876. 10.1126/science.abj8754.

37. Comeau, S.R., Kozakov, D., Brenke, R., Shen, Y., Beglov, D., and Vajda, S. (2007). ClusPro: performance in CAPRI rounds 6-11 and the new server. Proteins 69, 781–785. 10.1002/prot.21795.

38. Patil, A., Strom, A.R., Paulo, J.A., Collings, C.K., Ruff, K.M., Shinn, M.K., Sankar, A., Cervantes, K.S., Wauer, T., St Laurent, J.D., et al. (2023). A disordered region controls cBAF activity via condensation and partner recruitment. Cell 186, 4936–4955 e4926. 10.1016/j.cell.2023.08.032.

39. Barisic, D., Chin, C.R., Meydan, C., Teater, M., Tsialta, I., Mlynarczyk, C., Chadburn, A., Wang, X., Sarkozy, M., Xia, M., et al. (2024). ARID1A orchestrates SWI/SNF-mediated sequential binding of transcription factors with ARID1A loss driving pre-memory B cell fate and lymphomagenesis. Cancer Cell 42, 583–604 e511. 10.1016/j.ccell.2024.02.010.

40. Yang, Y., Gomez, N., Infarinato, N., Adam, R.C., Sribour, M., Baek, I., Laurin, M., and Fuchs, E. (2023). The pioneer factor SOX9 competes for epigenetic factors to switch stem cell fates. Nat Cell Biol 25, 1185–1195. 10.1038/s41556-023-01184-y.

41. Nakayama, R.T., Pulice, J.L., Valencia, A.M., McBride, M.J., McKenzie, Z.M., Gillespie, M.A., Ku, W.L., Teng, M., Cui, K., Williams, R.T., et al. (2017). SMARCB1 is required for widespread BAF complex-mediated activation of enhancers and bivalent promoters. Nat Genet 49, 1613–1623. 10.1038/ng.3958.

42. Valencia, A.M., Collings, C.K., Dao, H.T., St Pierre, R., Cheng, Y.C., Huang, J., Sun, Z.Y., Seo, H.S., Mashtalir, N., Comstock, D.E., et al. (2019). Recurrent SMARCB1 Mutations Reveal a Nucleosome Acidic Patch Interaction Site That Potentiates mSWI/SNF Complex Chromatin Remodeling. Cell 179, 1342–1356 e1323. 10.1016/j.cell.2019.10.044.

43. Wang, X., Lee, R.S., Alver, B.H., Haswell, J.R., Wang, S., Mieczkowski, J., Drier, Y., Gillespie, S.M., Archer, T.C., Wu, J.N., et al. (2017). SMARCB1-mediated SWI/SNF complex function is essential for enhancer regulation. Nat Genet 49, 289–295. 10.1038/ng.3746.

44. Chan, Y.S., Gao, Q., Robinson, S.A., Wang, W., Filandrova, R., Weinhold, L.M., Cabrera, M.L., Zhang, M., Ambati, C.S.R., Lerario, A.M., et al. (2025). beta-catenin functions as a molecular adapter for disordered cBAF interactions. Mol Cell 85, 3041–3056 e3049. 10.1016/j.molcel.2025.06.026.

45. Huang, K.K., Ma, H., Chong, R.H.H., Uchihara, T., Lian, B.S.X., Zhu, F., Sheng, T., Srivastava, S., Tay, S.T., Sundar, R., et al. (2023). Spatiotemporal genomic profiling of intestinal metaplasia reveals clonal dynamics of gastric cancer progression. Cancer Cell 41, 2019–2037 e2018. 10.1016/j.ccell.2023.10.004.

46. Centonze, A., Roura, A.J., Novillo-Font, M., Giordano, C., Hernando-Momblona, X., Llanses, M., Prats, P., Sevillano, M., Cabot, D., Novell, M., et al. (2026). A Plastic EMP1+ to LGR5+ Cell State Conversion as a Bypass to KRASG12D Pharmacologic Inhibition in Metastatic Colorectal Cancer. Cancer Discov 16, 320–344. 10.1158/2159-8290.CD-25-0679.

47. Mathur, R., Alver, B.H., San Roman, A.K., Wilson, B.G., Wang, X., Agoston, A.T., Park, P.J., Shivdasani, R.A., and Roberts, C.W. (2017). ARID1A loss impairs enhancer-mediated gene regulation and drives colon cancer in mice. Nat Genet 49, 296–302. 10.1038/ng.3744.

48. Hiramatsu, Y., Fukuda, A., Ogawa, S., Goto, N., Ikuta, K., Tsuda, M., Matsumoto, Y., Kimura, Y., Yoshioka, T., Takada, Y., et al. (2019). Arid1a is essential for intestinal stem cells through Sox9 regulation. Proc Natl Acad Sci U S A 116, 1704–1713. 10.1073/pnas.1804858116.

49. Gilbert, L.A., Larson, M.H., Morsut, L., Liu, Z., Brar, G.A., Torres, S.E., Stern-Ginossar, N., Brandman, O., Whitehead, E.H., Doudna, J.A., et al. (2013). CRISPR-mediated modular RNA-guided regulation of transcription in eukaryotes. Cell 154, 442–451. 10.1016/j.cell.2013.06.044.

50. Simeonov, D.R., Gowen, B.G., Boontanrart, M., Roth, T.L., Gagnon, J.D., Mumbach, M.R., Satpathy, A.T., Lee, Y., Bray, N.L., Chan, A.Y., et al. (2017). Discovery of stimulation-responsive immune enhancers with CRISPR activation. Nature 549, 111–115. 10.1038/nature23875.

51. Hilton, I.B., D’Ippolito, A.M., Vockley, C.M., Thakore, P.I., Crawford, G.E., Reddy, T.E., and Gersbach, C.A. (2015). Epigenome editing by a CRISPR-Cas9-based acetyltransferase activates genes from promoters and enhancers. Nat Biotechnol 33, 510–517. 10.1038/nbt.3199.

52. Wang, X., Haswell, J.R., and Roberts, C.W. (2014). Molecular pathways: SWI/SNF (BAF) complexes are frequently mutated in cancer--mechanisms and potential therapeutic insights. Clin Cancer Res 20, 21–27. 10.1158/1078-0432.CCR-13-0280.

53. Kadoch, C., Hargreaves, D.C., Hodges, C., Elias, L., Ho, L., Ranish, J., and Crabtree, G.R. (2013). Proteomic and bioinformatic analysis of mammalian SWI/SNF complexes identifies extensive roles in human malignancy. Nat Genet 45, 592–601. 10.1038/ng.2628.

54. Duronio, G.N., Liang, X., Hebbar, P., Islam, M., Spisak, S., and Sethi, N.S. (2022). Truncating SOX9 Alterations Are Heterozygous Null Alleles in Genome-Stable Colorectal Cancer. Gastro Hep Adv 1, 709–713. 10.1016/j.gastha.2022.04.011.

55. Ji, D., Shao, C., Yu, J., Hou, Y., Gao, X., Wu, Y., Wang, L., and Chen, P. (2024). FOXA1 forms biomolecular condensates that unpack condensed chromatin to function as a pioneer factor. Mol Cell 84, 244–260 e247. 10.1016/j.molcel.2023.11.020.

56. Chen, S., Zhou, Y., Chen, Y., and Gu, J. (2018). fastp: an ultra-fast all-in-one FASTQ preprocessor. Bioinformatics 34, i884–i890. 10.1093/bioinformatics/bty560.

57. Dobin, A., Davis, C.A., Schlesinger, F., Drenkow, J., Zaleski, C., Jha, S., Batut, P., Chaisson, M., and Gingeras, T.R. (2013). STAR: ultrafast universal RNA-seq aligner. Bioinformatics 29, 15–21. 10.1093/bioinformatics/bts635.

58. Li, B., and Dewey, C.N. (2011). RSEM: accurate transcript quantification from RNA-Seq data with or without a reference genome. BMC Bioinformatics 12, 323. 10.1186/1471-2105-12-323.

59. Love, M.I., Huber, W., and Anders, S. (2014). Moderated estimation of fold change and dispersion for RNA-seq data with DESeq2. Genome Biol 15, 550. 10.1186/s13059-014-0550-8.

60. Subramanian, A., Tamayo, P., Mootha, V.K., Mukherjee, S., Ebert, B.L., Gillette, M.A., Paulovich, A., Pomeroy, S.L., Golub, T.R., Lander, E.S., and Mesirov, J.P. (2005). Gene set enrichment analysis: A knowledge-based approach for interpreting genome-wide expression profiles. Proceedings of the National Academy of Sciences 102, 15545–15550. doi:10.1073/pnas.0506580102.

61. Liberzon, A., Birger, C., Thorvaldsdóttir, H., Ghandi, M., Mesirov, Jill P., and Tamayo, P. (2015). The Molecular Signatures Database Hallmark Gene Set Collection. Cell Systems 1, 417–425. 10.1016/j.cels.2015.12.004.

62. Haber, A.L., Biton, M., Rogel, N., Herbst, R.H., Shekhar, K., Smillie, C., Burgin, G., Delorey, T.M., Howitt, M.R., Katz, Y., et al. (2017). A single-cell survey of the small intestinal epithelium. Nature 551, 333–339. 10.1038/nature24489.

63. Munoz, J., Stange, D.E., Schepers, A.G., van de Wetering, M., Koo, B.K., Itzkovitz, S., Volckmann, R., Kung, K.S., Koster, J., Radulescu, S., et al. (2012). The Lgr5 intestinal stem cell signature: robust expression of proposed quiescent ’+4’ cell markers. EMBO J 31, 3079–3091. 10.1038/emboj.2012.166.

64. Kazakevych, J., Sayols, S., Messner, B., Krienke, C., and Soshnikova, N. (2017). Dynamic changes in chromatin states during specification and differentiation of adult intestinal stem cells. Nucleic Acids Res 45, 5770–5784. 10.1093/nar/gkx167.

65. Wang, Y., Song, W., Wang, J., Wang, T., Xiong, X., Qi, Z., Fu, W., Yang, X., and Chen, Y.G. (2020). Single-cell transcriptome analysis reveals differential nutrient absorption functions in human intestine. J Exp Med 217. 10.1084/jem.20191130.

66. Langmead, B., and Salzberg, S.L. (2012). Fast gapped-read alignment with Bowtie 2. Nature Methods 9, 357–359. 10.1038/nmeth.1923.

67. Ramírez, F., Dündar, F., Diehl, S., Grüning, B.A., and Manke, T. (2014). deepTools: a flexible platform for exploring deep-sequencing data. Nucleic Acids Research 42, W187–W191. 10.1093/nar/gku365.

68. Ou, J., Liu, H., Yu, J., Kelliher, M.A., Castilla, L.H., Lawson, N.D., and Zhu, L.J. (2018). ATACseqQC: a Bioconductor package for post-alignment quality assessment of ATAC-seq data. BMC Genomics 19, 169. 10.1186/s12864-018-4559-3.

69. Zhang, Y., Liu, T., Meyer, C.A., Eeckhoute, J., Johnson, D.S., Bernstein, B.E., Nusbaum, C., Myers, R.M., Brown, M., Li, W., and Liu, X.S. (2008). Model-based Analysis of ChIP-Seq (MACS). Genome Biology 9, R137. 10.1186/gb-2008-9-9-r137.

70. Quinlan, A.R., and Hall, I.M. (2010). BEDTools: a flexible suite of utilities for comparing genomic features. Bioinformatics 26, 841–842. 10.1093/bioinformatics/btq033.

71. Yu, G., Wang, L.-G., and He, Q.-Y. (2015). ChIPseeker: an R/Bioconductor package for ChIP peak annotation, comparison and visualization. Bioinformatics 31, 2382–2383. 10.1093/bioinformatics/btv145.

72. Ross-Innes, C.S., Stark, R., Teschendorff, A.E., Holmes, K.A., Ali, H.R., Dunning, M.J., Brown, G.D., Gojis, O., Ellis, I.O., Green, A.R., et al. (2012). Differential oestrogen receptor binding is associated with clinical outcome in breast cancer. Nature 481, 389–393. 10.1038/nature10730.

73. Lawrence, M., Huber, W., Pagès, H., Aboyoun, P., Carlson, M., Gentleman, R., Morgan, M.T., and Carey, V.J. (2013). Software for Computing and Annotating Genomic Ranges. PLOS Computational Biology 9, e1003118. 10.1371/journal.pcbi.1003118.

74. Fleming, S.J., Chaffin, M.D., Arduini, A., Akkad, A.-D., Banks, E., Marioni, J.C., Philippakis, A.A., Ellinor, P.T., and Babadi, M. (2023). Unsupervised removal of systematic background noise from droplet-based single-cell experiments using CellBender. Nature Methods 20, 1323–1335. 10.1038/s41592-023-01943-7.

75. McGinnis, C.S., Murrow, L.M., and Gartner, Z.J. (2019). DoubletFinder: Doublet Detection in Single-Cell RNA Sequencing Data Using Artificial Nearest Neighbors. Cell Systems 8, 329–337.e324. 10.1016/j.cels.2019.03.003.

76. Hao, Y., Hao, S., Andersen-Nissen, E., Mauck, W.M., Zheng, S., Butler, A., Lee, M.J., Wilk, A.J., Darby, C., Zager, M., et al. (2021). Integrated analysis of multimodal single-cell data. Cell 184, 3573– 3587.e3529. 10.1016/j.cell.2021.04.048.

77. Orouji, E., Raman, A.T., Singh, A.K., Sorokin, A., Arslan, E., Ghosh, A.K., Schulz, J., Terranova, C., Jiang, S., Tang, M., et al. (2022). Chromatin state dynamics confers specific therapeutic strategies in enhancer subtypes of colorectal cancer. Gut 71, 938. 10.1136/gutjnl-2020-322835.

78. Gulati, G.S., Sikandar, S.S., Wesche, D.J., Manjunath, A., Bharadwaj, A., Berger, M.J., Ilagan, F., Kuo, Angera H., Hsieh, R.W., Cai, S., et al. (2020). Single-cell transcriptional diversity is a hallmark of developmental potential. Science 367, 405–411. doi:10.1126/science.aax0249.

79. Granja, J.M., Corces, M.R., Pierce, S.E., Bagdatli, S.T., Choudhry, H., Chang, H.Y., and Greenleaf, W.J. (2021). ArchR is a scalable software package for integrative single-cell chromatin accessibility analysis. Nature Genetics 53, 403–411. 10.1038/s41588-021-00790-6.

80. Bala, P., Rennhack, J.P., Aitymbayev, D., Morris, C., Moyer, S.M., Duronio, G.N., Doan, P., Li, Z., Liang, X., Hornick, J.L., et al. (2023). Aberrant cell state plasticity mediated by developmental reprogramming precedes colorectal cancer initiation. Science Advances 9, eadf0927. doi:10.1126/sciadv.adf0927.

81. Fevr, T., Robine, S., Louvard, D., and Huelsken, J. (2007). Wnt/beta-catenin is essential for intestinal homeostasis and maintenance of intestinal stem cells. Mol Cell Biol 27, 7551–7559. 10.1128/mcb.01034-07.

82. Cox, J., and Mann, M. (2008). MaxQuant enables high peptide identification rates, individualized p.p.b.-range mass accuracies and proteome-wide protein quantification. Nature Biotechnology 26, 1367–1372. 10.1038/nbt.1511.

83. Szklarczyk, D., Kirsch, R., Koutrouli, M., Nastou, K., Mehryary, F., Hachilif, R., Gable, A.L., Fang, T., Doncheva, N.T., Pyysalo, S., et al. (2023). The STRING database in 2023: protein-protein association networks and functional enrichment analyses for any sequenced genome of interest. Nucleic Acids Res 51, D638–d646. 10.1093/nar/gkac1000.

84. Chen, E.Y., Tan, C.M., Kou, Y., Duan, Q., Wang, Z., Meirelles, G.V., Clark, N.R., and Ma’ayan, A. (2013). Enrichr: interactive and collaborative HTML5 gene list enrichment analysis tool. BMC Bioinformatics 14, 128. 10.1186/1471-2105-14-128.

85. Gu, Z., Eils, R., and Schlesner, M. (2016). Complex heatmaps reveal patterns and correlations in multidimensional genomic data. Bioinformatics 32, 2847–2849. 10.1093/bioinformatics/btw313.

86. Robinson, J.T., Thorvaldsdóttir, H., Winckler, W., Guttman, M., Lander, E.S., Getz, G., and Mesirov, J.P. (2011). Integrative genomics viewer. Nat Biotechnol 29, 24–26. 10.1038/nbt.1754.

