## Supplementary material for "The SOX9 C-terminus encodes a transferable cBAF-recruitment module for enhancer remodeling and cell-state control": SUpplemetary Figures Methods and Tables

**The PDF file includes:**

Supplementary Figures S1-S7

Materials and Methods

References

Tables 1 to 9

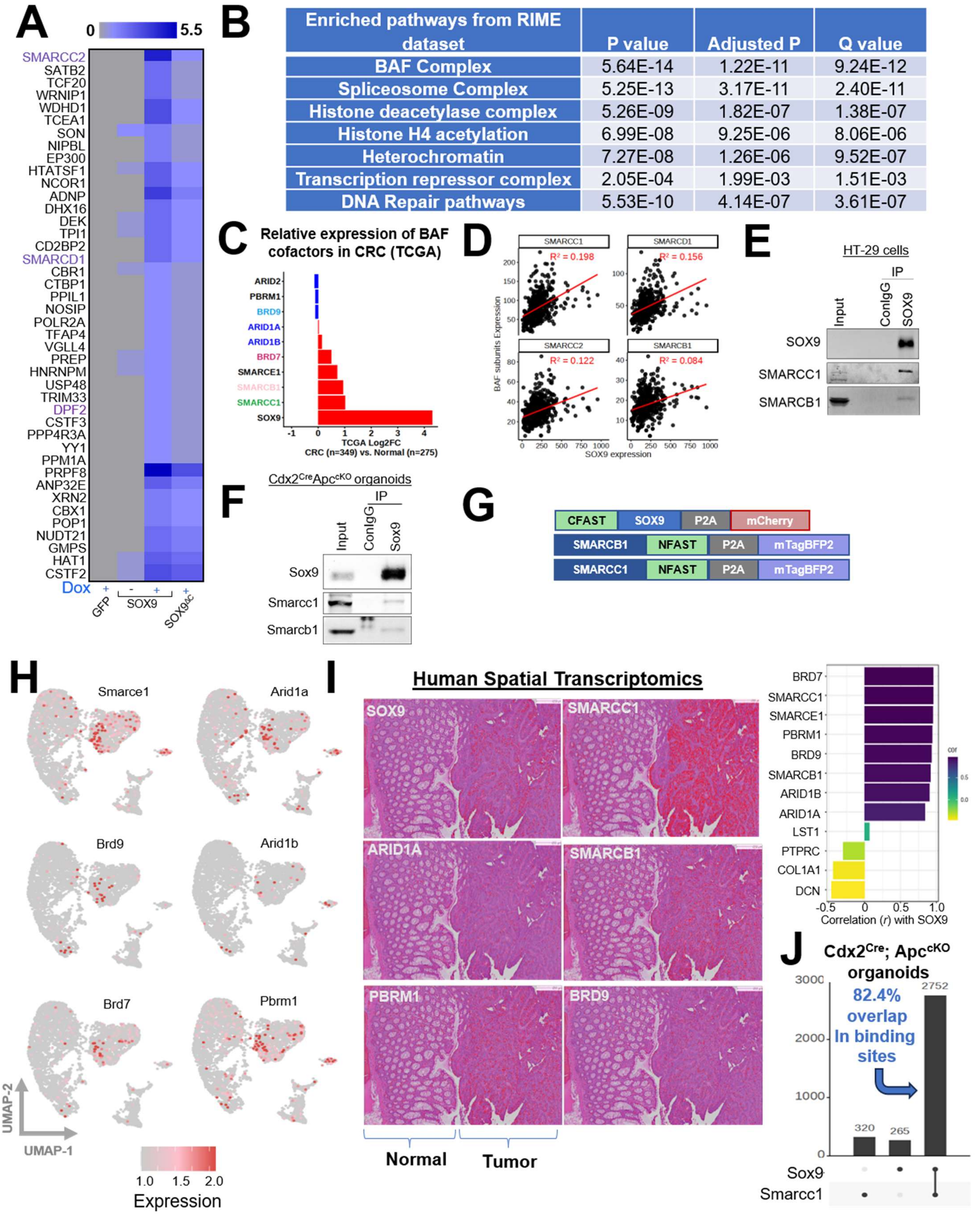

A
0
5.5
SMARCC2
SATB2
TCF20
WRNIP1
WDHD1
TCEA1
SON
NIPBL
EP300
HTATSF1
NCOR1
ADNP
DHX16
DEK
TPI1
CD2BP2
SMARCD1
CBR1
CTBP1
PPIL1
NOSIP
POLR2A
TFAP4
VGLL4
PREP
HNRNPM
USP48
TRIM33
DPF2
CSTF3
PPP4R3A
YY1
PPM1A
PRPF8
ANP32E
XRN2
CBX1
POP1
NUDT21
GMPS
HAT1
CSTF2
Dox
GFP
SOX9
SOX9ΔC
B
Enriched pathways from RIME dataset | P value | Adjusted P | Q value
BAF Complex | 5.64E-14 | 1.22E-11 | 9.24E-12
Spliceosome Complex | 5.25E-13 | 3.17E-11 | 2.40E-11
Histone deacetylase complex | 5.26E-09 | 1.82E-07 | 1.38E-07
Histone H4 acetylation | 6.99E-08 | 9.25E-06 | 8.06E-06
Heterochromatin | 7.27E-08 | 1.26E-06 | 9.52E-07
Transcription repressor complex | 2.05E-04 | 1.99E-03 | 1.51E-03
DNA Repair pathways | 5.53E-10 | 4.14E-07 | 3.61E-07
C
Relative expression of BAF cofactors in CRC (TCGA)
ARID2
PBRM1
BRD9
ARID1A
ARID1B
BRD7
SMARCE1
SMARCB1
SMARCC1
SOX9
TCGA Log2FC
CRC (n=349) vs. Normal (n=275)
D
SMARCC1
SMARCD1
SMARCC2
SMARCB1
R² = 0.198
R² = 0.156
R² = 0.122
R² = 0.084
BAF subunits Expression
SOX9 expression
E
HT-29 cells
IP
Input
ConIgG
SOX9
SOX9
SMARCC1
SMARCB1
F
Cdx2CreApccKO organoids
IP
Input
ConIgG
Sox9
Sox9
Smarcc1
Smarcb1
G
CFAST
SOX9
P2A
mCherry
SMARCB1
NFAST
P2A
mTagBFP2
SMARCC1
NFAST
P2A
mTagBFP2
H
Smarce1
Arid1a
Brd9
Arid1b
Brd7
Pbrm1
UMAP-2
UMAP-1
1.0
1.5
2.0
Expression
I
Human Spatial Transcriptomics
SOX9
SMARCC1
ARID1A
SMARCB1
PBRM1
BRD9
Normal
Tumor
BRD7
SMARCC1
SMARCE1
PBRM1
BRD9
SMARCB1
ARID1B
ARID1A
LST1
PTPRC
COL1A1
DCN
Correlation (r) with SOX9
J
Cdx2Cre; ApccKO organoids
82.4% overlap In binding sites
2752
320
265
Sox9
Smarcc1

**Figure S1. SOX9 associates with BAF cofactors across premalignant and CRC contexts**

**(A)** Heatmap of proteins detected by RIME in HT-115 CRC cells expressing V5-tagged GFP, SOX9 +/- dox induction, or $SOX9^{\Delta C}$; list of SOX9-interacting proteins detected by RIME are in Table 1)

**(B)** Chart shows P-values, adjusted P-values and Q-values obtained upon GENETS analysis for enrichment of pathways among SOX9-C terminal interactome,

**(C)** Ranked expression of SOX9 and BAF components in CRC tumors (n=349) compared to normal (n=275) in TCGA dataset.

**(D)** Correlation of SOX9 expression with genes encoding BAF components SMARCC1, SMARCC2, SMARCD1 and SMARCB1 across TCGA CRC tumors; correlation $R^2$ displayed for each protein in red.

**(E)** Immunoprecipitation of endogenous SOX9 in HT-29 CRC cells, showing complex formation between SOX9, SMARCC1 and SMARCB1.

**(F)** Immunoprecipitation of endogenous Sox9 in lysates of $Cdx2^{Cre}Apc^{cKO}$ colon organoids, showing interaction between Sox9, Smarcc1, Smarcd1, Smarcb1 and Dpf2.

**(G)** Schematic shows design of expression constructs used for Split-FAST assay.

**(H)** UMAPs of scRNA-seq data from $Lgr5^{CreERT2}$;$R26^{tdT}$ and $Lgr5^{CreERT2}$;$Apc^{f/f}$;$R26^{tdT}$ mice (Bala et al., 2023) showing expression of Smarce1, Arid1a, Brd9 and Arid1b.

**(I)** Spatial transcriptomics analysis shows correlated upregulation of SOX9 and indicated BAF components in Visium HD images, scale bars are 259µm (Left); associated quantification is shown in bar graph (right).

**(J)** Quantification of Cut&Run performed in $Cdx2^{Cre}$; $Apc^{cKO}$ organoids shows co-binding of chromatin by Sox9 and Smarcc1.

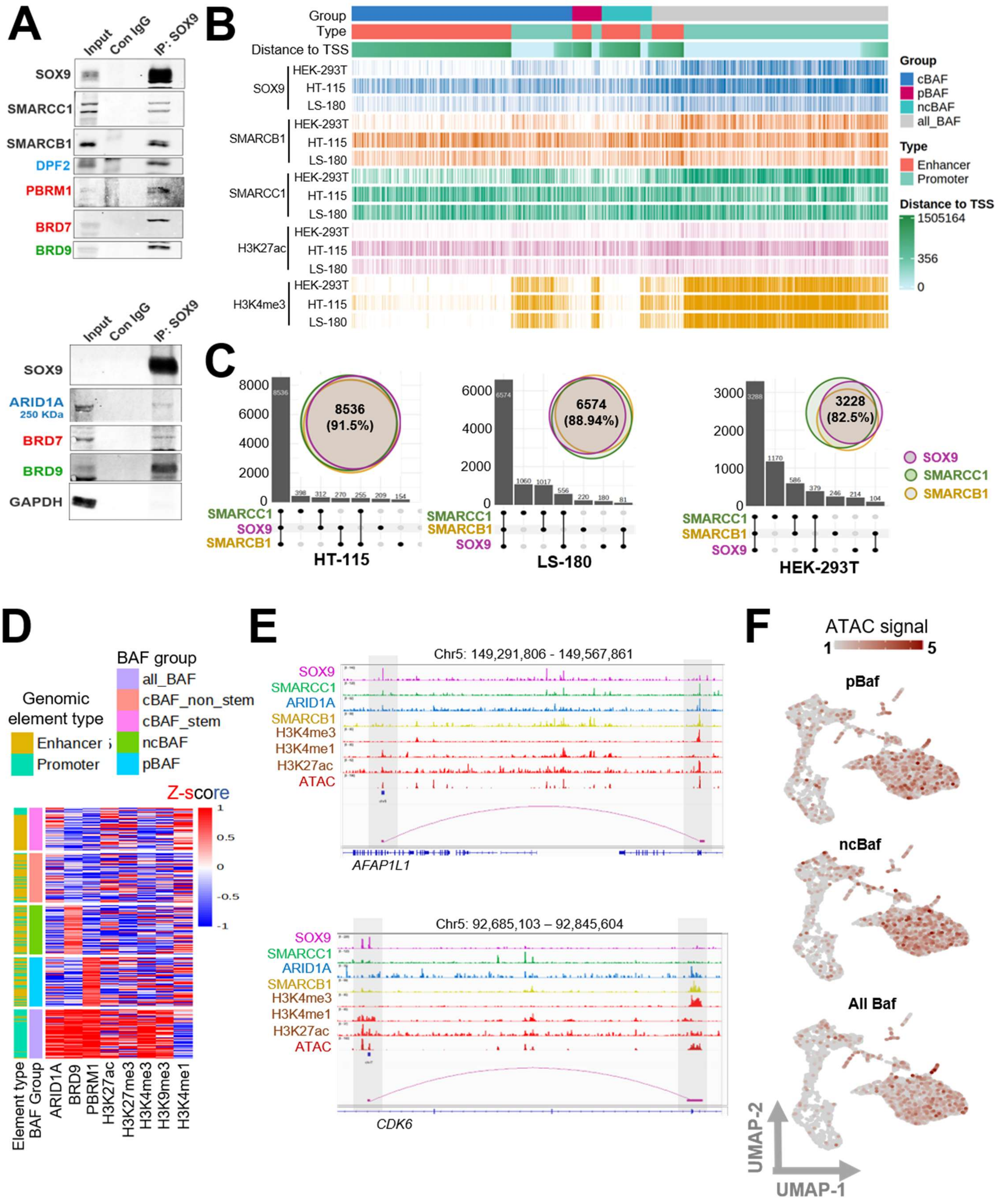

A
Input
Con IgG
IP: SOX9
SOX9
SMARCC1
SMARCB1
DPF2
PBRM1
BRD7
BRD9
ARID1A
250 KDa
GAPDH
B
Group
Type
Distance to TSS
HEK-293T
HT-115
LS-180
SOX9
SMARCB1
SMARCC1
H3K27ac
H3K4me3
Group
cBAF
pBAF
ncBAF
all_BAF
Type
Enhancer
Promoter
Distance to TSS
1505164
356
0
C
8536
(91.5%)
6574
(88.94%)
3228
(82.5%)
HT-115
LS-180
HEK-293T
SOX9
SMARCC1
SMARCB1
D
Genomic element type
Enhancer
Promoter
BAF group
all_BAF
cBAF_non_stem
cBAF_stem
ncBAF
pBAF
Z-score
Element type
BAF Group
ARID1A
BRD9
PBRM1
H3K27ac
H3K27me3
H3K4me3
H3K9me3
H3K4me1
E
Chr5: 149,291,806 - 149,567,861
SOX9
SMARCC1
ARID1A
SMARCB1
H3K4me3
H3K4me1
H3K27ac
ATAC
AFAP1L1
Chr5: 92,685,103 – 92,845,604
CDK6
F
ATAC signal
1
5
pBaf
ncBaf
All Baf
UMAP-2
UMAP-1

**Figure S2. SOX9 partitions BAF assemblies across distinct regulatory contexts**

**(A)** Western blot analysis of SOX9-immunoprecipitates from lysates of HT-29 cells showing cellular complex formation between SOX9 and representative core BAF (SMARCC1), cBAF (ARID1A and DPF2), pBAF (PBRM1 and BRD7), and ncBAF (BRD9) assemblies.

**(B)** Annotation tracks showing gene set association, distance to transcription start site, histone modification state, genomic element type, and BAF assembly class for SOX9-BAF co-bound loci in HT-115, LS-180 and HEK293T cells.

**(C)** Quantification of Cut&Run performed in HT-115, LS-180 and HEK293T cells shows co-binding of chromatin by SOX9, SMARCC1 and SMARCB1.

**(D)** Annotation tracks showing association between BAF assembly class, histone modification states, genomic element type, for SOX9-BAF co-bound loci.

**(E)** IGV plots visualize chromatin loci of *AFAP1L1* and *CDK8* in Hi-C analysis; associated gene list is in Table 3.

**(F)** Single-cell ATAC-seq of FACS-sorted tdT+ intestinal epithelium from $Lgr5^{CreERT2}$;$R26^{tdT}$ and $Lgr5^{CreERT2}$;$Apc^{f/f}$;$R26^{tdT}$ mice, showing UMAPs by SOX9-pBAF, SOX9-ncBAF and SOX9-All BAF co-bound enhancer accessibility.

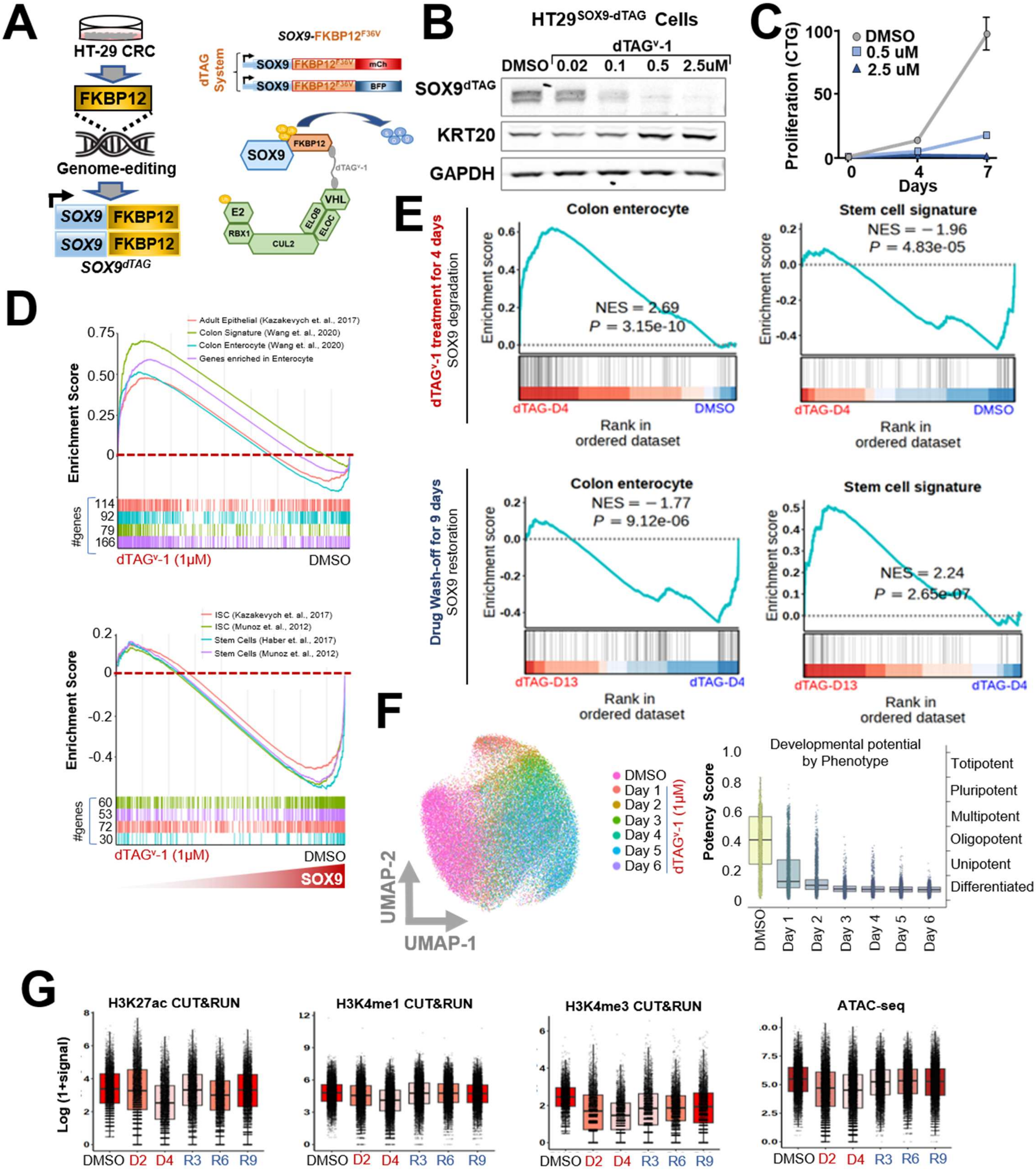

A
HT-29 CRC
FKBP12
Genome-editing
SOX9 FKBP12
SOX9 FKBP12
SOX9dTAG
SOX9-FKBP12F36V
dTAG System
SOX9 FKBP12F36V mCh
SOX9 FKBP12F36V BFP
SOX9
FKBP12
dTAGv-1
VHL
E2
RBX1
ELOB
ELOC
CUL2
B
HT29SOX9-dTAG Cells
dTAGv-1
DMSO 0.02 0.1 0.5 2.5uM
SOX9dTAG
KRT20
GAPDH
C
Proliferation (CTG)
DMSO
0.5 uM
2.5 uM
Days
D
Adult Epithelial (Kazakevych et. al., 2017)
Colon Signature (Wang et. al., 2020)
Colon Enterocyte (Wang et. al., 2020)
Genes enriched in Enterocyte
Enrichment Score
#genes
dTAGv-1 (1μM)
DMSO
ISC (Kazakevych et. al., 2017)
ISC (Munoz et. al., 2012)
Stem Cells (Haber et. al., 2017)
Stem Cells (Munoz et. al., 2012)
SOX9
E
dTAGv-1 treatment for 4 days
SOX9 degradation
Colon enterocyte
NES = 2.69
P = 3.15e-10
Stem cell signature
NES = −1.96
P = 4.83e-05
Enrichment score
dTAG-D4
DMSO
Rank in ordered dataset
Drug Wash-off for 9 days
SOX9 restoration
Colon enterocyte
NES = −1.77
P = 9.12e-06
Stem cell signature
NES = 2.24
P = 2.65e-07
dTAG-D13
dTAG-D4
F
DMSO
Day 1
Day 2
Day 3
Day 4
Day 5
Day 6
dTAGv-1 (1μM)
UMAP-1
UMAP-2
Developmental potential by Phenotype
Potency Score
Totipotent
Pluripotent
Multipotent
Oligopotent
Unipotent
Differentiated
G
H3K27ac CUT&RUN
H3K4me1 CUT&RUN
H3K4me3 CUT&RUN
ATAC-seq
Log (1+signal)
DMSO D2 D4 R3 R6 R9

**Figure S3. Endogenous SOX9 reversibly controls stem-like transcriptional and chromatin states**

**(A)** Schematic of biallelic *FKBP12*$^{F36V}$ (dTAG) degron knock-in at endogenous *SOX9* locus of HT-29 CRC cells. Schematic (right) of *FKBP12*$^{F36V}$ (dTAG) degron construct used to generate SOX9$^{dTAG}$; Schematic shows dTAG$^{v}$-1 mediated recruitment of E3 Ub-ligase and proteasomal machinery that degrades SOX9$^{dTAG}$.

**(B)** Immunoblot shows dose dependent degradation of SOX9$^{dTAG}$ and concurrent induction of KRT20 in HT-29$^{SOX9\text{-}dTAG}$ cells upon treatment with dTAG$^{v}$-1.

**(C)** Cell proliferation analysis by CTG assay in dose and time-course treatment of HT-29$^{SOX9\text{-}dTAG}$ cells with dTAG$^{v}$-1.

**(D)** GSEA of bulk mRNA-seq of dTAG$^{v}$-1 (1μM) treated HT-29$^{SOX9\text{-}dTAG}$ cells. Increased differentiation signatures (top) and Stem-like signatures (below) are shown; Normalized enrichment scores (NES) for each signature is listed; associated genes are listed in Table 2 and P-values and NES are listed in Table 4.

**(E)** GSEA of bulk mRNA-seq of HT-29$^{SOX9\text{-}dTAG}$ cells after 4 days of dTAG$^{v}$-1 treatment (sample D4; SOX9 degradation) and 9 days of drug wash-off (sample D13; SOX9 restoration). Panels show Increased colon enterocyte / differentiation signatures (top-left) and stem-like signatures (top-right) upon SOX9 degradation. Bottom panels show return of stem-like signatures upon SOX9 restoration. Normalized enrichment scores (NES) and P values for each signature are indicated.

**(F)** UMAP shows CytoTrace-based potency visualization from single-nucleus RNA-seq data of HT-29$^{SOX9\text{-}dTAG}$ cells at different timepoints dTAG$^{v}$-1 treatment with (top); Box plot shows developmental potential of dTAG$^{v}$-1 (1μM) treated HT-29$^{SOX9\text{-}dTAG}$ cells at each day of treatment.

**(G)** Box plots show CUT&RUN signals from H3k4me1- , H3k4me3- and H3K27ac- and ATAC signals (right) in HT-29$^{SOX9\text{-}dTAG}$ cells at different timepoints of dTAG$^{v}$-1 treatment (D2 and D4) and release (R3, R6 and R9). P-values are listed in Table 5.

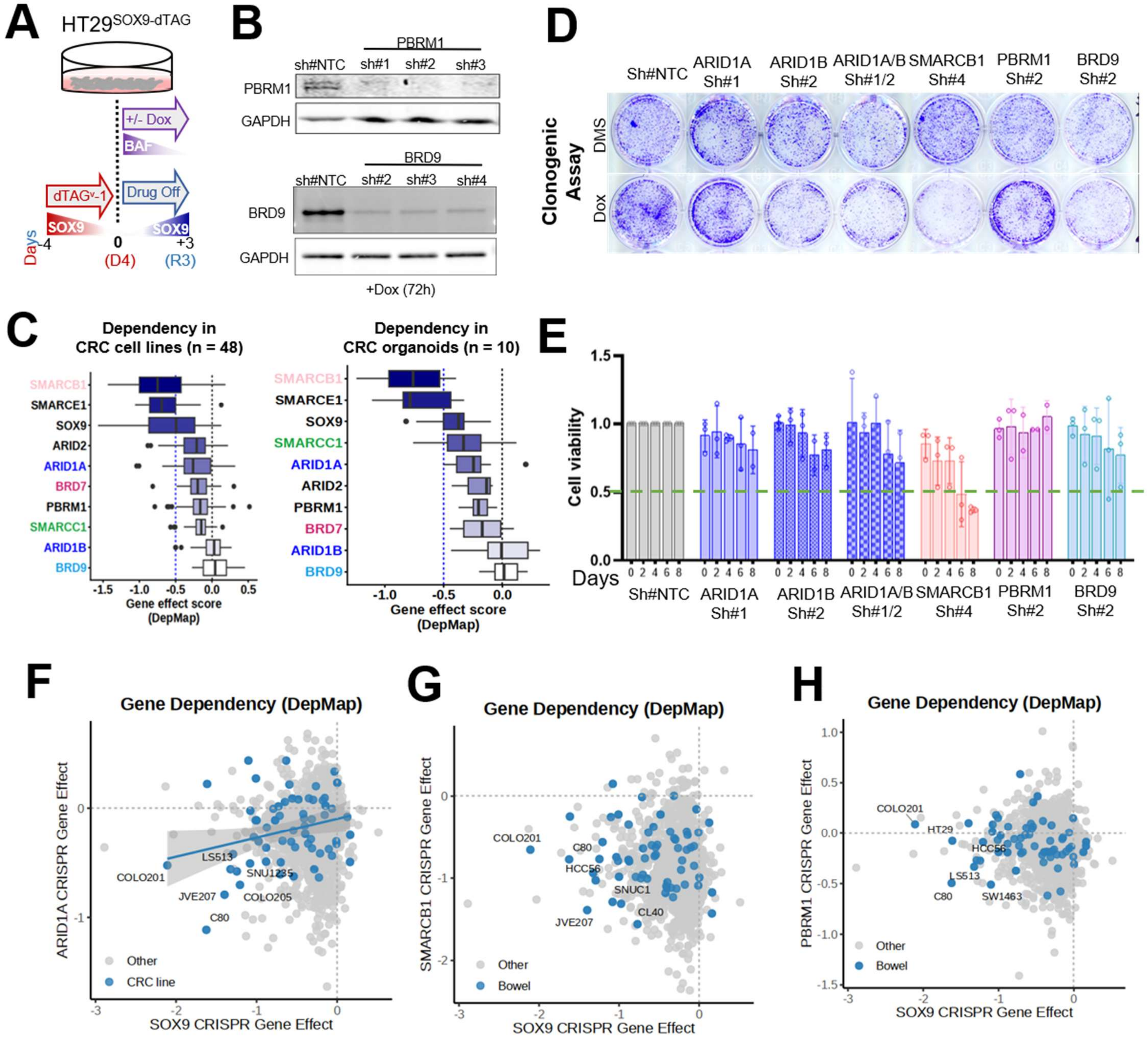

**Figure S4. CRC models show selective dependence on the SOX9-SMARCB1/cBAF axis**

**(A)** Schematic of the experimental design combining reversible SOX9 restoration with inducible shRNA-mediated suppression of individual BAF subunits.

**(B)** Immunoblot shows efficacy of dox-inducible shRNAs targeting PBRM1 and BRD9 in HT-29$^{SOX9\text{-}dTAG}$ cells.

**(C)** Dependency scores for SOX9 and BAF complex components across CRC cell lines and patient-derived CRC organoid models.

**(D)** Images show crystal violet assay performed on HT-29$^{SOX9\text{-}dTAG}$ cells with BAF subunit knockdown.

**(E)** Bar graph showing cell proliferation analysis by CTG assay in HT-29 cells upon BAF subunit knockdown.

**(F)** Graph shows co-dependency of colorectal cancer cell lines on SOX9 and ARID1A; Pearson coefficient (r) and P-values are displayed in red.

**(G)** Graph shows co-dependency of colorectal cancer cell lines on SOX9 and SMARCB1; Pearson coefficient (r) and P-values are displayed in red.

**(H)** Graph shows co-dependency of colorectal cancer cell lines on SOX9 and PBRM1.

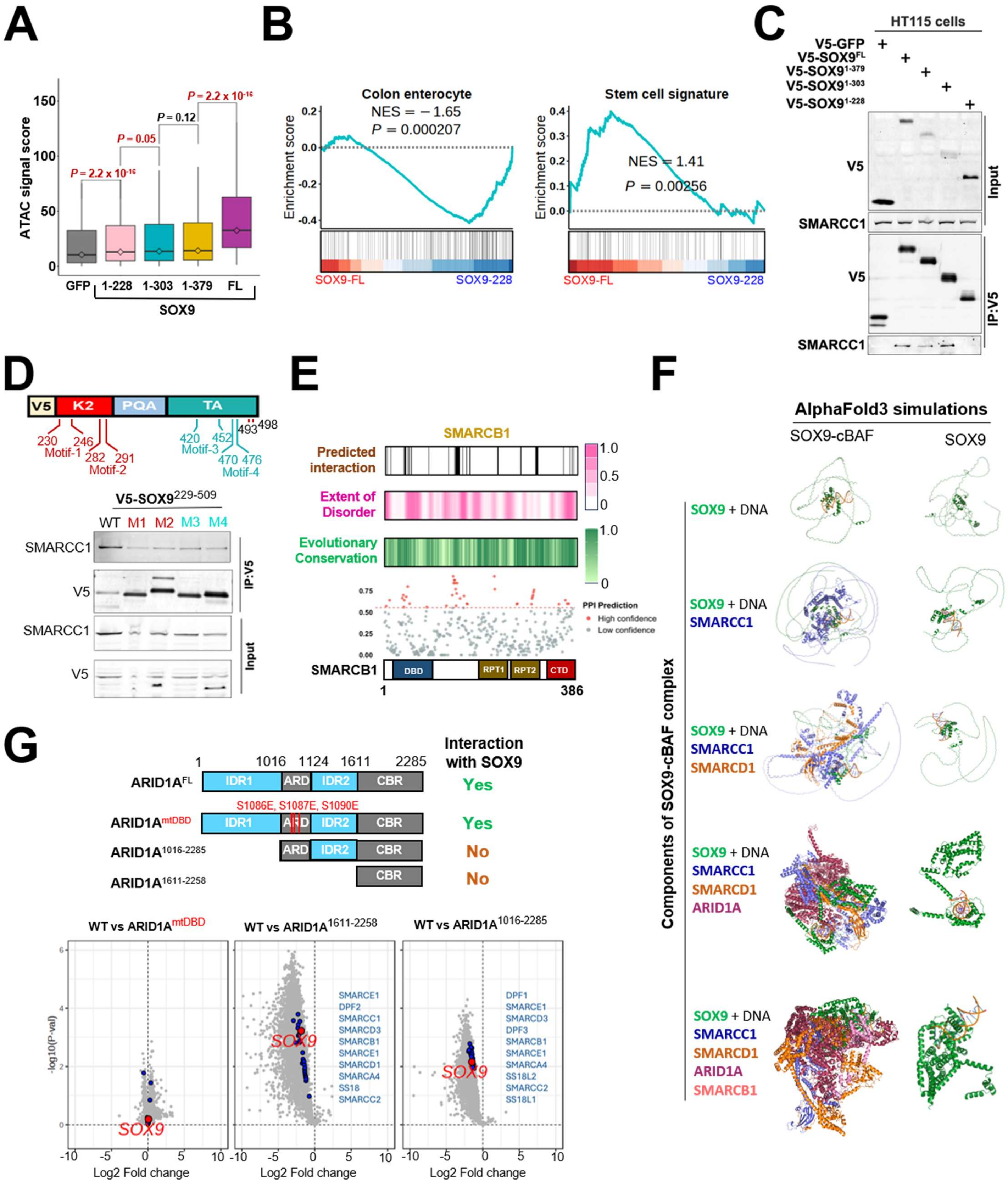

**Figure S5. The SOX9 C-terminal IDR contains a distributed cBAF-engagement surface**

**(A)** Bar graph shows ATAC signal scores chromatin accessibility after expression of GFP, $SOX9^{FL}$ (full-length) and SOX9 C-terminal truncation variants.

**(B)** GSEA shows enrichment of differentiation genes (left) and suppression of stem cell genes (right) upon expression of $SOX9^{1-228}$; NES and P values are indicated on respective graphs.

**(C)** V5-immunoprecipitation followed by immunoblotting for endogenous SMARCC1 and V5, performed on lysates of HT-115 cells overexpressing V5-tagged GFP, SOX9, or its C-terminal deletion variants.

**(D)** Schematic of SOX9 K2 and TA domain residues predicted to interact with SMARCC1 (top) and V5-immunoprecipitation followed by immunoblotting for V5, and SMARCC1 in cells expressing GFP, $SOX9^{229\text{-}509}$, or alanine substituted variants (below).

**(E)** Schematic of SMARCB1 shows residue level conservation, extent of disorder and predicted residues participating in SOX9-interaction.

**(F)** AlphaFold3 simulations for a SOX9-cBAF assembly; included cBAF components and simulated complex shown (top row); SOX9 conformations are extracted and displayed (below).

**(G)** Re-analysis of MS data-set from (Patil et. al., 2023); Schematic shows domain structure of ARID1A (top) and volcano plot shows interacting proteins (below)

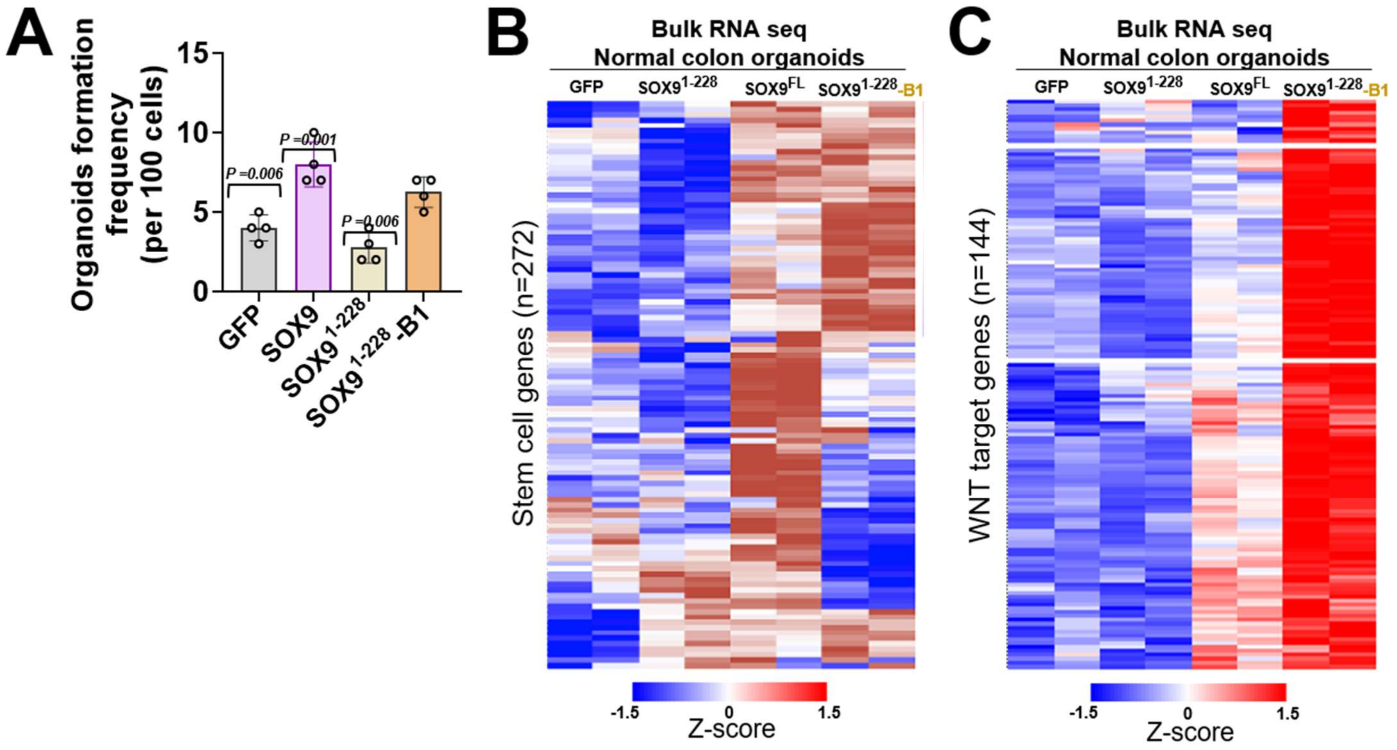

**Figure S6. Enforced SMARCB1 docking partially restores stem-like transcription and self-renewal to SOX9$^{\Delta C}$**

**(A)** Bar graph shows results from limiting dilution assay (LDA) as total number of organoids formed per 100 cells; this is a representative result from 3 independent experiments.

**(B)** Bulk RNA-seq heatmap of WNT target gene expression across normal colon organoids expressing GFP, SOX9$^{FL}$, SOX9$^{1\text{-}228}$, SOX9$^{1\text{-}228}$-SMARCB1

**(C)** Bulk RNA-seq heatmap of stem cell genes expressed across normal colon organoids expressing GFP, SOX9$^{FL}$, SOX9$^{1\text{-}228}$, SOX9$^{1\text{-}228}$-SMARCB1

A

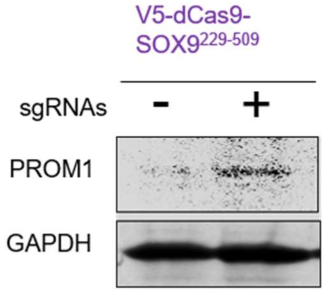

V5-dCas9-
SOX9229-509
sgRNAs
-
+
PROM1
GAPDH

B

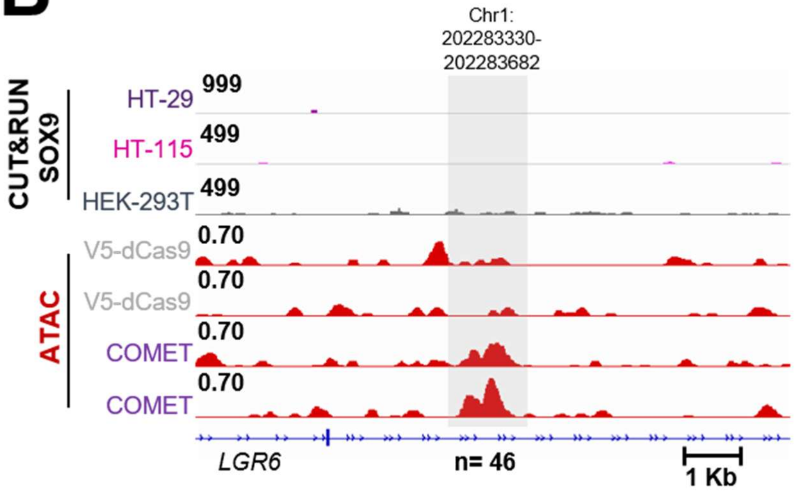

Chr1:
202283330-
202283682
CUT&RUN
SOX9
HT-29
999
HT-115
499
HEK-293T
499
ATAC
V5-dCas9
0.70
V5-dCas9
0.70
COMET
0.70
COMET
0.70
LGR6
n= 46
1 Kb

C

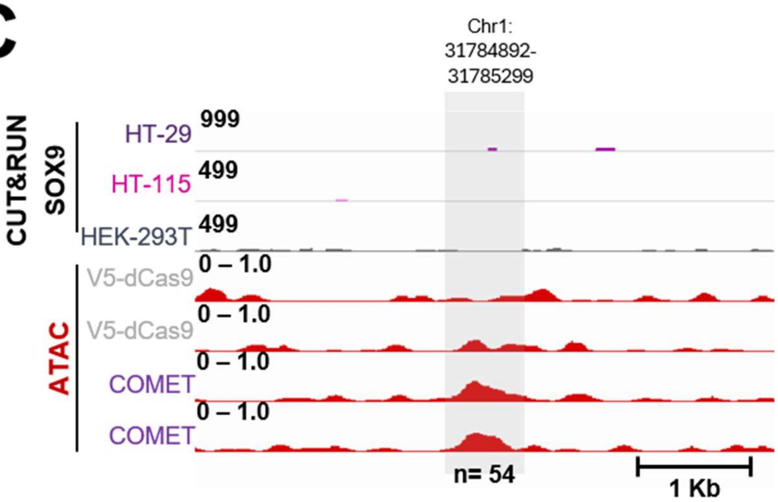

Chr1:
31784892-
31785299
CUT&RUN
SOX9
HT-29
999
HT-115
499
HEK-293T
499
ATAC
V5-dCas9
0 – 1.0
V5-dCas9
0 – 1.0
COMET
0 – 1.0
COMET
0 – 1.0
n= 54
1 Kb

**Figure S7. COMET extends SOX9 C-terminal chromatin-opening activity to non-native loci**

**(A)** Western blots shows expression of PROM1 in HEK293T cells upon enhancer activation by COMET.

**(B)** Genome browser views of CUT&RUN signals (SOX9 in HT29, HT115 and HEK293T cells), and ATAC-seq signal from HEK293T cells after targeting dCas9 or dCas9-SOX9$^{229\text{-}509}$ with pooled sgRNAs targeting LGR6 locus.

**(C)** Genome browser views of CUT&RUN signals (SOX9 in HT29, HT115 and HEK293T cells), and ATAC-seq signal from HEK293T cells after targeting dCas9 or dCas9-SOX9$^{229\text{-}509}$ with pooled sgRNAs targeting locus Chr1: 31784892-5299.

**Flow analysis**

Intestines from euthanized mice were washed with ice-cold PBS, villi were removed by scraping, and epithelial cells were dissociated in 5mM EDTA/PBS at 4°C for 20 minutes with intermittent shaking. The epithelial fraction was collected by centrifugation, incubated in prewarmed 4× TrypLE at 37°C for 30 minutes, filtered through a 70μm strainer, and resuspended in FACS buffer (2% BSA). Organoids were trypsinized to make single cell suspensions and passed through 70μm strainer before sorting. Cells were trypsinized to make single-cell suspension and passed through 70μm and/or 40μm filters before sorting. Sorted cells were collected in 50% FBS/DMEM containing 10μM Y27632.

For SOX9-pulldown experiments, lysates of Expi293 cells expressing ABF components were incubated with purified SOX9, immobilized on amylose resins for 4 hours at 4ºC. Unbound lysates were discarded and bound proteins were washed three times followed by western blot analysis.

Gene set enrichment analysis (GSEA v4.3.3)[6] was performed on pre-ranked gene lists ordered by the DESeq2 Wald statistic. Published gene signatures for intestinal stem cells (ISC), colon enterocytes, and stem cell programs were obtained from the Broad Institute MSigDB[6,7] and from Haber et al. (2017)[8], Munoz et al. (2012)[9], Kazakevych et al. (2017)[10], and Wang et al. (2020)[11]. A normalized enrichment score (NES) with FDR < 0.25 was considered significant, per GSEA recommended thresholds. Results were visualized as GSEA enrichment plots.

### Table S2 Number of chromatin loci co-bound by SOX9 and SMARCC1 in HEK-293T, HT-115, HT-29 and LS-180 cells related to Figure 1H

| cell_line | threshold | comparison | SOX9_replicate | SMARCC1_replicate | SOX9_column | SMARCC1_column | SOX9_boun | ARCC1_bou | SOX9_unique | SMARCC1_unique | Shared | ed_fraction_of_ | Avg Shared | Avg %shared |
|---|---|---|---|---|---|---|---|---|---|---|---|---|---|---|
| HEK | 10 | 9_r1_vs_SMARCC | 1 | 1 | HEK_Sx9_1 | HEK_SC1_1 | 4025 | 5502 | 526 | 2003 | 3499 | 86.90% | | |
| HEK | 10 | 9_r1_vs_SMARCC | 1 | 2 | HEK_Sx9_1 | HEK_SC1_2 | 4025 | 5450 | 602 | 2027 | 3423 | 85.00% | | |
| HEK | 10 | 9_r2_vs_SMARCC | 2 | 1 | HEK_Sx9_2 | HEK_SC1_1 | 4208 | 5502 | 589 | 1883 | 3619 | 86.00% | | |
| HEK | 10 | 9_r2_vs_SMARCC | 2 | 2 | HEK_Sx9_2 | HEK_SC1_2 | 4208 | 5450 | 684 | 1926 | 3524 | 83.70% | 3516.25 | 85.40% |
| HT115 | 10 | 9_r1_vs_SMARCC | 1 | 1 | HT115_Sx9_1 | HT115_SC1_1 | 9202 | 9331 | 733 | 862 | 8469 | 92.00% | | |
| HT115 | 10 | 9_r1_vs_SMARCC | 1 | 2 | HT115_Sx9_1 | HT115_SC1_2 | 9202 | 9498 | 570 | 866 | 8632 | 93.80% | | |
| HT115 | 10 | 9_r2_vs_SMARCC | 2 | 1 | HT115_Sx9_2 | HT115_SC1_1 | 9309 | 9331 | 789 | 811 | 8520 | 91.50% | | |
| HT115 | 10 | 9_r2_vs_SMARCC | 2 | 2 | HT115_Sx9_2 | HT115_SC1_2 | 9309 | 9498 | 616 | 805 | 8693 | 93.40% | 8578.5 | 92.68% |
| LS180 | 10 | 9_r1_vs_SMARCC | 1 | 1 | LS180_Sx9_1 | LS180_SC1_1 | 7274 | 9121 | 533 | 2380 | 6741 | 92.70% | | |
| LS180 | 10 | 9_r1_vs_SMARCC | 1 | 2 | LS180_Sx9_1 | LS180_SC1_2 | 7274 | 8825 | 610 | 2161 | 6664 | 91.60% | | |
| LS180 | 10 | 9_r2_vs_SMARCC | 2 | 1 | LS180_Sx9_2 | LS180_SC1_1 | 7633 | 9121 | 545 | 2033 | 7088 | 92.90% | | |
| LS180 | 10 | 9_r2_vs_SMARCC | 2 | 2 | LS180_Sx9_2 | LS180_SC1_2 | 7633 | 8825 | 620 | 1812 | 7013 | 91.90% | 6876.5 | 92.28% |
| HT29 | 10 | 9_r1_vs_SMARCC | 1 | 1 | DMSO_SOX9_1_Paul | DMSO_SMARCC1_1_Paul | 9615 | 13755 | 1058 | 5198 | 8557 | 89.00% | | |
| HT29 | 10 | 9_r2_vs_SMARCC | 2 | 1 | DMSO_SOX9_2_Paul | DMSO_SMARCC1_1_Paul | 9974 | 13755 | 1221 | 5002 | 8753 | 87.80% | | |
| HT29 | 10 | 9_r1_vs_SMARCC | 1 | 2 | DMSO_SOX9_1_Paul | DMSO_SMARCC1_2_Paul | 9615 | 15068 | 621 | 6074 | 8994 | 93.50% | | |
| HT29 | 10 | 9_r2_vs_SMARCC | 2 | 2 | DMSO_SOX9_2_Paul | DMSO_SMARCC1_2_Paul | 9974 | 15068 | 720 | 5814 | 9253 | 92.80% | 8889.25 | 90.78% |

### Table S2: List of Stem-like gene described in literature_related to Figure 2B-D and Figure S3D

| Stem-like signature genes | ISC Signature genes | Adult_ISC Signatue genes | Stem-like Signatue genes |
|---|---|---|---|
| Haber et.al., 2017 | Munoz et.al., 2012 | Kazakevych et.al., 2017 | Munoz et.al., 2012 |
| LGR5 | TLE3 | DPYSL5 | 1300002K09RIK |
| GKN3 | FLOT2 | PLCE1 | 6230424C14RIK |
| ASCL2 | CROCC | SYNE2 | ADRA2A |
| OLFM4 | SCML4 | NAV1 | AFAP1 |
| RGMB | FAM13A | LYST | ANGPT2 |
| IGFBP4 | PSIP1 | OSR2 | APCDD1 |
| 2210407C18RIK | CBX2 | ARHGEF28 | ARHGEF4 |
| JUN | SLC1A3 | MYCL1 | ARL4C |
| PDGFA | NME4 | TCF7 | ASCL2 |
| SOAT1 | NGEF | ANGPT2 | ATG16L2 |
| TNFRSF19 | L3MBTL3 | RNF32 | ATM |
| CYP2E1 | HES1 | MLL1 | AXIN2 |
| FSTL1 | ZMYM4 | PBX3 | B930095G15RIK |
| H2-EB1 | ENPP1 | SH3BP2 | CACHD1 |
| IFITM3 | GRLF1 | GM10548 | CAR12 |
| PRELP | DPYSL5 | UTRN | CASP12 |
| SCN2B | PHGDH | 1700001D01RIK | CBX6 |
| A930009A15RIK | STMN1 | ST3GAL3 | CC2D2A |
| H2-AB1 | ZFP280C | SAMD12 | CD320 |
| SLC1A2 | RBM38 | 1110028F11RIK | CDC42EP1 |
| CD74 | MARCKSL1 | AMICA1 | CDO1 |
| SP5 | ACSS1 | CPS1 | CES1D |
| NOXA1 | HK2 | CLCA4 | CGNL1 |
| RGCC | SLC25A4 | SLC6A7 | CITED4 |
| SORBS2 | HMGN1 | NGEF | CLCA1 |
| SECTM1B | PROM1 | SCML4 | CLCA2 |
| H2-AA | RRP1B | OAT | CLIC6 |
| CDO1 | CHEK1 | EHF | CPN1 |
| SLC14A1 | MGA | D630039A03RIK | CTTNBP2 |
| CLCA2 | FAM60A | ELF3 | CYP2E1 |
| TIFA | BCL11A | CLDN2 | D930015E06RIK |
| PLS3 | ITGA1 | PROSAPIP1 | DCT |
| HMGCS2 | ZMYM3 | SLC25A28 | DDB2 |
| ARID5B | THUMPD2 | STARD13 | DDO |
| AGR3 | PBLD | KRT23 | DYNC2H1 |
| SLC12A2 | FHIT | ARHGEF38 | E230029C05RIK |
| RASSF5 | CDCA5 | FHL2 | ELL3 |
| RNF43 | GTF2I | MYB | EMP2 |
| NRN1 | POLR1E | CCDC3 | EPHA4 |
| LAMB3 | ZFP422 | ACSL1 | EPHB3 |
| CD44 | SNTB2 | LGR5 | ESAM |
| AXIN2 | ESCO2 | SLC12A2 | ESRRG |
| SLC27A2 | ZFP629 | HOPX | EVL |
| AFAP1L1 | INSR | GAS6 | FARP1 |
| CCDC3 | BAG2 | ALG5 | FBXO21 |
| LRIG1 | PLSCR3 | LBH | FGFR4 |
| NOXO1 | UHRF2 | EGFR | FMNL2 |
| CDK6 | SLC27A1 | CARD10 | FSTL1 |
| AMICA1 | DKC1 | MYC | FZD2 |
| TGIF1 | DOCK11 | BLNK | GAS6 |
| TNS3 | TRIT1 | PMAIP1 | GKAP1 |
| NR2E3 | XPC | ARHGEF26 | GKN3 |
| EFNA4 | TIA1 | 1700040L02RIK | GPLD1 |
| RNF32 | MTUS1 | SEMA3B | GPR30 |
| PRSS23 | NARG2 | MAPKAPK3 | GRAMD1A |
| 2010009K17RIK | ZFP191 | IRF2BP2 | HUNK |
| SMOC2 | TRP53BP1 | AXIN2 | IGFBP4 |
| MECOM | CHEK2 | FOXA3 | IL17RD |
| ESRRG | CPS1 | GPR35 | IMMP2L |
| AQP1 | GINS4 | GMDS | IRS1 |
| ZNRF3 | MCM2 | FGFBP1 | KCNE3 |
| GRB7 | WHSC1 | ASCL2 | KLHL8 |
| PHGDH | MSH2 | ACSM3 | LARGE |
| 2410004N09RIK | TRIM28 | SOCS2 | LGR5 |
| CLCA4 | NRM | PRELID2 | LIPT2 |
| AQP4 | NFIB | QSOX2 | MAN2A2 |
| LCP1 | ATRX | MPZL2 | MEX3A |
| E030011O05RIK | EPHB2 | TMEM171 | MIA1 |
| SNHG1 | ZFP512 | TRIM65 | MPP3 |
| BC064078 | ZFP346 | GALNT12 | MSI1 |
| CAR12 | RPA2 | CAR12 | MYC |
| ZBTB38 | SURF2 | LHPP | MYCL1 |
| CDCA7 | HELLS | METTL23 | NAP1L1 |
| FAM13A | CCNB1 | ESRRG | NAPEPLD |
| SHISA2 | TTK | YDJC | NOTCH1 |
| DTX4 | ZFP451 | PTPLA | NOTCH2 |
| SLC19A2 | DNAJC9 | CCDC28A | PHLDB2 |
| FAM115C | POLA2 | CLUH | PLA2G4A |
| MIR703 | PDXK | PLAUR | PLEKHB1 |
| CD14 | SIPA1L1 | LMCD1 | PLXNB1 |
| METTL20 | ACSS2 | FOSL2 | POGK |
| MYO9A | PDP1 | SH2D4A | PPAP2B |
| APP | APEX1 | TMSB4X | PRELP |
| CLIC6 | SPATA13 | RPIA | PRKD3 |
| WEE1 | ZFP280D | 1300002K09RIK | PSD3 |
| 2410006H16RIK | HMGB3 | FAM195A | PSRC1 |
| LANCL1 | MCM5 | GIMAP5 | PTPRO |
| 1500012F01RIK | ZFP828 | KCNE3 | PTPRS |
| CASP12 | UNC84B | DDIT4 | RAD51L1 |
| SH3RF1 | MCM6 | HS3ST1 | RASA3 |
| LRP4 | CLIC4 | PLA2G4A | RGMB |
| ARHGEF26 | VGLL4 | SMOC2 | RGS12 |
| ETV6 | CDK4 | SNHG12 | RNF32 |
| 1700024F13RIK | MACROD1 | RPS12 | RNF43 |
| CTTNBP2 | TMX4 | PIGYL | SCN2B |
| SLC16A13 | NEIL1 | RANGRF | SECTM1B |
| HTR4 | MCM3 | FAM46C | SEMA7A |
| PDXK | ZMYM2 | MRPL12 | SESTD1 |
| IMMP2L | NMRAL1 | IER3 | SFRP5 |
| RPS15A-PS6 | PGCP | 3230401D17RIK | SHISA2 |
| RPS15A-PS4 | NUSAP1 | MRPS12 | SLC19A2 |
| NAP1L1 | MYCBP2 | GNG11 | SLC23A3 |
| SDC4 | MCM7 | HMGCS2 | SLC27A2 |
| EPN3 | SMAD5 | SNHG8 | SLCO3A1 |
| SIPA1L1 | MECP2 | 2810008D09RIK | SMO |
| WFDC15B | BCL11B | 2410006H16RIK | SNX10 |
| ZFP341 | INCENP | PSMG3 | SORCS2 |
| NGEF | HMGA1 | 2310039H08RIK | SOX4 |
| NRG4 | NUP210 | KCNQ1 | SOX9 |
| CSAD | MCM4 | CLCA2 | SP5 |
| RPL34-PS1 | FOXP4 | MCC | SRL |
| RIN2 | RAD50 | MUC4 | SYCN |
| CD81 | LZTS2 | SLCO3A1 | TEAD2 |
| IRF2BP2 | POU2F1 | NFIX | TEX9 |
| SESN3 | FLNA | RGMB | TFPI |
| PHLPP1 | RTN4 | DNAHC8 | TGIF2 |
| YAP1 | FHL2 | IFITM3 | TMEM209 |
| MFGE8 | HRSP12 | ACPP | TNFRSF19 |
| ZFP825 | SDSL | DAPK2 | TRIM44 |
| ITGA1 | NUBPL | LIMK2 | TTC21B |
| PCDH8 | MBP | NFIA | TTPA |
| VDR | IVD | CLCA1 | TXNDC16 |
| KCNQ1 | ADNP | GM8979 | UTRN |
| SLC28A2 | ERI1 | DACH1 | VAV3 |
| ZFP36L1 | PKIG | RASA3 | WDR35 |
| UROD | BPHL | CTTNBP2 | WDR90 |
| RGS12 | SPIN1 | SEMA7A | WWP1 |
| NFIB | LHPP | PROM1 | WWTR1 |
| SDSL | UROD | ITGA1 | ZFP12 |
| NFIA | ISG20L2 | NOXA1 | ZKSCAN17 |
| | DTD1 | APOBEC3 | ZNRF3 |
| | LYPLAL1 | SLC1A2 | ZRANB3 |
| | ATXN10 | AQP4 | ACSM3 |
| | METT5D1 | CD44 | AFAP1L1 |
| | PHF6 | CARD11 | AGR3 |
| | RNF2 | ZBTB38 | AQP4 |
| | ARSB | OLFM4 | CD44 |
| | PPP1R1B | FAM49A | CDCA7 |
| | ZFP740 | KCNN4 | CDK6 |
| | FTO | 4933417O13RIK | CLCA4 |
| | WDR55 | SLC14A1 | CNN3 |
| | POLR1A | SEMA5A | DACH1 |
| | CHCHD6 | HOOK1 | DMPK |
| | NHP2 | EPHB3 | FAM115C |
| | BRD8 | LRIG1 | FZD7 |
| | BCKDHA | FAM65B | GLRX |
| | CYP2E1 | CDO1 | GRB7 |
| | LGR5 | CD177 | KCNQ1 |
| | TNFRSF19 | SORCS2 | LCP1 |
| | FSTL1 | E030011O05RIK | MECOM |
| | GKN3 | SHANK2 | NAV1 |
| | SCN2B | EPN3 | OLFM4 |
| | CES1D | CNNM1 | PAICS |
| | PRELP | GPR110 | PICALM |
| | SP5 | CDCA7 | |
| | E230029C05RIK | BC021891 | SLC12A2 |
| | IL17RD | TNFAIP8L1 | SMOC2 |
| | FMNL2 | CERS4 | SOAT1 |
| | ARL4C | TNS3 | SORBS2 |
| | CTTNBP2 | GM684 | TNS3 |
| | RGMB | GPR30 | VDR |
| | ANGPT2 | PRELP | 1500011H22RIK |
| | AXIN2 | SCN2B | 2310008H04RIK |
| | HUNK | SLC1A3 | 4933439C10RIK |
| | CAR12 | AI464131 | ACPL2 |
| | PTPRO | RASSF4 | ACVR2B |
| | DCT | FHDC1 | ADRBK2 |
| | SFRP5 | NCALD | AI480653 |
| | SYCN | CHRM3 | ALDH7A1 |
| | CLCA2 | FHOD1 | ALMS1 |
| | 1300002K09RIK | DGKG | ALS2CR4 |
| | CASP12 | GKN3 | APP |
| | EMP2 | AGR3 | AQP1 |
| | MIA1 | ZDHHC14 | ARHGAP39 |
| | PSRC1 | STK39 | ASPM |
| | NOTCH1 | NR2E3 | ATP11A |
| | RNF43 | CAP2 | ATR |
| | DDO | IGSF9B | AU020206 |
| | ARHGEF4 | SLC27A1 | BCKDHB |
| | CLCA1 | MLXIP | BCL2 |
| | EVL | WHRN | BCL7A |
| | UTRN | CBS | BLNK |
| | ESRRG | HTR4 | BRCA2 |
| | ESAM | MGAT4B | C79407 |
| | FGFR4 | ST6GALNAC2 | CCBL2 |
| | SEMA7A | PSD3 | CCDC18 |
| | WWTR1 | CASP12 | CDCA7L |
| | SORCS2 | LDLRAD4 | CENPF |
| | RAD51L1 | 1700003D09RIK | CEP192 |
| | IRS1 | RASSF5 | CIT |
| | MPP3 | AFAP1L1 | CKAP2 |
| | TTC21B | SLIT2 | CSAD |
| | CDO1 | ITGA6 | CSNK1E |
| | PLA2G4A | PDCD4 | DAPK2 |
| | ATG16L2 | SIPA1L1 | DCTD |
| | EPHA4 | GAS7 | DTL |
| | SMO | ZBTB20 | EFNA4 |
| | CPN1 | SHISA2 | EHF |
| | KLHL8 | ACPL2 | ELMO1 |
| | RNF32 | NAPEPLD | ENGASE |
| | FZD2 | ATP8A2 | ERI2 |
| | D930015E06RIK | ADRA2A | ESPL1 |
| | TGIF2 | TRAF3IP2 | ETV6 |
| | WDR35 | PHF15 | FADS1 |
| | ELL3 | CITED4 | FAM115A |
| | APCDD1 | FRY | FAM189B |
| | CDC42EP1 | E230029C05RIK | FAM92A |
| | FZD7 | PLEKHG5 | FHDC1 |
| | AFAP1 | RDH1 | FRAS1 |
| | MEX3A | DENND4C | GABBR1 |
| | ADRA2A | BOC | GEMIN8 |
| | SHISA2 | SZT2 | GINS1 |
| | POGK | FILIP1L | GM3626 |
| | GRAMD1A | ENGASE | GM5918 |
| | SOX4 | RNF39 | HAUS4 |
| | NAPEPLD | CREG1 | HDAC11 |
| | TXNDC16 | 2210039B01RIK | HEMGN |
| | SECTM1B | IGF1R | HIRIP3 |
| | SLC23A3 | 4933406C10RIK | IFITM2 |
| | CITED4 | KANK1 | IFT122 |
| | B930095G15RIK | PIK3R1 | ILF3 |
| | EPHB3 | ATXN7L1 | ISYNA1 |
| | GPR30 | SDC4 | KIF12 |
| | PPAP2B | SMARCA2 | KIF20A |
| | ZNRF3 | A930009A15RIK | KLHL23 |
| | TFPI | LONRF3 | KRT23 |
| | PLEKHB1 | UTY | LANCL1 |
| | RGS12 | VPS13A | LASS6 |
| | FARP1 | ITIH5 | LIMK2 |
| | CD320 | MYO5C | LPAR6 |
| | CC2D2A | 4930474M22RIK | LRIG1 |
| | CBX6 | GM10791 | LYSMD2 |
| | TEAD2 | SLC9A5 | MAGED1 |
| | MYCL1 | JUN | MARVELD1 |
| | RASA3 | ASCC2 | MCC |
| | PLXNB1 | UCK2 | MCM8 |
| | MAN2A2 | AIG1 | MDN1 |
| | LARGE | CCDC101 | MFGE8 |
| | NAP1L1 | PDE4B | MIF |
| | TRIM44 | RHOB | MLH3 |
| | PRKD3 | TPT1 | MLXIP |

GKAP1
SRL
TEX9
6230424C14RIK
GPLD1
CACHD1
MYC
ZRANB3
WWP1
LIPT2
DYNC2H1
GAS6
ATM
PTPRS
SNX10
ZFP12
KCNE3
TTPA
WDR90
IMMP2L
DMPK
FBXO21
PSD3
DDB2
ZKSCAN17
SLC19A2
CLIC6
AFAP1L1
ASCL2
SORBS2
NAV1
SLCO3A1
OLFM4
AGR3
MSI1
SOAT1
CDCA7
CD44
CDK6
AQP4
CGNL1
SLC12A2
MECOM
TNS3
PHLDB2
SMOC2
DACH1
CLCA4
LCP1
FAM115C
ACSM3
KCNQ1
SOX9
PAICS

CNN3
PICALM
GLRX
VDR
GRB7
SESN3
ZBTB38
FAM115A
PTPRM
RASSF4
ZFHX3
PDE3B
FHDC1
ZFP292
TUBB2B
EFNA4
LRIG1
FRAS1
ADRBK2
KIF12
PRKACB
ATP11A
SPATA24
POLI
MCC
SLC44A2
TBC1D19
ZFP462
1500011H22RIK
ARHGAP39
ZBTB12
EHF
APP
GM5918
TRIM37
MARVELD1
PPP1R9A
PELI2
TNS4
MFGE8
SLC39A10
PPAT
ZFP566
KLHL23
TIAM1
GM3626
AQP1
TMEM132A
RHBDF1
2310008H04RIK
PBX1
TNFSF10
ALS2CR4
PHLPP1
IFT122
BCL2
LYSMD2
ZFP397
TSPAN5
ACVR2B
MLH3
BRCA2
ZFP275
ENGASE
NELF
GABBR1
RGNEF
PPM1F
PRRC2B
AU020206
FADS1
NOD1
RNF219
4933439C10RIK
PNPLA7
ZFP704
C79407
ERI2
TNFAIP8L1
TTC23
MPZL3
ZBTB25
MAGED1
FAM92A
CCDC18
ZFP783
KRT23
MLXIP
LANCL1
LPAR6
ZFP553
POGLUT1
POLE
MCM8
HAUS4
ZFP90
PLCE1
FAM189B
PDS5B
TGIF1
PHC1
BLNK
SPICE1
ASPM
ISYNA1
PALB2
HIRIP3
ALMS1
LASS6
MPZL1
CENPF
MDN1
SEMA3B
ZMYM1
CEP192
UNG
PRPF40B
ATR
AI480653
ZFP318
TMEM9
GINS1
IFITM2
VNN1
SMPX
TMEM194
DTL
CSNK1E
DCTD
XRCC5
NT5C3L
ALDH7A1
LIMK2
ETV6
ELMO1
DAPK2
NFIC
TIMELESS
CKAP2
MYO1B
NFIA
CSAD
TBC1D4
BCKDHB
BCL7A
PCK2
YAP1
STK39
RPRD1A
ILF3
KIF20A
MIF
MKL2
FAM120C
NCOR2
PCDH8
RASSF5
SLC14A1
NR2E3
1190002H23RIK
SMARCD3
AK7
BHLHB9
PRSS23
IFITM3
WFDC15B
RDH16
ZFP618
LECT2
H2-EB1
IFFO2
ZC3HAV1L
KLHL24
TBC1D9
CIITA
NRTN
NRG4
TCF7
GREB1
NRN1
AI747448
DTX4
TRIM32
RASL11B
ZFP503
BC021891
PDGFA
TLR2
S100PBP
4933431E20RIK
FGD1
1110051M20RIK
SLC16A13

ARID5B
STEAP3
CNP
KLHL8
ATOH8
MKL2
MMD
MIR1192
CYP2E1
TMEM63A
PPP2R3A
RNF43
JUND
NFIB
ETS2
SLC9A8
TNFRSF19
SIDT1
SMARCD2
GLCCI1
EDN1
ADCY6
BCL9L
MB21D1
SNX10
DIAP2
SLC7A2
PLEKHA6
PCCA
D2HGDH
TMEM194
NFKBID
2010009K17RIK
CDKN1B
TRIB1
NOD1
IL17RD
FAM115C
TGM2
TOX
PTPRM
HDAC4
SYPL
SORT1
MPZL3
NRBP2
USP43
RGS3
ZFP612
LGALS12
CYP4F15
ETV6
RAPGEF5
B3GALTL
MAP3K14
LENG8
NCEH1
8430419L09RIK
WIPI1
CYR61
A730020E08RIK
FRYL
MANSC1
2010111I01RIK
MAP3K5
SIMC1
PCK2
OPHN1
CDK6
PNPLA7
ARHGAP39
FAM120C
METTL20
KRT18
CCBE1
GLTSCR1
GLTSCR1L
CCDC171
P2RY6
TIFA
GDF15
GM6548
DMPK
MBP
GM15713
CHD9
XYLB
EPHB2
MYO9A
CFTR
NR4A2
RGCC
PRICKLE3
DOPEY1
RASEF
TFPI
GPX2-PS1
EFNA3
DDB2
ICA1
ALPK1
KLHL24
NOTCH1
AI414108
FRMD4B
TTLL10
4933439C10RIK
SFI1
ATF7
SH3RF1
SORBS2
CYP39A1
PAWR
LRRK1
DNAJB1
DYM
CCPG1
AMIGO1
CDC42EP1
MECOM
PIK3IP1
CIT
SLFN9
HDAC8
PBX1
B230120H23RIK
CLYBL
RBM43
SORL1
VEZF1
SRD5A1
GPS2
GM19461
ABHD14A
PDE8B
POLR3GL
MCTP2
MLL2
ITGA2
IGBP1
ACHE
E030018B13RIK
GTF2IRD2
WDR16
ELP5
NHS
2310040G24RIK
CD81
ABHD11
ZFP341
NFATC2IP
GM7120
GLTSCR2
ZDHHC1
4930414L22RIK
6430573F11RIK
ADRBK2
CNTD1
SETD7
LIPT2
SPATS1
CPQ
ZAN
DCT
PVR
MYCBPAP
CNPPD1
SNORA31
EHBP1
LFNG
IFFO2
LRCH1
SLC1A5
MIR568
NEIL3
TPD52L1
CBX7
GRK5
RING1
GM53
IFITM1
FYCO1
TNFRSF12A
GM16907
DAGLA
PM20D2
DCP1B
PPT2
ATXN7L2
SLC7A4
EFNA4
MCM9
ARHGEF39
A430071A18RIK
VAMP1
PLCH2
CCND1
CEP70
WWOX
GNE
RPL38
AK7
ZC3H6
ATP2C2
G6PC3
DYNLL2
SULT1A1
HSD17B14
RIPK2
1110032A03RIK
NHLRC1
SGSM3
4930415O20RIK
CLN3
PTMS
HOXB6
GM2A
ISG20
SMTN
HIST1H4K
0610009O20RIK
NSMF
ATG16L2
HSPB6
RPS13
PGC
TBC1D4
GLIPR2
LTC4S
NUDT6
METTL15
HIST1H3F
EMP2
FBXO24
TCAIM
1700031P21RIK
EGR3
LYRM9
9330162B11RIK
LIN37
DDRGK1
SYCN
4930471I20RIK
RP9
SIK1
PLEKHB1
CCNL1
HIST1H2BJ

MPZL1
MPZL3
MYO1B
NEDD4
NELF
NFIA
NFIC
NIN
NOD1
NT5C3L
ORC1
PALB2
PBX1
PCK2
PDE3B
PDS5B
PELI2
PHC1
PHLPP1
PLCE1
PLS3
PNPLA7
POGLUT1
POLD1
POLE
POLI
PPAT
PPM1F
PPP1R9A
PRKACB
PRPF40B
PRPS2
PRRC2B
PTPRM
QSOX2
RASSF4
RGNEF
RHBDF1
RNF219
RPRD1A
SEMA3B
SESN3
SLC39A10
SLC44A2
SMPX
SORD
SPAG5
SPATA24
SPICE1
STK39
TBC1D19
TBC1D4
TGIF1
TIAM1
TIMELESS
TMEM132A
TMEM194
TMEM9
TNFAIP8L1
TNFSF10
TNS4
TRIM37
TSPAN5
TTC23
TUBB2B
UNG
VNN1
XRCC5
YAP1
ZBTB12
ZBTB25
ZBTB38
ZFHX3
ZFP275
ZFP292
ZFP318
ZFP397
ZFP462
ZFP553
ZFP566
ZFP704
ZFP783
ZFP90
ZMYM1
1110051M20RIK
1190002H23RIK
1700040L02RIK
4933431E20RIK
ACOT1
AI747448
AK7
BC021891
BHLHB9
CCDC111
CIITA
CLU
DTX4
FAM171A1
FGD1
GREB1
H2-AA
H2-EB1
HMBOX1
HMGA2
IFFO2
IFITM3
IMPDH2
IRF2BP2
KANK1
KLHL24
LAMB3
LECT2
NR2E3
NRG4
NRN1
NRTN
PCDH8
PDE8B
PDGFA
PHF20
PRSS23
RASL11B
RASSF5
RCCD1
RDH16
RHOBTB3
S100PBP
SDC4
SGK1
SLC14A1
SLC16A13
SMARCD3
TBC1D9
TCF7
TIFA
TLR2
TRIM24
TRIM32
WFDC15B
ZC3HAV1L
ZFP503
ZFP618
ZFP956

| | |
|---|---|
| KANK1 | ATP6V1G2 |
| CLU | PIK3AP1 |
| PHF20 | NR4A1 |
| ZFP956 | GPR19 |
| 1700040L02RIK | GM15545 |
| RHOBTB3 | HIST1H3C |
| FAM171A1 | 4933426D04RIK |
| PDE8B | HSBP1L1 |
| RCCD1 | BC025920 |
| SGK1 | ARC |
| IMPDH2 | DUSP10 |
| IRF2BP2 | UBE2S |
| ACOT1 | ZFP579 |
| TIFA | EGR1 |
| TRIM24 | SNORD47 |
| HMGA2 | GM5918 |
| HMBOX1 | GM14318 |
| HMGCS2 | TSACC |
| ARID5B | ZFP661 |
| NOXA1 | HIST1H3A |
| | FGL1 |
| | RPL13 |
| | CBX8 |
| | RPS25 |
| | ZFP956 |
| | PER1 |
| | KLF11 |
| | SCARNA17 |
| | 3110079O15RIK |
| | THBS1 |
| | GM5129 |
| | HIST1H3D |
| | MIR429 |
| | IMP3 |
| | TMED6 |
| | FKBP7 |
| | PIM3 |
| | MIR694 |
| | MIR1190 |
| | MIR1941 |
| | D630032N06RIK |
| | SNORD12 |
| | SCARNA9 |
| | MIR5106 |
| | GM16675 |
| | MIR210 |
| | SNORA47 |
| | DTX4 |
| | RBM38 |
| | FARP1 |
| | IVNS1ABP |
| | EPHB4 |
| | SPAG5 |
| | TAF4B |
| | SLC16A11 |
| | NLE1 |
| | PLK1S1 |
| | FAM214A |
| | HES1 |
| | ZBTB44 |
| | TROAP |
| | GM13212 |
| | 2610015P09RIK |
| | SFMBT1 |
| | RAD52 |
| | LRDD |
| | OVGP1 |
| | ZFP398 |
| | KNSTRN |
| | RPS15 |
| | CCHCR1 |
| | RCCD1 |
| | EIF2A |
| | FADS2 |
| | RPL10A |
| | RPS7 |
| | MFGE8 |
| | THAP3 |
| | ARHGAP8 |
| | WWP1 |
| | BAIAP2 |
| | B130034C11RIK |
| | CCDC61 |
| | 2610035D17RIK |
| | AGXT2L2 |
| | PBK |
| | 1810022K09RIK |
| | HIST1H1D |
| | GGNBP1 |
| | PTPRD |
| | METTL22 |
| | PDGFA |
| | FERMT1 |
| | EEF1B2 |
| | RWDD3 |
| | KCNK1 |
| | 1810014B01RIK |
| | HYAL3 |
| | CCDC163 |
| | GNAT2 |
| | CCNB2 |
| | METAP1D |
| | 1500012F01RIK |
| | DANCR |
| | CYB5RL |
| | LRRC51 |
| | PPIFOS |
| | RPS5 |
| | RSL1 |
| | ALG1 |
| | RPS26 |
| | RPL32 |
| | RPS9 |
| | TMEM107 |
| | GNB2L1 |
| | 1190007I07RIK |
| | RPS20 |
| | HSPH1 |
| | RPS10 |
| | RPS15A-PS4 |
| | PSMG4 |
| | HIST1H4I |
| | 1700007L15RIK |
| | 1190002F15RIK |
| | RPL22L1 |
| | SNORA62 |
| | MIR1960 |
| | SNORD65 |

**Table S3: List of genes related to Figure S2E**

| target_chr | target_start | target_end | bait_chr | bait_start | bait_end | X | CHiCAGO.Score | Number.of.reads | target_name | bait_name | target_peak_name | target2enhancer | Promoter_bait | gene | distanceToTSS | Bait_gene | Bait_distanceToTSS |
|---|---|---|---|---|---|---|---|---|---|---|---|---|---|---|---|---|---|
| chr1 | 203477867 | 203481430 | chr1 | 203270113 | 203274326 | interaction_10754 | 10.79 | 57 | 46996-203450558:targ | 9242-203243454: | r1:203477867-2034814 | chr1:203480781-203481181 | 1:203270113-2032743 | PRELP | 4975 | LINC01353 | 0 |
| chr1 | 203477867 | 203481430 | chr1 | 203350754 | 203354000 | interaction_10763 | 8.55 | 93 | 46996-203450558:targ | 9883-203323128: | r1:203477867-2034814 | chr1:203480781-203481181 | 1:203350754-2033540 | PRELP | 4975 | FMOD | 0 |
| chr5 | 52921450 | 52923400 | chr5 | 52787557 | 52789141 | interaction_35275 | 5.45 | 104 | 17281-52219230:targ | 3392-52084975:B | chr5:52921450-5292340 | chr5:52922363-52922763 | r5:52787557-5278914 | ITGA1 | -8389 | LOC105378963 | 0 |
| chr5 | 149320108 | 149322122 | chr5 | 149551756 | 149555421 | interaction_39423 | 5.07 | 27 | 99672-148701685:targ | 1320-148934984: | r5:149320108-1493221 | chr5:149320803-149321203 | 5:149551756-1495554 | AFAP1L1 | 5163 | ARHGEF37 | 0 |
| chr7 | 92698015 | 92699204 | chr7 | 92831298 | 92838147 | interaction_52127 | 8.24 | 69 | 27330-92328518:targ | 0613-92467461:B | chr7:92698015-92699204 | chr7:92698463-92698863 | r7:92831298-9283814 | CDK6 | 26847 | CDK6-AS1 | 0 |
| chr7 | 117716898 | 117722215 | chr7 | 117460295 | 117472070 | interaction_53372 | 5.78 | 52 | 56953-117362269:targ | 0350-117112124: | r7:117716898-11772221 | chr7:117716921-117717321 | 7:117460295-1174720 | CTTNBP2 | 10887 | LOC105375468 | 0 |
| chr11 | 102268499 | 102269206 | chr11 | 101870606 | 101876440 | interaction_73861 | 5.16 | 45 | 139231-102139937:ta | 41338-101747171 | r11:102268499-1022692 | chr11:102268638-10226903 | 11:101870606-101876 | YAP1 | 41665 | TRPC6 | 0 |
| chr11 | 102268499 | 102269206 | chr11 | 101912108 | 101921139 | interaction_73863 | 5.07 | 28 | 139231-102139937:ta | 82840-101791870 | r11:102268499-1022692 | chr11:102268638-10226903 | 11:101912108-101921 | YAP1 | 41665 | ANGPTL5 | 0 |
| chr11 | 102268499 | 102269206 | chr11 | 102109745 | 102114597 | interaction_73871 | 7.09 | 77 | 139231-102139937:ta | 80477-101985328 | r11:102268499-1022692 | chr11:102268638-10226903 | 11:102109745-102114 | YAP1 | 41665 | YAP1 | 0 |
| chr15 | 71744683 | 71750775 | chr15 | 71114395 | 71116621 | interaction_90533 | 6.05 | 35 | 37023-72043114:targ | 06735-71408960:E | hr15:71744683-7175077 | chr15:71746588-71746988 | r15:71114395-711166 | NR2E3 | -45650 | CT62 | 0 |

### Table S4: GSEA related to Figure S3D

| | NES | pvalue | p.adjust | core_enrichment | group | Number of genes |
|---|---|---|---|---|---|---|
| Colon signature (Wang 2020) | 2.682666866 | 1.00E-10 | 1.90E-09 | .DN4/FTL/APOC3/ADI | D4.vs.DMSO | 79 |
| Genes enriched in enterocytes cluster | 2.375025955 | 1.00E-10 | 1.90E-09 | KRT8/RNF128/IRF7/G | D4.vs.DMSO | 166 |
| Enterocyte colon signature (Wang 2020) | 1.937653778 | 7.25E-08 | 9.18E-07 | S100A11/TMED9/MUC | D4.vs.DMSO | 92 |
| Stem_Haber_2017 | -2.079043846 | 1.27E-06 | 1.45E-05 | 8/CCDC3/NRN1/LRIG | D4.vs.DMSO | 30 |
| adult.ISC.signature_Kazakevych_2017 | -1.669532303 | 3.31E-06 | 3.43E-05 | K2/MCC/KCNN4/KAN | D4.vs.DMSO | 72 |
| Stem_Munoz_2012 | -1.622557331 | 7.01E-05 | 0.000570526 | /SLC12A2/PCK2/MCC | D4.vs.DMSO | 53 |
| ISC-marker genes (Munoz et al., 2012) | -1.560933831 | 8.81E-05 | 0.000669241 | A2/PCK2/MCC/KANK1 | D4.vs.DMSO | 60 |
| adult.epithelium.signature_Kazakevych_2017 | 1.634074392 | 0.000109731 | 0.000735844 | CDC107/REC8/NTN4/ | D4.vs.DMSO | 114 |

### Table S5: P-values related to Figure 3E-F and S3G

| group1 | group2 | n1 | n2 | median_group1 | median_group2 | delta_median | U | z | log10_p | neg_log10_p | p_value_display | group |
|---|---|---|---|---|---|---|---|---|---|---|---|---|
| DMSO | D2 | 12811 | 12811 | 1.605134363 | 1.052714543 | -0.55241982 | 153245839 | 120.2650847 | -3142.928735 | 3142.928735 | P < 2.23e-308 | SOX9 |
| DMSO | D4 | 12811 | 12811 | 1.605134363 | 0.739106477 | -0.866027886 | 162752246 | 136.4057458 | -4042.589022 | 4042.589022 | P < 2.23e-308 | SOX9 |
| D4 | D7 | 12811 | 12811 | 0.739106477 | 1.19386254 | 0.454756064 | 19139131 | -106.3683603 | -2458.978322 | 2458.978322 | P < 2.23e-308 | SOX9 |
| D4 | D10 | 12811 | 12811 | 0.739106477 | 1.523559169 | 0.784452692 | 2068384 | -135.2245415 | -3972.913368 | 3972.913368 | P < 2.23e-308 | SOX9 |
| D4 | D13 | 12811 | 12811 | 0.739106477 | 1.617638116 | 0.878531639 | 2376630 | -134.704476 | -3942.428398 | 3942.428398 | P < 2.23e-308 | SOX9 |
| DMSO | D2 | 11518 | 11518 | 1.577164833 | 1.466044777 | -0.111120056 | 81725153 | 30.50185263 | -203.6086132 | 203.6086132 | P = 2.463e-204 | SMARCC1 |
| DMSO | D4 | 11518 | 11518 | 1.577164833 | 1.349290608 | -0.227874225 | 95200229 | 57.20347985 | -712.4130917 | 712.4130917 | P < 2.23e-308 | SMARCC1 |
| D4 | D7 | 11518 | 11518 | 1.349290608 | 1.291549144 | -0.057741464 | 71629445 | 10.49693393 | -25.04948202 | 25.04948202 | P = 8.923e-26 | SMARCC1 |
| D4 | D10 | 11518 | 11518 | 1.349290608 | 1.535478036 | 0.186187428 | 42791872 | -46.64628043 | -474.252433 | 474.252433 | P < 2.23e-308 | SMARCC1 |
| D4 | D13 | 11518 | 11518 | 1.349290608 | 1.558715745 | 0.209425137 | 41856937 | -48.49938003 | -512.5555195 | 512.5555195 | P < 2.23e-308 | SMARCC1 |
| DMSO | D2 | 7775 | 7775 | 1.870353612 | 1.742075021 | -0.128278591 | 37496950 | 25.98026058 | -148.0820732 | 148.0820732 | P = 8.278e-149 | ATAC |
| DMSO | D4 | 7775 | 7775 | 1.870353612 | 1.707841948 | -0.162511664 | 39258775 | 32.27496731 | -227.8038714 | 227.8038714 | P = 1.571e-228 | ATAC |
| D4 | D7 | 7775 | 7775 | 1.707841948 | 1.833187532 | 0.125345584 | 23425556.5 | -24.29432879 | -129.6477285 | 129.6477285 | P = 2.250e-130 | ATAC |
| D4 | D10 | 7775 | 7775 | 1.707841948 | 1.847091941 | 0.139249993 | 22711626 | -26.84507787 | -158.0164662 | 158.0164662 | P = 9.628e-159 | ATAC |
| D4 | D13 | 7775 | 7775 | 1.707841948 | 1.837569139 | 0.129727192 | 23333130.5 | -24.62455124 | -133.1613935 | 133.1613935 | P = 6.896e-134 | ATAC |
| DMSO | D2 | 7511 | 7511 | 1.756895724 | 1.718787992 | -0.038107732 | 31203210 | 11.27211941 | -28.744294 | 28.744294 | P = 1.802e-29 | H3K4me1 |
| DMSO | D4 | 7511 | 7511 | 1.756895724 | 1.63166679 | -0.125228934 | 36480940 | 31.13132356 | -212.041974 | 212.041974 | P = 9.079e-213 | H3K4me1 |
| D4 | D7 | 7511 | 7511 | 1.63166679 | 1.749532482 | 0.117865692 | 21053423 | -26.91980539 | -158.8901062 | 158.8901062 | P = 1.288e-159 | H3K4me1 |
| D4 | D10 | 7511 | 7511 | 1.63166679 | 1.752544841 | 0.120878051 | 20540853 | -28.8485187 | -182.2766789 | 182.2766789 | P = 5.288e-183 | H3K4me1 |
| D4 | D13 | 7511 | 7511 | 1.63166679 | 1.744809818 | 0.113143028 | 21021794.5 | -27.03881738 | -160.2864762 | 160.2864762 | P = 5.170e-161 | H3K4me1 |
| DMSO | D2 | 4228 | 4228 | 1.242333093 | 0.995078935 | -0.247254158 | 12508795 | 31.82667273 | -221.5577747 | 221.5577747 | P = 2.768e-222 | H3K4me3 |
| DMSO | D4 | 4228 | 4228 | 1.242333093 | 0.902151111 | -0.340181982 | 14138718 | 46.35374367 | -468.3420117 | 468.3420117 | P < 2.23e-308 | H3K4me3 |
| D4 | D7 | 4228 | 4228 | 0.902151111 | 1.046128634 | 0.143977523 | 7188482.5 | -15.6270853 | -54.32231309 | 54.32231309 | P = 4.761e-55 | H3K4me3 |
| D4 | D10 | 4228 | 4228 | 0.902151111 | 1.052261626 | 0.150110515 | 6826407.5 | -18.83322022 | -78.39419522 | 78.39419522 | P = 4.035e-79 | H3K4me3 |
| D4 | D13 | 4228 | 4228 | 0.902151111 | 1.077269669 | 0.175118558 | 6682526.5 | -20.13188048 | -89.41118168 | 89.41118168 | P = 3.880e-90 | H3K4me3 |
| DMSO | D2 | 5721 | 5721 | 1.476284473 | 1.453910099 | -0.022374374 | 16889601 | 2.969925452 | -2.525970283 | 2.525970283 | P = 2.979e-03 | H3K27ac |
| DMSO | D4 | 5721 | 5721 | 1.476284473 | 1.260325974 | -0.215958499 | 21998584 | 31.88946042 | -222.4273458 | 222.4273458 | P = 3.738e-223 | H3K27ac |
| D4 | D7 | 5721 | 5721 | 1.260325974 | 1.462560561 | 0.202234587 | 11473301 | -27.6892048 | -168.026001 | 168.026001 | P = 9.419e-169 | H3K27ac |
| D4 | D10 | 5721 | 5721 | 1.260325974 | 1.389407275 | 0.129081301 | 13608504 | -15.60304292 | -54.1586055 | 54.1586055 | P = 6.941e-55 | H3K27ac |
| D4 | D13 | 5721 | 5721 | 1.260325974 | 1.462848874 | 0.202522899 | 11869588 | -25.44641629 | -142.1115584 | 142.1115584 | P = 7.735e-143 | H3K27ac |

### Table S6. Peak stats related to Figure 3E-F, 3H-I and S3G

| | Count in DMSO | | D4 vs. DMSO | | | D4 vs. DMSO | | | | |
|---|---|---|---|---|---|---|---|---|---|---|
| Antibody | Total | Promoter | s_in_Prom | Gain_in_Promoter | Enhancer | s_in_Enhar | n_in_Enhar | Total_loss | | |
| SMARCC1 | 12697 | 2628 | 2439 | 189 | 10069 | 9037 | 1032 | 11476 | | |
| ARID1A | 11718 | 2568 | 1923 | 645 | 9150 | 5970 | 3180 | 7893 | 65.2% | Loss |
| SMARCB1 | 11425 | 2529 | 1399 | 1130 | 8896 | 5477 | 3419 | 6876 | 61.6% | Loss |
| BRD9 | 10517 | 2494 | 710 | 1784 | 8023 | 4123 | 3900 | 4833 | 71.5% | Gain |
| PBRM1 | 10230 | 2466 | 631 | 1835 | 7764 | 3597 | 4167 | 4228 | 74.4% | Gain |
| ATAC | 12412 | 2573 | 1159 | 1414 | 9839 | 6608 | 3231 | 7767 | | |
| H3K4me1 | 12574 | 2608 | 1667 | 941 | 9966 | 5825 | 4141 | 7492 | | |

Note: Threshold=1

Note: The Total refers to the number of peaks in the SOX9 loss universe (12,824 peaks), specifically the peaks present under DMSO for each antibody. These peaks were then categorized into promoters and enhancers, followed by counting the loss and gain during the dTAG time course (Day 4 vs. DMSO).

http://localhost:17449/lab/tree/projects/BAF_SOX9/diffbind/2.2-Diffbind-Raghwan.ipynb#enhancer-and-promoter---Fig.3

Table S7: List of sgRNAs related to Figure 7D

| Target | sgRNA_sequence |
|---|---|
| chr4_16140800_16141398 | AAGCACTATTCTGAAAATGT |
| chr4_16140800_16141398 | CACTATTCTGAAAATGTTGG |
| chr4_16140800_16141398 | GAAAGTTTTCTCAGATTCCC |
| chr4_16140800_16141398 | TCAGATTCCCTGGCATCTCC |
| chr4_16140800_16141398 | CAGATTCCCTGGCATCTCCT |
| chr4_16140800_16141398 | CTCCTGGGCTTCTCCATAAA |
| chr4_16140800_16141398 | CTTCTCCATAAATGGAAAGA |
| chr4_16140800_16141398 | GTTCAGCTTCACTGAAGTCT |
| chr4_16140800_16141398 | GTCTTGGATTCTTGTCCCCC |
| chr4_16140800_16141398 | TCTTGGATTCTTGTCCCCCA |
| chr4_16140800_16141398 | CCAGGGACTGAAACCAAACC |
| chr4_16140800_16141398 | TGAAACCAAACCTGGAGCCC |
| chr4_16140800_16141398 | CCTGGAGCCCAGGTCACTAC |
| chr4_16140800_16141398 | CTGGAGCCCAGGTCACTACA |
| chr4_16140800_16141398 | GGGAAATGATCCAGACCTCA |
| chr4_16140800_16141398 | GAATTTCAAAACACTTACTC |
| chr4_16140800_16141398 | ACTCAGGCAACAGCTTTACC |
| chr4_16140800_16141398 | CAACAGCTTTACCTGGAAGC |
| chr4_16140800_16141398 | ACCTGGAAGCTGGCCAACAA |
| chr4_16140800_16141398 | GGCCAACAAAGGCCATATAA |
| chr4_16140800_16141398 | ATGGACAGAGCAACATTCTG |
| chr4_16140800_16141398 | ACAGAGCAACATTCTGTGGT |
| chr4_16140800_16141398 | CAGAGCAACATTCTGTGGTT |
| chr4_16140800_16141398 | AGAGCAACATTCTGTGGTTG |
| chr4_16140800_16141398 | TTGGGGAAATGCTTTCAGAG |
| chr4_16140800_16141398 | CTTTCAGAGAGGCAAGCCTT |
| chr4_16140800_16141398 | TTTCAGAGAGGCAAGCCTTT |
| chr4_16140800_16141398 | AGCCTTTGGGACTGTAGATT |
| chr4_16140800_16141398 | GTAAATAGAGTAGAGCTTCC |
| chr4_16140800_16141398 | AGAGTAGAGCTTCCTGGTAG |
| chr4_16140800_16141398 | GAGTAGAGCTTCCTGGTAGA |
| chr4_16140800_16141398 | GAGCTTCCTGGTAGAGGGCT |
| chr4_16140800_16141398 | GTAGAGGGCTTGGTTGCTCA |
| chr4_16140800_16141398 | ACAGTGTGTGTCTTTGTCCA |
| chr4_16140800_16141398 | TACTCTATCATCAGAGTTAA |
| chr4_16140800_16141398 | CTATCATCAGAGTTAATGGC |
| chr4_16140800_16141398 | GGAAACAGAAAGTCCCCTAG |
| chr4_16140800_16141398 | GAAACAGAAAGTCCCCTAGT |
| chr4_16140800_16141398 | AAAGTCCCCTAGTGGGTTTT |
| chr4_16140800_16141398 | GAGAATTAATACAGTTAATT |
| chr4_16140800_16141398 | GAAATCATCTGAAAGCTGTC |
| chr4_16140800_16141398 | TCTGAAAGCTGTCAGGAAGT |
| chr4_16140800_16141398 | CTGAAAGCTGTCAGGAAGTT |
| chr4_16140800_16141398 | AAAGCTGTCAGGAAGTTGGG |
| chr4_16140800_16141398 | GAAGTTGGGTGGCTCTAGAC |
| chr4_16140800_16141398 | AAGTTGGGTGGCTCTAGACT |
| chr4_16140800_16141398 | TGGCTCTAGACTGGGTCTCT |
| chr4_16140800_16141398 | TAGTGCTTTGGCTCTCCTTG |
| chr4_16140800_16141398 | ACATTTTCAGAATAGTGCTT |
| chr4_16140800_16141398 | GAGAAGCCCAGGAGATGCCA |
| chr4_16140800_16141398 | GGAGAAGCCCAGGAGATGCC |
| chr4_16140800_16141398 | TTCCATTTATGGAGAAGCCC |
| chr4_16140800_16141398 | CTGAGCCTTCTTTCCATTTA |
| chr4_16140800_16141398 | GGTTTGGTTTCAGTCCCTGG |
| chr4_16140800_16141398 | AGGTTTGGTTTCAGTCCCTG |
| chr4_16140800_16141398 | CAGGTTTGGTTTCAGTCCCT |
| chr4_16140800_16141398 | CCAGGTTTGGTTTCAGTCCC |
| chr4_16140800_16141398 | AGTGACCTGGGCTCCAGGTT |
| chr4_16140800_16141398 | CCTGTAGTGACCTGGGCTCC |
| chr4_16140800_16141398 | TCATTTCCCTGTAGTGACCT |
| chr4_16140800_16141398 | ATCATTTCCCTGTAGTGACC |
| chr4_16140800_16141398 | TTTTGAAATTCCTTGAGGTC |
| chr4_16140800_16141398 | AAGTGTTTTGAAATTCCTTG |
| chr4_16140800_16141398 | GCCTTTGTTGGCCAGCTTCC |
| chr4_16140800_16141398 | GTCCATTATATGGCCTTTGT |
| chr4_16140800_16141398 | ATGTTGCTCTGTCCATTATA |
| chr4_16140800_16141398 | ATCCTAATCTACAGTCCCAA |
| chr4_16140800_16141398 | ACTCTATTTACACTGGTATT |
| chr4_16140800_16141398 | AAGCTCTACTCTATTTACAC |
| chr4_16140800_16141398 | GAGCAACCAAGCCCTCTACC |
| chr4_16140800_16141398 | CTGGACTGTGGAGCTGACCA |
| chr4_16140800_16141398 | GAGTAACTTACTCTGGACTG |
| chr4_16140800_16141398 | GATGATAGAGTAACTTACTC |
| chr4_16140800_16141398 | GTATACCTAAAACCCACTAG |
| chr4_16140800_16141398 | TGTATACCTAAAACCCACTA |
| chr4_16140800_16141398 | CTGTATACCTAAAACCCACT |
| chr4_16140800_16141398 | GACAGCTTTCAGATGATTTC |
| chr4_16033958_16034556 | GAATTGAGTAATTAAATACA |
| chr4_16033958_16034556 | TAATTAAATACATGGCCAAG |
| chr4_16033958_16034556 | ATACATGGCCAAGTGGTTAC |
| chr4_16033958_16034556 | GAAGATGTCTTTCAACCCCT |
| chr4_16033958_16034556 | ATGCTAGTTTGAGCTGCAAA |
| chr4_16033958_16034556 | TGGTTTAATTTTAACCCCAA |
| chr4_16033958_16034556 | GTATTTTGTTTGTTTTTAAA |
| chr4_16033958_16034556 | TTAAATGGATCAATAAATCT |
| chr4_16033958_16034556 | ATGGATCAATAAATCTTGGC |
| chr4_16033958_16034556 | TGCATAACAATCTACATATA |
| chr4_16033958_16034556 | GTTAAAGATTCAAAGCACTT |
| chr4_16033958_16034556 | AGATTCAAAGCACTTTGGAT |
| chr4_16033958_16034556 | TCAAAGCACTTTGGATTGGC |
| chr4_16033958_16034556 | CAAACAATTACTCAAGAATC |
| chr4_16033958_16034556 | TGTATCTTAATTTAGAAAAA |
| chr4_16033958_16034556 | AAAGGCTCAGACTCTAAGTC |
| chr4_16033958_16034556 | AAGTCAGGCACGTTAAACAC |
| chr4_16033958_16034556 | TTAAACACAGGCCAGTCACA |
| chr4_16033958_16034556 | AGGTTCCTCCTCTGATACAC |
| chr4_16033958_16034556 | GATTAGAAATAATGTGACAA |
| chr4_16033958_16034556 | ACTGAGTAGACACTTACTAA |
| chr4_16033958_16034556 | GTGCCTCTCATGCCATGCCC |
| chr4_16033958_16034556 | AGGAGCCAGATCCACAGTGA |
| chr4_16033958_16034556 | CCACAGTGAAGGACCGACCT |

Table S7: List of sgRNAs related to Figure S7B-S7C

| | |
|---|---|
| chr1:31784892-31785299 | CTCCATCACCGCTAACCCGC |
| chr1:31784892-31785299 | CTACAGGACACAGAAGGACT |
| chr1:31784892-31785299 | CCGCTAACCCGCTGGCTGCC |
| chr1:31784892-31785299 | GCCTGTCAGCCTCCACCATC |
| chr1:31784892-31785299 | CTAACCCGCTGGCTGCCAGG |
| chr1:31784892-31785299 | ACACCCCAGCCTACTTCTGG |
| chr1:31784892-31785299 | GGCTCATCTCAGAGAGTGAT |
| chr1:31784892-31785299 | CCAGAGGGGTATCCTGCGAC |
| chr1:31784892-31785299 | CCAGGATAGAGCCTGTCCCA |
| chr1:31784892-31785299 | AGCCTGTGCACACACTGGCT |
| chr1:31784892-31785299 | TAACCCGCTGGCTGCCAGGC |
| chr1:31784892-31785299 | TAGCCAGTGTGTGCACAGGC |
| chr1:31784892-31785299 | AGATGAGCCGAGTGCCAAGC |
| chr1:31784892-31785299 | GCCAGATGGTGGAGGCTGAC |
| chr1:31784892-31785299 | AAGGTGGGGGGACCTGTCGC |
| chr1:31784892-31785299 | GGACAGGCTCTATCCTGGCC |
| chr1:31784892-31785299 | ACAGGCTCTATCCTGGCCTG |
| chr1:31784892-31785299 | CTCAGAGAGTGATGGGCAAG |
| chr1:31784892-31785299 | CGGCTCATCTCAGAGAGTGA |
| chr1:31784892-31785299 | AGACATCTACAGGACACAGA |
| chr1:31784892-31785299 | TCAGCCTCCACCATCTGGCC |
| chr1:31784892-31785299 | GTGATGGGCAAGTGGACATG |
| chr1:31784892-31785299 | TGTGCACACACTGGCTAGGT |
| chr1:31784892-31785299 | TGGAGGCTGACAGGCCCCAA |
| chr1:31784892-31785299 | AGCCTACTTCTGGCGGAGCC |
| chr1:31784892-31785299 | TGAGTGTGAATGAGAGACCT |
| chr1:31784892-31785299 | GAGCCGAGTGCCAAGCCGGA |
| chr1:31784892-31785299 | GGTGATGGAGAGCTCCCTTG |
| chr1:31784892-31785299 | GGCCTGGCTCCGCCAGAAGT |
| chr1:31784892-31785299 | GACCTAGCCAGTGTGTGCAC |
| chr1:31784892-31785299 | TGAATGAGAGACCTTGGGAC |
| chr1:31784892-31785299 | TCCCCAGCCTGGCCAGATGG |
| chr1:31784892-31785299 | GACAGGCTCTATCCTGGCCT |
| chr1:31784892-31785299 | CGGTGATGGAGAGCTCCCTT |
| chr1:31784892-31785299 | TGGCTCCGCCAGAAGTAGGC |
| chr1:31784892-31785299 | GTGCACACACTGGCTAGGTC |
| chr1:31784892-31785299 | CCTTGGGACAGGCTCTATCC |
| chr1:31784892-31785299 | CTGTCGCAGGATACCCCTCT |
| chr1:31784892-31785299 | AGCCAGCGGGTTAGCGGTGA |
| chr1:31784892-31785299 | TGCCAAGCCGGAGGGCCCAG |
| chr1:31784892-31785299 | CCTGGCAGCCAGCGGGTTAG |
| chr1:31784892-31785299 | CCAGCCTGGCCAGATGGTGG |
| chr1:31784892-31785299 | TGAGCCGAGTGCCAAGCCGG |
| chr1:31784892-31785299 | GGATACCCCTCTGGGCCCTC |
| chr1:31784892-31785299 | CCTGTCGCAGGATACCCCTC |
| chr1:31784892-31785299 | TGCACACACTGGCTAGGTCG |
| chr1:31784892-31785299 | TGGCCAGCCTGTGCACACAC |
| chr1:31784892-31785299 | GAGTGTGAATGAGAGACCTT |
| chr1:31784892-31785299 | GCTCCGCCAGAAGTAGGCTG |
| chr1:31784892-31785299 | TTTACACCCCAGCCTACTTC |
| chr1:31784892-31785299 | GGCTCCGCCAGAAGTAGGCT |
| chr1:31784892-31785299 | AGTAGGCTGGGGTGTAAATA |
| chr1:31784892-31785299 | GCGGTGATGGAGAGCTCCCT |
| chr1:31784892-31785299 | ATTCACACTCAGACATCTAC |

| | |
|---|---|
| chr1:202283330-202283682 | AGCCATACCACCTGGGGAGG |
| chr1:202283330-202283682 | GCCATACCACCTGGGGAGGA |
| chr1:202283330-202283682 | GGGCCGCTTTCCCCACTTAG |
| chr1:202283330-202283682 | CCATACCACCTGGGGAGGAG |
| chr1:202283330-202283682 | AGGGGCAGCCAGATGTGGCA |
| chr1:202283330-202283682 | AACCCCTGCGGCCCAAGCTG |
| chr1:202283330-202283682 | CTGCGGCCCAAGCTGCGGTT |
| chr1:202283330-202283682 | AACCGCAGCTTGGGCCGCAG |
| chr1:202283330-202283682 | CTCCCCAGGTGGTATGGCTG |
| chr1:202283330-202283682 | GCAGAGATTTATGGGGCCTG |
| chr1:202283330-202283682 | CCTCCCCAGGTGGTATGGCT |
| chr1:202283330-202283682 | GTCTAAGTGGCAGAGATTTA |
| chr1:202283330-202283682 | CTCCAGCCAAACCGCAGCTT |
| chr1:202283330-202283682 | TTGGCACGCTTTGCAGTCTC |
| chr1:202283330-202283682 | CGGCCCCCAGCCATACCACC |
| chr1:202283330-202283682 | TCTAAGTGGCAGAGATTTAT |
| chr1:202283330-202283682 | TCCCCAGGTGGTATGGCTGG |
| chr1:202283330-202283682 | CAGAGATTTATGGGGCCTGA |
| chr1:202283330-202283682 | GGCCCCCAGCCATACCACCT |
| chr1:202283330-202283682 | CAGGGGCAGCCAGATGTGGC |
| chr1:202283330-202283682 | GGAGATGAGAGAAGTCTAAG |
| chr1:202283330-202283682 | GCCCCCAGCCATACCACCTG |
| chr1:202283330-202283682 | CTAAGTGGCAGAGATTTATG |
| chr1:202283330-202283682 | ATGTGAGCTTCTCACTGGAA |
| chr1:202283330-202283682 | GAGGGAAGATCAAACCCCTG |
| chr1:202283330-202283682 | AAACCGCAGCTTGGGCCGCA |
| chr1:202283330-202283682 | TTTGATCTTCCCTCTGCTGC |
| chr1:202283330-202283682 | GGTGCGGGCATGCACACACG |
| chr1:202283330-202283682 | TGGAAGGGTATTTCTCCCTC |
| chr1:202283330-202283682 | CGTGTGTGCATGCCCGCACC |
| chr1:202283330-202283682 | CCCCTCCTCCCCAGGTGGTA |
| chr1:202283330-202283682 | GCTCCAGCCAAACCGCAGCT |
| chr1:202283330-202283682 | ACTCCTCTAAGTGGGGAAAG |
| chr1:202283330-202283682 | CTCTCATCTCCGGCAGCAGA |
| chr1:202283330-202283682 | TCTCTCATCTCCGGCAGCAG |
| chr1:202283330-202283682 | ACTCCTCCCCCTGCCACATC |
| chr1:202283330-202283682 | CAAACCGCAGCTTGGGCCGC |
| chr1:202283330-202283682 | TCCTCCCCAGGTGGTATGGC |
| chr1:202283330-202283682 | CCTCTGCAGACTCCTCTAAG |
| chr1:202283330-202283682 | CCCAGCCATACCACCTGGGG |
| chr1:202283330-202283682 | CATGTGAGCTTCTCACTGGA |
| chr1:202283330-202283682 | TCTGCAGACTCCTCTAAGTG |
| chr1:202283330-202283682 | CTCTGCAGACTCCTCTAAGT |
| chr1:202283330-202283682 | GGCCCAAGCTGCGGTTTGGC |
| chr1:202283330-202283682 | ACTTAGACTTCTCTCATCTC |
| chr1:202283330-202283682 | CCACTTAGAGGAGTCTGCAG |

| | |
|---|---|
| chr4_16033958_16034556 | TTGGCAGTCCGTGTCTCAGA |
| chr4_16033958_16034556 | CAGACGGTTCCTGAGTTCCT |
| chr4_16033958_16034556 | GACATCTTCCAGTAACCACT |
| chr4_16033958_16034556 | GCATTTAGACATCTTCCTAG |
| chr4_16033958_16034556 | AGCATTTAGACATCTTCCTA |
| chr4_16033958_16034556 | TAGCATTTAGACATCTTCCT |
| chr4_16033958_16034556 | AAACAAACAAAATACCTTTG |
| chr4_16033958_16034556 | AAAACAAACAAAATACCTTT |
| chr4_16033958_16034556 | AAAAACAAACAAAATACCTT |
| chr4_16033958_16034556 | CAGAGGAGGAACCTTGTGAC |
| chr4_16033958_16034556 | TATTTCCAGTGTATCAGAGG |
| chr4_16033958_16034556 | TATTATTTCCAGTGTATCAG |
| chr4_16033958_16034556 | ATTTTCACAATGACTTGACA |
| chr4_16033958_16034556 | GTAAGTGTCTACTCAGTGCT |
| chr4_16033958_16034556 | GCTCCTGGGCATGGCATGAG |
| chr4_16033958_16034556 | GTGGATCTGGCTCCTGGGCA |
| chr4_16033958_16034556 | TCACTGTGGATCTGGCTCCT |
| chr4_16033958_16034556 | TTCACTGTGGATCTGGCTCC |
| chr4_16033958_16034556 | TCGGTCCTTCACTGTGGATC |
| chr4_16033958_16034556 | CCAAGGTCGGTCCTTCACTG |
| chr4_16033958_16034556 | GAGACACGGACTGCCAAGGT |
| chr4_16033958_16034556 | GTCTGAGACACGGACTGCCA |
| chr4_16033958_16034556 | CTCAGGAACCGTCTGAGACA |
| chr4_16033958_16034556 | TGTGCTGTTCCTAGGAACTC |
| chr4_16033958_16034556 | CTAGAAATTGTGCTGTTCCT |
| chr4_16123302_16123900 | CCTCATATCCAGTAAGAGGC |
| chr4_16123302_16123900 | CTCATATCCAGTAAGAGGCA |
| chr4_16123302_16123900 | CCAGTAAGAGGCAGGGATTT |
| chr4_16123302_16123900 | GCACTGAGCTGCTCCTCTCG |
| chr4_16123302_16123900 | GCTCCTCTCGAGGCAGTGAG |
| chr4_16123302_16123900 | TCTCGAGGCAGTGAGAGGTT |
| chr4_16123302_16123900 | CTCGAGGCAGTGAGAGGTTT |
| chr4_16123302_16123900 | TCGAGGCAGTGAGAGGTTTG |
| chr4_16123302_16123900 | AGTGAGAGGTTTGGGGTCCC |
| chr4_16123302_16123900 | GTGAGAGGTTTGGGGTCCCT |
| chr4_16123302_16123900 | TGAGAGGTTTGGGGTCCCTG |
| chr4_16123302_16123900 | CCCTGGGGTTTTCCCACACC |
| chr4_16123302_16123900 | GGTTTTCCCACACCTGGCAC |
| chr4_16123302_16123900 | TCTCGCGCTGTAAACAGAGA |
| chr4_16123302_16123900 | ACAGAGAAGGCTCTGTAATT |
| chr4_16123302_16123900 | GATTATTCTCCACTCTTCTG |
| chr4_16123302_16123900 | TTCTCCACTCTTCTGTGGTT |
| chr4_16123302_16123900 | AATGCTGCCAATTATCTCCC |
| chr4_16123302_16123900 | TTATCTCCCTGGTGCTTACA |
| chr4_16123302_16123900 | TATCTCCCTGGTGCTTACAT |
| chr4_16123302_16123900 | ACAGAGCACCTGCCTCTCTG |
| chr4_16123302_16123900 | CAGAGCACCTGCCTCTCTGA |
| chr4_16123302_16123900 | AGAGCACCTGCCTCTCTGAG |
| chr4_16123302_16123900 | GAGCACCTGCCTCTCTGAGG |
| chr4_16123302_16123900 | TGCCTCTCTGAGGGGGTGCC |
| chr4_16123302_16123900 | GGGTGCCAGGATCCCATTCC |
| chr4_16123302_16123900 | TCCCATTCCTGGCAGCCAAG |
| chr4_16123302_16123900 | CTGGCAGCCAAGAGGATGAA |
| chr4_16123302_16123900 | TTGTTTACACTGCAAAGCTG |
| chr4_16123302_16123900 | TGCAAAGCTGAGGAAAGCCT |
| chr4_16123302_16123900 | TGAGGAAAGCCTTGGTTGTG |
| chr4_16123302_16123900 | GCACCAGCTCTTTCAACCCG |
| chr4_16123302_16123900 | ACCTCAGCCTCTCACACCTC |
| chr4_16123302_16123900 | CCTCAGCCTCTCACACCTCT |
| chr4_16123302_16123900 | CTCACACCTCTGGGAGCCAG |
| chr4_16123302_16123900 | CACCTCTGGGAGCCAGAGGC |
| chr4_16123302_16123900 | ACCTCTGGGAGCCAGAGGCT |
| chr4_16123302_16123900 | ATTTCCCCCTTTCACAGACA |
| chr4_16123302_16123900 | GACAAGGAAACCCAAGCTCA |
| chr4_16123302_16123900 | AAAGTCCCCCAGCTAGTTAC |
| chr4_16123302_16123900 | TAGTTACTGGCAGAGCTGAG |
| chr4_16123302_16123900 | AGCACTTGCTGTTTGCACTG |
| chr4_16123302_16123900 | CCTGCCTCTTACTGGATATG |
| chr4_16123302_16123900 | CCAAAATCCCTGCCTCTTAC |
| chr4_16123302_16123900 | CAGTGCAGTGGAAAATTTCA |
| chr4_16123302_16123900 | AGAGGAGCAGCTCAGTGCAG |
| chr4_16123302_16123900 | AAACCTCTCACTGCCTCGAG |
| chr4_16123302_16123900 | CCAGGTGTGGGAAAACCCCA |
| chr4_16123302_16123900 | GCCAGGTGTGGGAAAACCCC |
| chr4_16123302_16123900 | GTGCAGCCTGTGCCAGGTGT |
| chr4_16123302_16123900 | AGTGCAGCCTGTGCCAGGTG |
| chr4_16123302_16123900 | GCGAGAGTGCAGCCTGTGCC |
| chr4_16123302_16123900 | GTTACCTAACCACAGAAGAG |
| chr4_16123302_16123900 | GTAAGCACCAGGGAGATAAT |
| chr4_16123302_16123900 | TCTGTCCCATGTAAGCACCA |
| chr4_16123302_16123900 | CTCTGTCCCATGTAAGCACC |
| chr4_16123302_16123900 | TGGCACCCCCTCAGAGAGGC |
| chr4_16123302_16123900 | ATCCTGGCACCCCCTCAGAG |
| chr4_16123302_16123900 | GGCTGCCAGGAATGGGATCC |
| chr4_16123302_16123900 | TCCTCTTGGCTGCCAGGAAT |
| chr4_16123302_16123900 | ATCCTCTTGGCTGCCAGGAA |
| chr4_16123302_16123900 | CTTTCATCCTCTTGGCTGCC |
| chr4_16123302_16123900 | TGCTGGGCCTTTCATCCTCT |
| chr4_16123302_16123900 | AAACAAGGAAACACACTGCT |
| chr4_16123302_16123900 | TAAACAAGGAAACACACTGC |
| chr4_16123302_16123900 | TCAGCTTTGCAGTGTAAACA |
| chr4_16123302_16123900 | GAGCTGGTGCCTCACAACCA |
| chr4_16123302_16123900 | GCTCCTCGGGTTGAAAGAGC |
| chr4_16123302_16123900 | AGGTGGGTGAAGGGCTCCTC |
| chr4_16123302_16123900 | GAGGTGGGTGAAGGGCTCCT |
| chr4_16123302_16123900 | GAGAGGCTGAGGTGGGTGAA |
| chr4_16123302_16123900 | TGAGAGGCTGAGGTGGGTGA |
| chr4_16123302_16123900 | GAGGTGTGAGAGGCTGAGGT |
| chr4_16123302_16123900 | AGAGGTGTGAGAGGCTGAGG |
| chr4_16123302_16123900 | CCCAGAGGTGTGAGAGGCTG |
| chr4_16123302_16123900 | CTGGCTCCCAGAGGTGTGAG |
| chr4_16123302_16123900 | GCCCAGCCTCTGGCTCCCAG |
| chr4_16123302_16123900 | GGAAATAAAAGCCCAGCCTC |
| chr4_16123302_16123900 | GTTTCCTTGTCTGTGAAAGG |
| chr4_16123302_16123900 | GGTTTCCTTGTCTGTGAAAG |

| | |
|---|---|
| chr4_16123302_16123900 | GGGTTTCCTTGTCTGTGAAA |
| chr4_16123302_16123900 | TGGGTTTCCTTGTCTGTGAA |
| chr4_16123302_16123900 | TTTTCTGCTTCCTTGAGCTT |
| chr4_16123302_16123900 | TTTTTCTGCTTCCTTGAGCT |
| chr4_16123302_16123900 | GTAACTAGCTGGGGGACTTT |
| chr4_16123302_16123900 | AGTAACTAGCTGGGGGACTT |
| chr4_16123302_16123900 | CTCTGCCAGTAACTAGCTGG |
| chr4_16123302_16123900 | GCTCTGCCAGTAACTAGCTG |
| chr4_16123302_16123900 | AGCTCTGCCAGTAACTAGCT |
| chr4_16123302_16123900 | CAGCTCTGCCAGTAACTAGC |
| chr4_16123302_16123900 | TTCTATGGAGAGACAACAGA |
| chr4_16123302_16123900 | TGGTTACTAGATGAATTCTA |
| chr4_16123302_16123900 | AAACAGCAAGTGCTCAGTAA |
| chr4_16067392_16067990 | AGATAACAGCAGTTAGCTCC |
| chr4_16067392_16067990 | CTCGCCTAAATCCCTGAAAG |
| chr4_16067392_16067990 | TCGCCTAAATCCCTGAAAGC |
| chr4_16067392_16067990 | GAAAGCGGGCACACAGTGAG |
| chr4_16067392_16067990 | TGCTTGCCATTCTGACTGTT |
| chr4_16067392_16067990 | TGCCATTCTGACTGTTAGGA |
| chr4_16067392_16067990 | GCCCAGCAGAGTGACTGACA |
| chr4_16067392_16067990 | CAGCAGAGTGACTGACATGG |
| chr4_16067392_16067990 | CAGAGTGACTGACATGGAGG |
| chr4_16067392_16067990 | GTGACTGACATGGAGGAGGA |
| chr4_16067392_16067990 | TCTGTGTGTGACCTAATTTG |
| chr4_16067392_16067990 | GTGTGACCTAATTTGTGGCT |
| chr4_16067392_16067990 | GCATGAGTCACATGTTCTAA |
| chr4_16067392_16067990 | TGTTCTAAAGGCGACAATGC |
| chr4_16067392_16067990 | GGCTTTGTCTGAATTAAGTA |
| chr4_16067392_16067990 | AAGTAAGGACACGCAGATGT |
| chr4_16067392_16067990 | TGTTGGAGCCTGATCCTACG |
| chr4_16067392_16067990 | AGCCTGATCCTACGTGGACA |
| chr4_16067392_16067990 | GCCTGATCCTACGTGGACAT |
| chr4_16067392_16067990 | TGGGCATAGACACGTCGATT |
| chr4_16067392_16067990 | TCACTCAGCCGCCCCTTTCC |
| chr4_16067392_16067990 | CACTCAGCCGCCCCTTTCCT |
| chr4_16067392_16067990 | ACCGCTTCTCTCTGCCTCTG |
| chr4_16067392_16067990 | TCTCTCTGCCTCTGAGGCCA |
| chr4_16067392_16067990 | CTGAGGCCATGGTGACAGTG |
| chr4_16067392_16067990 | TGAGGCCATGGTGACAGTGA |
| chr4_16067392_16067990 | AGTGAGGGCCCACAGCAGCC |
| chr4_16067392_16067990 | CAGGCTCACACAGACGACCC |
| chr4_16067392_16067990 | ACCCCGGCCACGTGACTCAA |
| chr4_16067392_16067990 | CACGTGACTCAATGGCTCCA |
| chr4_16067392_16067990 | ATGGCTCCACGGTTTATGAT |
| chr4_16067392_16067990 | TCCACGGTTTATGATTGGTT |
| chr4_16067392_16067990 | CCACGGTTTATGATTGGTTC |
| chr4_16067392_16067990 | GGTGAGCCAGAAGCCCTCCC |
| chr4_16067392_16067990 | GTGAGCCAGAAGCCCTCCCT |
| chr4_16067392_16067990 | CCTCCCTGGGAATTGAGAAT |
| chr4_16067392_16067990 | CTCCCTGGGAATTGAGAATT |
| chr4_16067392_16067990 | CCAGCCACTCTCCAAGAAGC |
| chr4_16067392_16067990 | TTTTACAGAAGAGGAAGTCG |
| chr4_16067392_16067990 | GTTATCTCATTTTACAGAAG |
| chr4_16067392_16067990 | TTAGGCGAGTCAACAGCACC |
| chr4_16067392_16067990 | GTGCCCGCTTTCAGGGATTT |
| chr4_16067392_16067990 | TCACTGTGTGCCCGCTTTCA |
| chr4_16067392_16067990 | CTCACTGTGTGCCCGCTTTC |
| chr4_16067392_16067990 | TGCCATCCTAACAGTCAGAA |
| chr4_16067392_16067990 | TCCATGTCAGTCACTCTGCT |
| chr4_16067392_16067990 | CTCCATGTCAGTCACTCTGC |
| chr4_16067392_16067990 | ACAAATTAGGTCACACACAG |
| chr4_16067392_16067990 | AATGGGCCTAGCCACAAATT |
| chr4_16067392_16067990 | ATGTGACTCATGCTGGCAAT |
| chr4_16067392_16067990 | CATGTGACTCATGCTGGCAA |
| chr4_16067392_16067990 | TTAGAACATGTGACTCATGC |
| chr4_16067392_16067990 | GCCCATGTCCACGTAGGATC |
| chr4_16067392_16067990 | GTCTATGCCCATGTCCACGT |
| chr4_16067392_16067990 | AGCGGTACCCAGGAAAGGGG |
| chr4_16067392_16067990 | AGAAGCGGTACCCAGGAAAG |
| chr4_16067392_16067990 | GAGAAGCGGTACCCAGGAAA |
| chr4_16067392_16067990 | AGAGAAGCGGTACCCAGGAA |
| chr4_16067392_16067990 | GGCAGAGAGAAGCGGTACCC |
| chr4_16067392_16067990 | GCCTCAGAGGCAGAGAGAAG |
| chr4_16067392_16067990 | CACTGTCACCATGGCCTCAG |
| chr4_16067392_16067990 | GTGGGCCCTCACTGTCACCA |
| chr4_16067392_16067990 | TGTGTGAGCCTGGCTGCTGT |
| chr4_16067392_16067990 | CTGTGTGAGCCTGGCTGCTG |
| chr4_16067392_16067990 | CGGGGTCGTCTGTGTGAGCC |
| chr4_16067392_16067990 | GCCATTGAGTCACGTGGCCG |
| chr4_16067392_16067990 | AGCCATTGAGTCACGTGGCC |
| chr4_16067392_16067990 | GAGCCATTGAGTCACGTGGC |
| chr4_16067392_16067990 | CGTGGAGCCATTGAGTCACG |
| chr4_16067392_16067990 | CCCGAACCAATCATAAACCG |
| chr4_16067392_16067990 | CAATTCCCAGGGAGGGCTTC |
| chr4_16067392_16067990 | CAATTCTCAATTCCCAGGGA |
| chr4_16067392_16067990 | CCAATTCTCAATTCCCAGGG |
| chr4_16067392_16067990 | ATCCCAATTCTCAATTCCCA |
| chr4_16067392_16067990 | TATCCCAATTCTCAATTCCC |
| chr4_16067392_16067990 | CAGCTTCTTGGAGAGTGGCT |
| chr4_16067392_16067990 | CCAGCTTCTTGGAGAGTGGC |
| chr4_16067392_16067990 | CAGGCCAGCTTCTTGGAGAG |

**Table 1: SOX9 interacting proteins detected by RIME (related to Figure 1A and Extended data Fig. 1A)**

| Rank | RIME-Interacting protein Gene Symbol | Pathway / Group (GENETS Analysis) | Relative binding score | | | |
|---|---|---|---|---|---|---|
| | | | GFP (+DOX) | $SOX9^{FL}$ (No DOX) | $SOX9^{FL}$ (+DOX) | $SOX9^{\Delta C}$ (+DOX) |
| 1 | SMARCC1 | BAF complex | 0 | 0 | 3.5 | 0 |
| 2 | OGT | Histone deacetylase | 0 | 0 | 2.5 | 0 |
| 3 | TRIM24 | Others | 0 | 0 | 2 | 0 |
| 4 | CBX5 | Histone deacetylase | 0 | 0 | 2 | 0 |
| 5 | KDM3B | Histone deacetylase | 0 | 0 | 2 | 0 |
| 6 | CYB5R3 | Others | 0 | 0 | 1.5 | 0 |
| 7 | DDX23 | Spliceosome complex | 0 | 0 | 1.5 | 0 |
| 8 | CHD5 | BAF complex | 0 | 0 | 1.5 | 0 |
| 9 | PTBP3 | Others | 0 | 0 | 1.5 | 0 |
| 10 | BRD2 | Others | 0 | 0 | 1.5 | 0 |
| 11 | IRF2BP1 | Others | 0 | 0 | 1 | 0 |
| 12 | ZMYM3 | Others | 0 | 0 | 1 | 0 |
| 13 | PPP1R8 | Spliceosome complex | 0 | 0 | 1 | 0 |
| 14 | ZNF687 | Others | 0 | 0 | 1 | 0 |
| 15 | CNDP2 | Others | 0 | 0 | 1 | 0 |
| 16 | ACOT13 | Others | 0 | 0 | 1 | 0 |
| 17 | MBD2 | BAF complex | 0 | 0 | 1 | 0 |
| 18 | PPIE | Spliceosome complex | 0 | 0 | 1 | 0 |
| 19 | NOP58 | Others | 0 | 0 | 1 | 0 |
| 20 | QRICH1 | Others | 0 | 0 | 1 | 0 |
| 21 | SMARCB1 | BAF complex | 0 | 0 | 1 | 0 |
| 22 | DHX38 | Spliceosome complex | 0 | 0 | 1 | 0 |
| 23 | HINT1 | Histone deacetylase | 0 | 0.5 | 1 | 0 |
| 24 | RBBP6 | Others | 0 | 0.5 | 1 | 0 |
| 25 | MRPL12 | Others | 0 | 0.5 | 1 | 0 |
| 26 | FAM133B | Others | 0 | 0.5 | 1 | 0 |
| 27 | SMARCC2 | BAF complex | 0 | 0 | 4.5 | 1 |
| 28 | SATB2 | Histone deacetylase | 0 | 0 | 2 | 0.5 |
| 29 | TCF20 | Others | 0 | 0 | 2 | 0.5 |
| 30 | WRNIP1 | DNA repair | 0 | 0 | 2 | 0.5 |
| 31 | WDHD1 | Others | 0 | 0 | 3 | 1 |
| 32 | TCEA1 | Transcription regulation | 0 | 0 | 3 | 1 |
| 33 | SON | Others | 0 | 1 | 1.5 | 0.5 |
| 34 | NIPBL | Histone acetylation | 0 | 0 | 1.5 | 0.5 |
| 35 | EP300 | Histone acetylation | 0 | 0 | 1.5 | 0.5 |
| 36 | HTATSF1 | Spliceosome complex | 0 | 0.5 | 2.5 | 1 |
| 37 | NCOR1 | Histone deacetylase | 0 | 0 | 2.5 | 1 |
| 38 | ADNP | Transcription regulation | 0 | 0 | 3.5 | 1.5 |
| 39 | DHX16 | Spliceosome complex | 0 | 0 | 2 | 1 |
| 40 | DEK | BAF complex | 0 | 0.5 | 2 | 1 |
| 41 | TPI1 | Others | 0 | 0.5 | 2 | 1 |
| 42 | CD2BP2 | Spliceosome complex | 0 | 0 | 2 | 1 |
| 43 | SMARCD1 | BAF complex | 0 | 0 | 2 | 1 |
| 44 | CBR1 | Others | 0 | 0.5 | 1 | 0.5 |
| 45 | CTBP1 | Histone acetylation | 0 | 0 | 1 | 0.5 |
| 46 | PPIL1 | Spliceosome complex | 0 | 0 | 1 | 0.5 |
| 47 | NOSIP | Others | 0 | 0 | 1 | 0.5 |
| 48 | POLR2A | Others | 0 | 0 | 1 | 0.5 |
| 49 | TFAP4 | Transcription regulation | 0 | 0 | 1 | 0.5 |
| 50 | VGLL4 | Others | 0 | 0 | 1 | 0.5 |
| 51 | PREP | Others | 0 | 0.5 | 1 | 0.5 |
| 52 | HNRNPM | Spliceosome complex | 0 | 0.5 | 1 | 0.5 |
| 53 | USP48 | Others | 0 | 0 | 1 | 0.5 |
| 54 | TRIM33 | Others | 0 | 0 | 1 | 0.5 |

| | | | | | | |
|---|---|---|---|---|---|---|
| 55 | DPF2 | BAF complex | 0 | 0 | 1 | 0.5 |
| 56 | CSTF3 | Others | 0 | 0 | 1 | 0.5 |
| 57 | PPP4R3A | Others | 0 | 0 | 1 | 0.5 |
| 58 | YY1 | BAF complex | 0 | 0 | 1 | 0.5 |
| 59 | PPM1A | Others | 0 | 0 | 1 | 0.5 |
| 60 | PRPF8 | Spliceosome complex | 0 | 0 | 5.5 | 3 |
| 61 | ANP32E | BAF complex | 0 | 0 | 2.5 | 1.5 |
| 62 | XRN2 | Others | 0 | 0 | 1.5 | 1 |
| 63 | CBX1 | heterochromatin | 0 | 0 | 1.5 | 1 |
| 64 | POP1 | Others | 0 | 0 | 1.5 | 1 |
| 65 | NUDT21 | Others | 0 | 0 | 2 | 1.5 |
| 66 | GMPS | Others | 0 | 0 | 2 | 1.5 |
| 67 | HAT1 | Histone acetylation | 0 | 0.5 | 2.5 | 2 |
| 68 | SOX9 | Others | 0 | 2 | 15 | 12.5 |
| 69 | CSTF2 | Others | 0 | 0.5 | 3 | 2.5 |
| 70 | BRD7 | BAF complex | 0 | 0 | 0.5 | 0 |
| 71 | ISY1 | Spliceosome complex | 0 | 0 | 0.5 | 0 |
| 72 | NEXN-AS1 | Others | 0 | 0 | 0.5 | 0 |
| 73 | CPSF3 | Others | 0 | 0 | 0.5 | 0 |
| 74 | TMSB10 | Others | 0 | 0 | 0.5 | 0 |
| 75 | SRSF12 | Others | 0 | 0 | 0.5 | 0 |
| 76 | ANKRD62 | Others | 0 | 0 | 0.5 | 0 |
| 77 | HDGFL2 | DNA repair | 0 | 0 | 0.5 | 0 |
| 78 | VAMP2 | Others | 0 | 0 | 0.5 | 0 |
| 79 | ADIRF | Others | 0 | 0 | 0.5 | 0 |
| 80 | EPHB2 | Others | 0 | 0 | 0.5 | 0 |
| 81 | CTNNBL1 | Spliceosome complex | 0 | 0 | 0.5 | 0 |
| 82 | WBP11 | Spliceosome complex | 0 | 0 | 0.5 | 0 |
| 83 | MYD88 | Others | 0 | 0 | 0.5 | 0 |
| 84 | STXBP2 | Others | 0 | 0 | 0.5 | 0 |
| 85 | CCAR1 | Others | 0 | 0 | 0.5 | 0 |
| 86 | SUGP1 | Spliceosome complex | 0 | 0 | 0.5 | 0 |
| 87 | MESDC1 | Others | 0 | 0 | 0.5 | 0 |
| 88 | MED13 | Others | 0 | 0 | 0.5 | 0 |
| 89 | MTCH2 | Others | 0 | 0 | 0.5 | 0 |
| 90 | DHRS2 | Others | 0 | 0 | 0.5 | 0 |
| 91 | MAGED2 | Others | 0 | 0 | 0.5 | 0 |
| 92 | SATB1 | Others | 0 | 0 | 0.5 | 0 |
| 93 | ZC3H11A | Others | 0 | 0 | 0.5 | 0 |
| 94 | TMX1 | Others | 0 | 0 | 0.5 | 0 |
| 95 | PRR12 | Others | 0 | 0 | 0.5 | 0 |
| 96 | KLF16 | Others | 0 | 0 | 0.5 | 0 |
| 97 | THOC2 | Transcription regulation | 0 | 0 | 0.5 | 0 |
| 98 | NCAPD3 | heterochromatin | 0 | 0 | 0.5 | 0 |
| 99 | DUS3L | Others | 0 | 0 | 0.5 | 0 |
| 100 | SIRT1 | Histone acetylation | 0 | 0 | 0.5 | 0 |
| 101 | MED23 | Transcription regulation | 0 | 0 | 0.5 | 0 |
| 102 | ACTL6A | BAF complex | 0 | 0 | 0.5 | 0 |
| 103 | ATR | DNA repair | 0 | 0 | 0.5 | 0 |
| 104 | RAD21 | DNA repair | 0 | 0 | 0.5 | 0 |
| 105 | FAM50A | Others | 0 | 0 | 0.5 | 0 |
| 106 | HMCES | DNA repair | 0 | 0 | 0.5 | 0 |
| 107 | PC | Others | 0 | 0 | 0.5 | 0 |
| 108 | MRPL22 | Others | 0 | 0 | 0.5 | 0 |
| 109 | PPIL2 | Spliceosome complex | 0 | 0 | 0.5 | 0 |
| 110 | HEBP1 | Others | 0 | 0 | 0.5 | 0 |
| 111 | NEK5 | Others | 0 | 0 | 0.5 | 0 |
| 112 | ZC3H13 | Others | 0 | 0 | 0.5 | 0 |
| 113 | PRPF38A | Spliceosome complex | 0 | 0 | 0.5 | 0 |
| 114 | CDK5RAP3 | Others | 0 | 0 | 0.5 | 0 |
| 115 | SF3B6 | Spliceosome complex | 0 | 0 | 0.5 | 0 |
| 116 | SGPL1 | Others | 0 | 0 | 0.5 | 0 |
| 117 | PPIB | Others | 0 | 0 | 0.5 | 0 |

| | | | | | | |
|---|---|---|---|---|---|---|
| 118 | BAZ1B | BAF complex | 0 | 0 | 0.5 | 0 |
| 119 | AKAP1 | Others | 0 | 0 | 0.5 | 0 |
| 120 | NOL6 | Others | 0 | 0 | 0.5 | 0 |
| 121 | TIGAR | DNA repair | 0 | 0 | 0.5 | 0 |
| 122 | C11orf98 | Others | 0 | 0 | 0.5 | 0 |
| 123 | SCARB2 | Others | 0 | 0 | 0.5 | 0 |
| 124 | ATN1 | Others | 0 | 0 | 0.5 | 0 |
| 125 | NCOR2 | Transcription regulation | 0 | 0 | 0.5 | 0 |
| 126 | CEP290 | Others | 0 | 0 | 0.5 | 0 |
| 127 | MAST1 | Others | 0 | 0 | 0.5 | 0 |
| 128 | NDUFAF3 | Others | 0 | 0 | 0.5 | 0 |
| 129 | PFKFB3 | Others | 0 | 0 | 0.5 | 0 |
| 130 | SENP3 | Others | 0 | 0 | 0.5 | 0 |
| 131 | TLE1 | Transcription regulation | 0 | 0 | 0.5 | 0 |
| 132 | UBA2 | Others | 0 | 0 | 0.5 | 0 |
| 133 | UHRF2 | heterochromatin | 0 | 0 | 0.5 | 0 |
| 134 | PAPSS1 | Others | 0 | 0 | 0.5 | 0 |
| 135 | FAM76B | Others | 0 | 0 | 0.5 | 0 |
| 136 | STX7 | Others | 0 | 0 | 0.5 | 0 |
| 137 | TOMM6 | Others | 0 | 0 | 0.5 | 0 |
| 138 | EIF3D | Others | 0 | 0 | 0.5 | 0 |
| 139 | PLOD3 | Others | 0 | 0 | 0.5 | 0 |
| 140 | SHMT1 | Others | 0 | 0 | 0.5 | 0 |
| 141 | ECHDC1 | Others | 0 | 0 | 0.5 | 0 |
| 142 | GPATCH8 | Others | 0 | 0 | 0.5 | 0 |
| 143 | P4HA1 | Others | 0 | 0 | 0.5 | 0 |
| 144 | HOXD10 | Others | 0 | 0 | 0.5 | 0 |
| 145 | KIF23 | Others | 0 | 0 | 0.5 | 0 |
| 146 | RNF213 | Others | 0 | 0 | 0.5 | 0 |
| 147 | UBR7 | Others | 0 | 0 | 0.5 | 0 |

### List of Antibodies

| Antibody | Vendor | Catalog# | Used For |
|---|---|---|---|
| SOX9 | CST | #82630 | IP, WB |
| Con IgG | Epicypher | #13-0042 | CnR |
| BRD7 | Invitrogen | #PA5-100009 | CnR, WB |
| BRD9 | Invitrogen | #PA5-113488 | CnR, WB |
| DPF2 | Invitrogen | #PA5-21079 | CnR, WB |
| PBRM1 | Thermo Fischer Scientific | #A700-019 | CnR, WB |
| SOX9 | Millipore | AB5535 | CnR, WB, IP |
| ARID1A | CST | #12354 | CnR, WB |
| ARID2 | CST | #MA5-27862 | CnR, WB |
| V5 | abcam | ab9116 | CnR, IP |
| V5 | ProteinTech | v5ta | IP |
| Anti Rabbit Normal IgG | CST | 2729 | IP |
| PROM1 / CD133 | CST | 86721 | WB |
| SMARCB1 | CST | 91735 | IP, WB, CnR |
| SMARCC1 | CST | 11956 | IP, WB, CnR |
| SMARCD1 | CST | 35070 | WB |
| SMARCE1 | CST | 33360 | WB |

### List of Oligos used for qRT-PCR

| Species | Gene | Forward | Reverse |
|---|---|---|---|
| Human | PROM1 | AACCTACAGCATATTCTTCA | AACGAACAGCATTTCTCTCTCAAGA |
| Human | GAPDH | TGTTGCCATCAATGACCCCTT | CTCCACGACGTACTCAGCG |
